# Executable Network of SARS-CoV-2-Host Interaction Predicts Drug Combination Treatments

**DOI:** 10.1101/2021.07.27.453973

**Authors:** Rowan Howell, Matthew A. Clarke, Ann-Kathrin Reuschl, Tianyi Chen, Sean Abbott-Imboden, Mervyn Singer, David M. Lowe, Clare L. Bennett, Benjamin Chain, Clare Jolly, Jasmin Fisher

**Author notes:** Correspondence should be addressed to Jasmin Fisher or Clare Jolly. These authors contributed equally to this work (Rowan Howell, Matthew A. Clarke, Ann-Kathrin Reuschl).

## Abstract

The COVID-19 pandemic has pushed healthcare systems globally to a breaking point. The urgent need for effective and affordable COVID-19 treatments calls for repurposing combinations of approved drugs. The challenge is to identify which combinations are likely to be most effective and at what stages of the disease. Here, we present the first disease-stage executable signalling network model of SARS-CoV-2-host interactions used to predict effective repurposed drug combinations for treating early- and late-stage severe disease. Using our executable model, we performed *in silico* screening of 9870 pairs of 140 potential targets and have identified 12 new drug combinations. Camostat and Apilimod were predicted to be the most promising combination in effectively supressing viral replication in the early stages of severe disease and were validated experimentally in human Caco-2 cells. Our study further demonstrates the power of executable mechanistic modelling to enable rapid pre-clinical evaluation of combination therapies tailored to disease progression. It also presents a novel resource and expandable model system that can respond to further needs in the pandemic.

## Introduction

COVID-19 is a complex disease in both dynamics and severity^1–4^. While most infections only result in mild symptomatic presentation, a proportion suffer more severe disease, for example, the CDC estimate 4.9% of SARS-CoV-2 infections in the USA resulted in hospitalisation in the period up to March 2021^5,6^. Mild cases see an innate immune and interferon (IFN-I/III) response to SARS-CoV-2 infection, as typically observed in other viral infections, likely supporting clearance of the virus and the onset of adaptive immunity^1,7,8^. By contrast, a delayed or absent IFN I/III response to infection may contribute to presentation of severe disease^7,9–14^. This is especially of concern as emerging variants have adapted to allow potent host immune antagonism^15^, allowing the virus to replicate with little opposition in the early stages of infection^1,10^. This can then be followed by a maladaptively strong and persistent inflammatory response only after the majority of viral replication has occurred^9,16^, leading to life threatening conditions such as acute respiratory distress syndrome (ARDS)^17^.

There is an urgent need for new effective drugs to manage COVID-19 infection to lower morbidity, mortality and reduce the strain on healthcare systems^18,19^. While several vaccines have been approved^20–22^, many millions more patients will need treatment during the years it will take to deliver vaccination worldwide^23,24^. The path for de novo drug discovery is long and complex; the best hope for rapid development of new therapies is therefore to combine readily available drugs^25^, with a focus on affordable and well-tested treatments, such as Dexamethasone, which will be necessary to treat COVID-19 in low-income countries that are struggling to obtain sufficient doses of vaccines. Given the large range of potentially suitable compounds, several challenges arise: which drugs are most effective and at what stage of the infection? Are there combinations of drugs that allow effective treatment at lower doses and reduced toxicity?

Driven by the different characteristics of early and late stages of severe COVID-19, and the need to find therapies appropriate to each stage of the disease without interfering with the effective immune response in mild cases, we have developed a computational model that can reproduce these different phases of COVID-19. This approach builds upon our previous success at finding novel drug targets and effective drug combinations for cancer therapy^26–29^. We created a detailed network map of the interaction between SARS-CoV-2 and lung epithelial cells, as pulmonary involvement in particular is a hallmark of severe disease. We use this model to screen thousands of drug combinations to find treatments that block either key viral-host interactions important to viral replication, or the pathogenic dysregulation of the immune response. We focus especially on host-directed therapies, which may be more robust to future variants of the virus^30^. We identify several combinations of re-purposed drugs that are predicted to act together to target viral replication in the early stages of the disease or inflammation in the late stage. Moreover, we propose that the model and screen results represent a powerful resource for the community to generate hypotheses about the potential effects of targeting host and viral targets in combination.

## Results

### Executable network map of the interaction between SARS-CoV-2 and lung epithelium

We model the interaction map between SARS-CoV-2 and lung epithelial cells as an executable Qualitative Network^31^ model (Figure 1) using the BioModelAnalyzer (BMA) tool (http://www.biomodelanalyzer.org). This model is an executable computer program in graphical form characterising the signalling regulatory network (Figure 2A), made up of 175 nodes, representing viral and host proteins, as well as cellular processes such as viral replication and inflammation. There are 387 edges joining these nodes representing activating or inhibiting interactions (Supplementary Table S1). At any point, each node has a discrete level that represents its activity that is determined by the input of activations and inhibitions that the node receives and the node’s target function (Methods, Supplementary Table S2).

**Figure 1.**
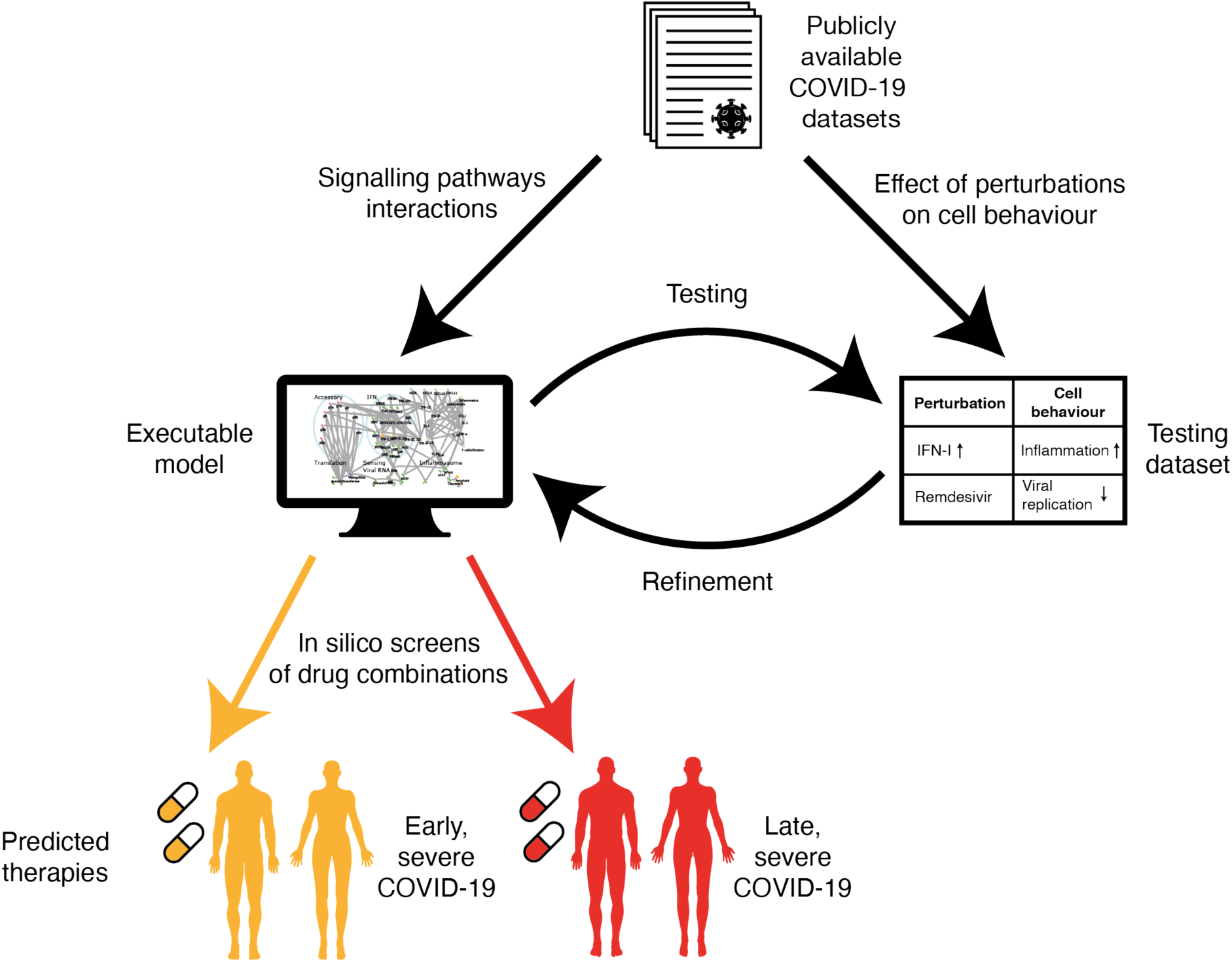
Schematic workflow of SARS-CoV-2 infection modelling. Publicly available COVID-19 datasets are collated into two subsets. Data on the point-to-point signalling pathway interactions of the virus with the host cell, and of the host cell in response to infection, are used to build a network model. Data from experiments showing how the overall behaviour of the infected cell changes under perturbations, such as a potential treatment, are used as a testing dataset to validate the model. The readouts of the model are compared to the testing dataset and the model is refined iteratively until it reproduces all the experimentally observed behaviours. We then screen the effects of potential drug treatments, either singly or in combination, on the model, to find the best predicted therapies for early and late stage severe COVID-19.

**Figure 2.**
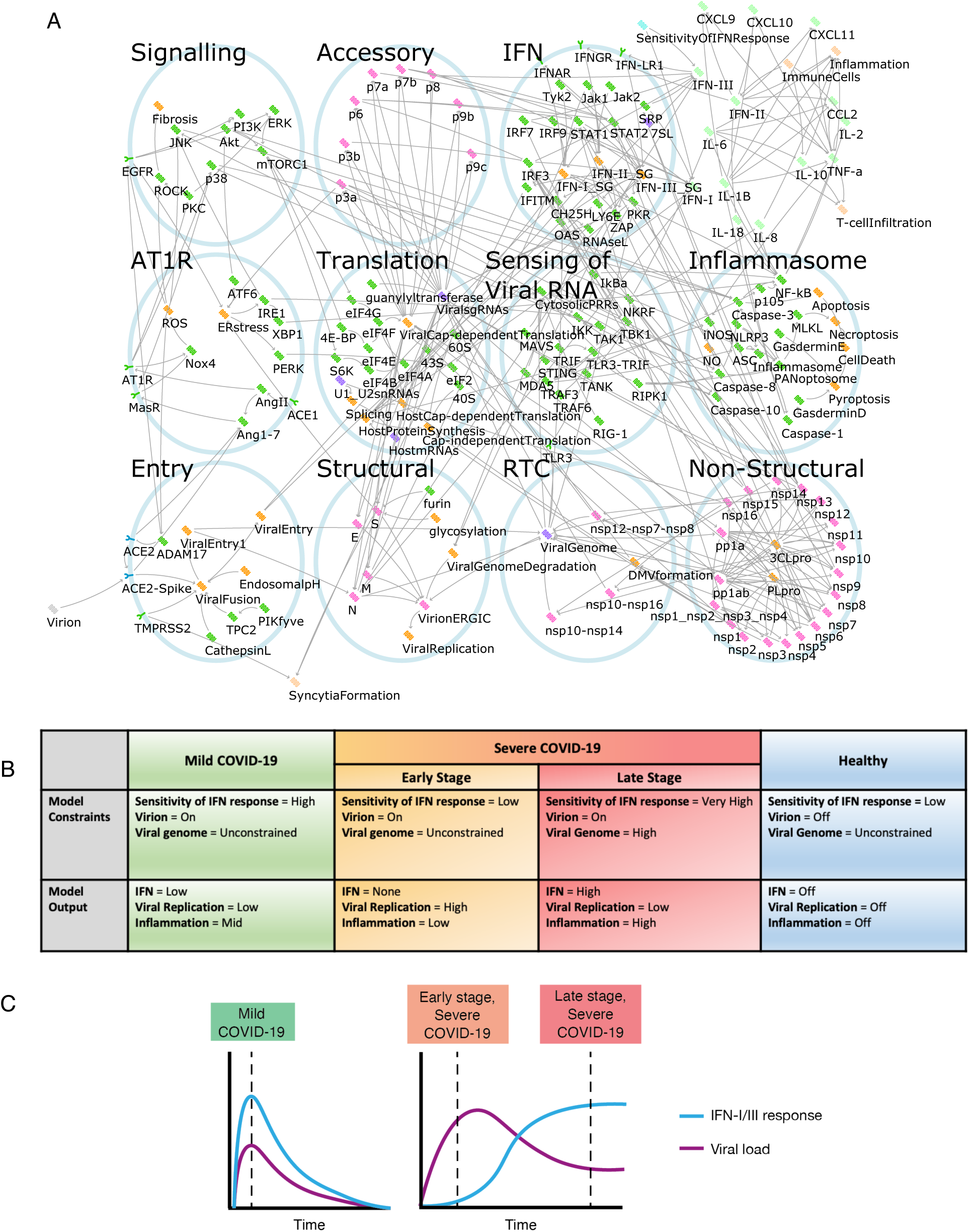
Computational modelling of host-virus interaction in COVID-19. (A) The SARS-CoV-2 infection signalling network as modelled in the BioModelAnalyzer tool. Viral proteins depicted in pink, host proteins depicted in green, cellular processes depicted in orange, RNAs depicted in purple. Activating interactions are represented by arrows (→), inactivating by bars (⊣). (B) Description of the model constraints and outputs for the three states of the disease and the control healthy state. (C) Trajectories of IFN –I/III response and viral load in mild and severe COVID-19 (as described in Park and Iwasaki, *Cell Host and Microbe*, 2020) used to inform the model.

Given the importance of the type I/III IFN response, we use it in our modelling to reproduce the mild case and both the early and late stages of severe cases of COVID-19, as well as an uninfected control, through setting key input nodes in the model (see Figure 2B, Supplementary Table S3). For example, at the presentation stage of severe COVID-19 the IFN-I/III response is evaded and suppressed^1,7^; we have termed this the “*Sensitivity of IFN Response*” to convey both the response to, and magnitude of, IFN-I/III production, and set this node in the model to a low level. Conversely, in the late stage of severe COVID-19, IFN-I/III response is increased at the site of infection^32^, and most viral reproduction has already occurred, leaving behind viral RNA and proteins that trigger further maladaptive immune responses. We reproduce this by setting the *Sensitivity of IFN Response* node to a high level and specify a high burden of viral RNA and protein by the *Viral Genome* node, assuming replication has peaked (Figure 2B, C, Supplementary Table S3). Finally, our model also represents mild disease and uninfected individuals to ensure that any potential new treatment does not have adverse effects in these cases.

### Network model validation by comparison to known effects of SARS-CoV-2 infection and treatment

Having generated a network model of viral infection, we first verified that it reproduced key results from experiments that have defined SARS-CoV-2 interaction with lung epithelial cells, as well as known clinical results establishing the utility of different treatments at different stages of the disease (Supplementary Table S4). In particular, we tested the model responses to monotherapies that have been used as COVID-19 treatments, such as the antivirals Remdesivir^33^ and Lopinavir^34^; and the immune modulators Tocilizumab^35^, Dexamethasone^36^, Interferon α-2a (Roferon-A)^37^ and Ruxolitinib^38^. Drugs were modelled by setting the affected node in the signalling network to either zero, to represent an inhibitory drug, or to the maximum value, to represent an agonist. The effect of the treatment was then assessed by measuring the level of the nodes representing cellular processes such as viral replication and inflammation (Figure 3).

**Figure 3.**
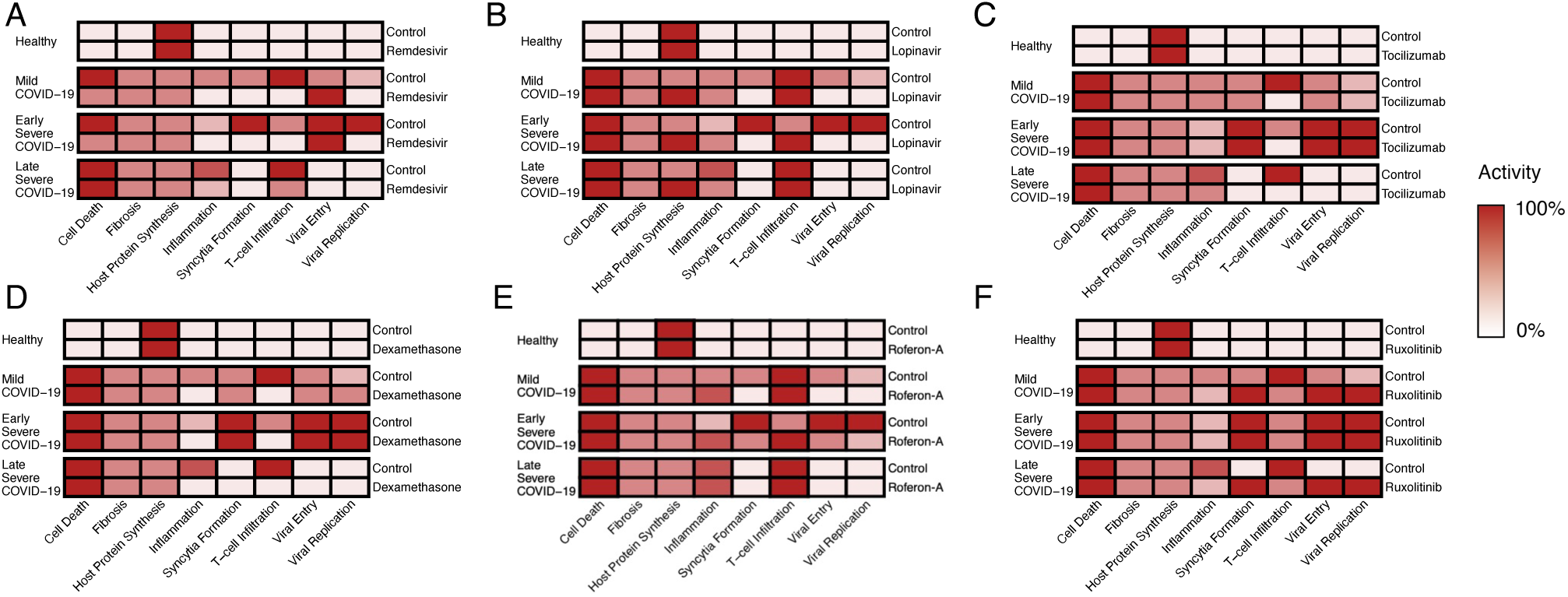
Model output reproducing the effect of known drug treatments for COVID-19. The behaviour of (A) Remdesivir, (B) Lopinavir, (C) Tocilizumab, (D) Dexamethasone, (E) Roferon-A and (F) Ruxolitinib. Colour strength represents strength of symptom predicted by the model following drug intervention. All nodes normalised to maximal level of respective nodes, and range between 0-100%.

### Inhibiting protein translation predicted to suppresses viral replication in early stages of severe COVID-19

Our approach to find new and improved treatments is to test the response of our computational model to an exhaustive search across combinations of many different drugs, either clinically approved or in phase II/III trials, to determine optimal treatments for each stage of development of the disease (Table 1). We implemented the screening by fixing the activity of nodes in the network to a constant value to represent the effect of mono- or combination treatments over all drug combinations and disease stages (Figure 2B, Supplementary Table S3, S5). This *in silico* screen predicted the most effective drug treatments against each stage of COVID-19 and ensured no adverse effects in mild cases of the disease.

**Table 1.** List of approved drugs used for *in silico* screening. The network model was used to evaluate the best single and combinations of 38 drugs, either clinically approved or in phase II/III trials.

| Drug | Pathway | Gene/Protein Affected | References |
| --- | --- | --- | --- |
| 4-PBA | AT1R | ERstress | Hsu, A. C.-Y., 2020 |
| ACEi | AT1R | ACE1 | Vaduganathan, M., 2020 |
| Actimmune | IFN | IFN-II | Razaghi, A., 2016 |
| Aliskiren | AT1R | AngII, Ang1-7 | Sriram, K., 2020 |
| Anifrolumab | IFN | IFNAR | Riggs, J. M., 2018 |
| Apilimod | Entry | PIKfyve | Ou, X., 2020 |
| ARB | AT1R | AT1R | Vaduganathan, M., 2020 |
| Baricitinib | IFN | Jak1, Jak2, ViralFusion | Richardson, P., 2020 |
| Basiliximab | Cytokines & Chemokines | IL-2 | Onrust, S et al. (1999) |
| Belnacasan | Inflammasome | Caspase-1 | Kudelova, J., 2015 |
| Camostat | Entry | TMPRSS2 | Hoffmann, M., 2020 |
| Canakinumab | Cytokines & Chemokines | IL-1B | Ucciferri, C., 2020 |
| Carlumab | Cytokines & Chemokines | CCL2 | Lim, S. Y., 2016 |
| Cetuximab | Signalling | EGFR | Drugbank 2020 |
| Chloroquine | Structural | EndosomalpH, glycosylation, TNF-a, IL-6 | Wang, M., 2020; Savarino, A., 2003 |
| Colchicine | Inflammasome | Inflammasome | Martinez, G. J., 2017 |
| Copanlisib | Signalling | PI3K | Yang, J., 2019 |
| Dexamethasone | Inflammasome | IL-10, TNF-a, NF-kB, JNK, ERK, p38, IL-1B, IL-8 | Selvaraj, V., 2020; Smoak, K. A., 2004 |
| Disulfiram | Inflammasome | GasderminD | Hu, J. J., 2020 |
| Emricasan | Inflammasome | Caspase-1, Caspase-3, Caspase-8, Caspase-10 | Kudelova, J., 2015 |
| Fedratinib | IFN | Jak2 | Szelag, M., 2016 |
| Homoharringtonine | Translation | 40S | Choy, K.-T., 2020 |
| Infliximab | Cytokines & Chemokines | TNF-a | Robinson, P. C., 2020 |
| Lopinavir | Non-structural | 3CLpro | Choy, K.-T., 2020; Ma, C., 2020 |
| Losmapimod | Signalling | p38 | Drugbank 2020 |
| Miltefosine | Signalling | Akt | Nitulescu, G. M., 2015 |
| N-acetylcysteine | AT1R | ROS | Meng, Y., 2015 |
| Nivocasan | Inflammasome | Caspase-1, Caspase-8 | Kudelova, J., 2015 |
| Rapamycin | Signalling | mTORC1 | Ballou, L. M., 2008 |
| Recombinant IL-10 | Cytokines & Chemokines | IL-10 | Asadullah, K., 2003 |
| Remdesivir | Non-structural | nsp12-nsp7-nsp8 | Wang, M., 2020 |
| Roferon-A | IFN | IFN-I | Mantlo, E., 2020 |
| Ruxolitinib | IFN | Jak1, Jak2 | Szelag, M., 2016 |
| SAR113945 | Sensing of Viral RNA | IKK | Herrington, F. D., 2015 |
| Tetrandrine | Entry | TPC2 | Ou, X., 2020 |
| Tocilizumab | Cytokines & Chemokines | IL-6 | Mihara, M., 2011 |
| Ulixertinib | Signalling | ERK | Hsu, A. C.-Y., 2020 |
| Umifenovir | Entry | ACE2-Spike, S | Wang, X., 2020; Kim, S. Y., 2020 |

At the presentation stage of severe COVID-19, characterised by a weak IFN-I/III response (Figure 2B, Supplementary Table S3), treatments that are able to arrest viral replication before it can trigger an excessive inflammatory response are needed. As this stage occurs mainly prior to hospitalisation, these treatments must be taken as an anticipatory therapy; at or close to the time of suspected exposure or infection. It is therefore critical that they do not adversely affect mild COVID-19 patients, as they will likely be administered before it is possible to stratify patients by prognosis. Our model predicts that, for example, consistent with known treatments in clinical use and trials, Remdesivir^33^ and recombinant Interferon *α*-2a (Roferon-A)^37^ are effective at reducing viral replication at the early stage of severe disease (Figure 4A). In addition, several host translation inhibitors drawn from anti-cancer therapies were also shown to be effective by our model, including Homoharringtonine or Rapamycin (Figure 4A). Homoharringtonine has previously been identified in a profile of potential drugs targeting SARS-CoV-2^39^, believed to act through inhibition of the eukaryotic small ribosomal subunit (40S)^40^, suppressing viral replication as the virus depends on host translation machinery for production of viral proteins^41^. Homoharringtonine has cleared Phase II trials for chronic myeloid leukaemia^40^ and so is a good candidate to be fast tracked for use. Similarly, Rapamycin and other mTOR inhibitors, which block translation, are in trials as post-exposure prophylaxis for COVID-19, to prevent the early stage of severe disease from progressing^42,43^. Our model predicts that Rapamycin is only effective against cap-dependent rather than cap-independent protein synthesis. However, as SARS-CoV-2 is dependent upon hijacking the cap-dependent form of protein synthesis^44^ this allows equal effectiveness in suppressing viral replication as with other translation inhibitors, with less effect on host protein synthesis and possibly fewer side effects (Figure 5I). Other translation inhibitors drawn from anti-cancer therapies further demonstrate the potential of host-directed treatments^45,46^.

**Figure 4.**
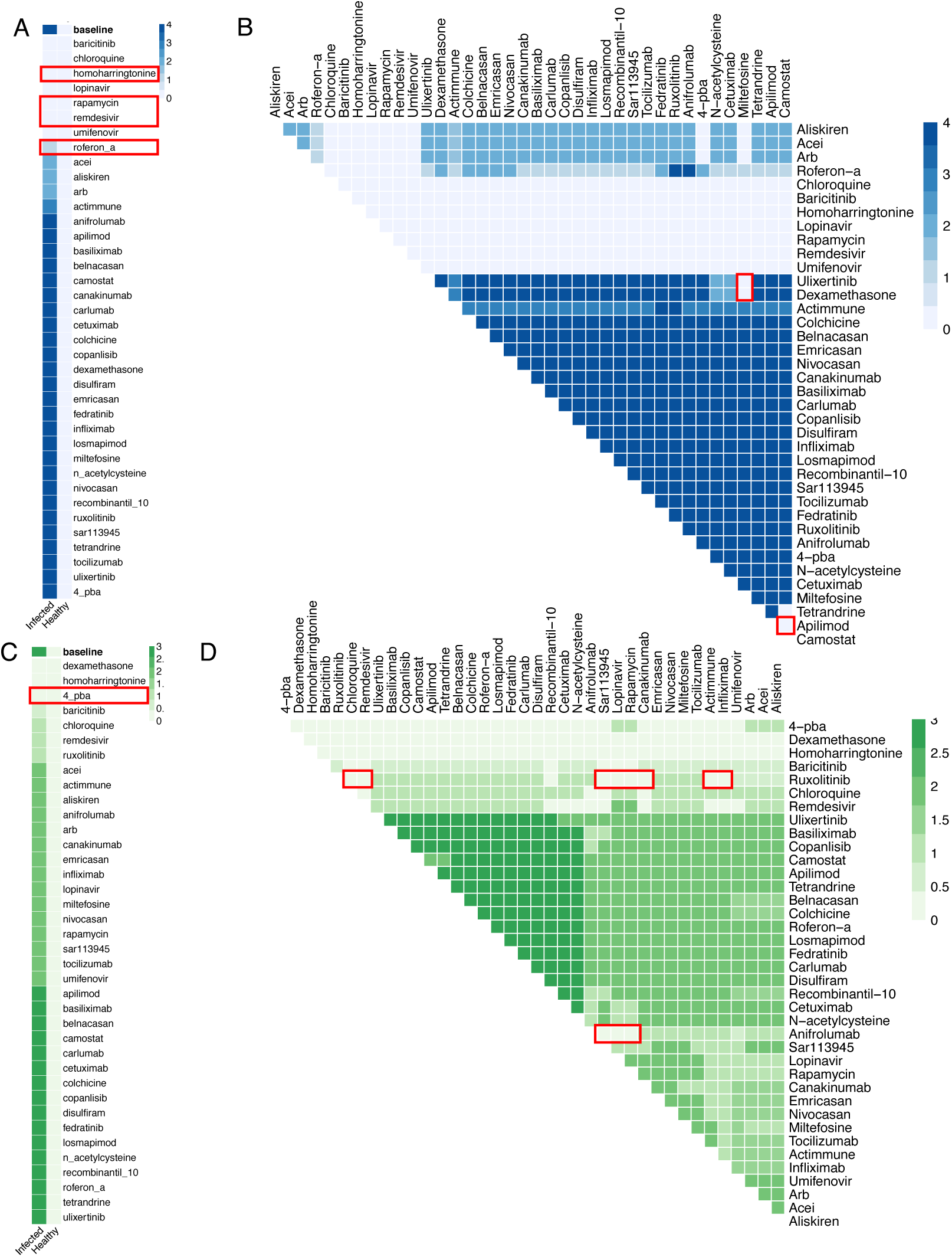
*In silico* screens for optimised drug combinations to treat COVID-19. (A) Effect of single drugs (rows) on viral replication in infected cells in early stage of severe disease and in control healthy cells. (B) The effects of drug combinations on viral replication in early stage of severe COVID-19. (C) The effect of single drugs (rows) on inflammation in infected cells in late stage of severe disease and in control healthy cells. (D) Effect of drug combinations on inflammation in late stage of severe COVID-19. Colour of cell corresponds to level of viral replication (blue) and inflammation (green). Red boxes indicate therapies discussed in detail.

**Figure 5.**
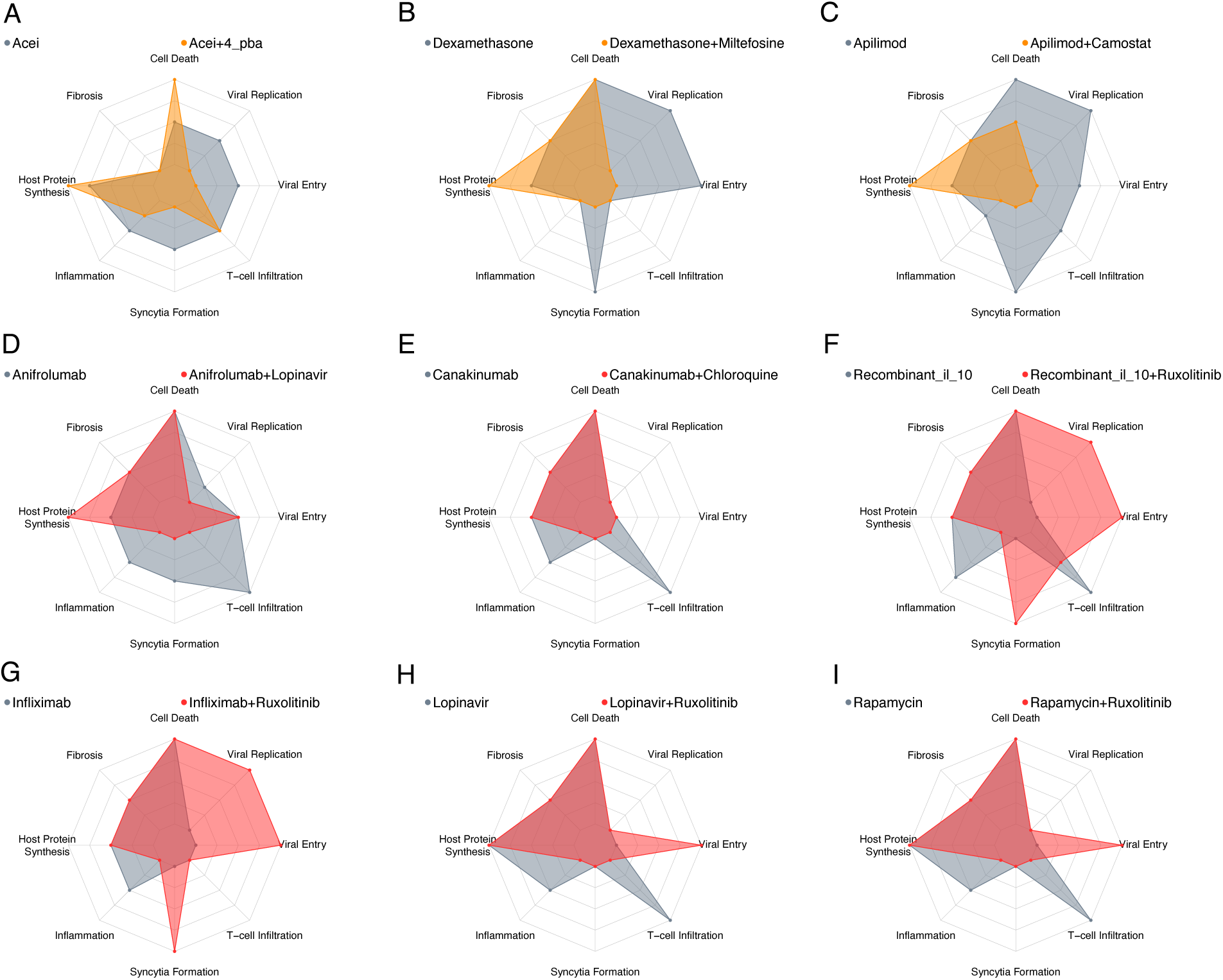
Predicted effective drug treatments for severe COVID-19. (A-C) The effect of monotherapy vs drug combinations identified to reduce viral replication in early stage of severe COVID-19. Monotherapy (grey), combination (orange), with the strength of the biological process denoted by radial distance. (D-I) The effect of monotherapy vs drug combinations identified to reduce inflammation in late stage of severe COVID-19. Monotherapy (grey), combination (red). All nodes normalised to maximal level of respective nodes, and range between 0-100%.

### Combination of Camostat and Apilimod suppress viral replication in early severe COVID-19

Our analysis further reveals options to combine different treatments, focusing on cases where there is an improvement in other pathological processes, in addition to decreasing viral replication. Viral entry inhibitors Camostat (TMPRSS2 inhibitor) and Apilimod (PIKfyve inhibitor) were shown to reduce viral entry but not viral replication in the model, whereas in combination they showed additional ability to prevent viral replication (Figure 4A-B). This combination also showed improved effects on other key aspects of pathology compared to monotherapy (the effect of mono- vs combination therapy is shown in Figure 5C for all simulated symptoms), recapitulating the effects of Apilimod seen in TMPRSS2-negative cell lines^47^. Miltefosine (AKT inhibitor) was shown in the model to be ineffective against viral replication alone but exhibits several promising combinations. A combination of Miltefosine with Ulixertinib (MAPK inhibitor) predicted effects across the redundant pathways controlling translation, leading to similar efficacy as Rapamycin (Figure 4A-B). However, increased inflammation is also predicted as a side effect (Figure S1F). Conversely, a combination of Miltefosine with the approved anti-inflammatory Dexamethasone was predicted by the model to target both viral replication and inflammation (Fig. 4B, 5B), if the ERK-inhibitory effects of Dexamethasone are sufficient to replace a dedicated inhibitor like Ulixertinib^48^.

### Reduction of ER-stress using 4-PBA predicted to reduces excessive inflammation in late stage severe COVID-19

We next considered the late stage of the severe disease, as defined by an inappropriate inflammatory response that must be curbed (Figure 2B, Supplementary Table S3). These treatments must be used with care, lest they reduce the useful anti-viral immune response in the early severe or mild cases. As seen in the landmark RECOVERY trial^36^, Dexamethasone alone is a partially effective anti-inflammatory treatment for this stage of COVID-19 infection (Figure 4C). Our model also predicts that sodium phenylbutyrate (4-PBA) may have some efficacy at this stage. Endoplasmic reticulum (ER) stress is dysregulated by SARS-CoV spike protein (S)^49,50^. It has been suggested that this pathway plays a pro-inflammatory role, via activation of NFκB, in coronavirus infections in general^51,52^; preliminary results show similar effects in SARS-CoV-2^53,54^. 4-PBA is a chemical chaperone that helps reduce protein misfolding and so reduces ER-stress (Figure 4C). 4-PBA is already FDA approved for urea cycle disorders, and is under consideration for use in cystic fibrosis, glioma and acute myeloid leukaemia^55,56^. However, this therapy is predicted by the model to slightly worsen viral replication in the mild case by decreasing IFN-II activity, and anti-viral immunity (Supplementary Figure S2H). While this change is small, it may prevent this therapy being used prophylactically, but it can be safely applied in the late stage of severe disease, once viral proliferation has peaked, and viral load is waning. Similarly, the IFN-I blocking agent Anifrolumab decreases inflammation in the late stage of severe disease, but this has counterproductive effect on viral replication in mild disease (Supplementary Figure S2H).

### Anti-viral Lopinavir predicted to suppress inflammation when used in combination with Ruxolitinib in late stage severe COVID-19

Ruxolitinib is already predicted to be effective as a monotherapy and is currently being trialled for COVID-19^38^. However, Ruxolitinib alone may, through inhibition of JAK-STAT and IFN-I signalling, impair innate immune-mediated suppression of viral entry and replication in mild cases (Figure 3F, S2G, H) and has no benefit in the early severe disease (Fig 3F, S3D). While our model does not suggest combination treatments that can ameliorate this, meaning that Ruxolitinib is likely too risky to use in mild disease, we predict several ways to enhance the effects of Ruxolitinib appropriate for the late stage of severe disease. These include drugs that are ineffective alone such as SAR113945 (IKK inhibitor), Infliximab (TNF-*α* inhibitor), Canukinumab (IL-1B inhibitor) and recombinant IL-10 (Figure 4C, D).

The trade-off of these combinations suggests that they should be used only in the late stage of the disease. For example, a combination of Rapamycin and Ruxolitinib is predicted to be more effective against inflammation than either drug alone. However, together these drugs are also predicted to increase viral entry (Figure 5I). This may suggest that Rapamycin alone could be a good early monotherapy and, if the disease progresses, could be combined with Ruxolitinib in the late stage when viral entry has already peaked. However, secondary infection, especially bacterial, could be a concern given the potent immunosuppressive effects of some of these combinations.

Lopinavir, often administered with Ritonavir, an agent that increases the half-life of Lopinavir in plasma, acts against the virus directly by blocking viral proteases, but has not demonstrated success in treating hospitalised COVID-19 patients^34,57^. This is consistent with our model’s prediction, which suggests that while Lopinavir may be effective at reducing viral replication in the early stages of severe COVID-19 (Figure 4A), it is not predicted to provide additional benefit in the late stage of the disease (Figure S4G, H). However, while Lopinavir inhibits the 3CL protease of SARS-CoV-1^59^, it may only be effective against SARS-CoV-2 at toxic doses^39^ or not at all^60^ though dedicated 3CLPro inhibitors are in development^61^. Lopinavir is one of the therapies currently being tested in the FLARE trial^58^, which may shed light on this.

Promisingly, targeting 3CL protease in our network model showed interesting anti-inflammatory effects, especially when used in combination with other drugs. 3CL protease is responsible for the proper processing of many viral proteins, including those involved in RNA-dependent RNA polymerase activity (RDRP). Without RDRP activity, the virus is unable to produce sgRNAs required for translation of structural and accessory proteins, including p3a and S, which have been shown to have a pro-inflammatory effect^53,54^. Direct targets of 3CLpro activity like viral nsp10 have also been shown to modulate expression of pro-inflammatory cytokines^62^. This means that 3CL protease inhibition could potentially increase the effectiveness of anti-inflammatories such as Anifrolumab (IFNAR inhibitor) or Ruxolitinib (JAK1/2 inhibitor) (Figure 4C-D). As such, anti-3CLpro drugs may provide benefit even after viral replication has peaked by also playing a role in a synergistic combination with other drugs to control dysregulated inflammation, and so can be beneficial in both phases of the disease. This contrasts with drugs such as Remdesivir, which blocks viral replication effectively by blocking RDRP, but leaves other pro-inflammatory proteins intact. Consequently, our model predicts that Remdesivir will only be effective when applied early in the course of the disease. This is in line with clinical trials showing only modest effects when applied in patients who are already hospitalised^33^.

### Characteristic differences in cytokine level distinguish mild, early severe and late severe COVID-19

As our model predictions show that the effect of drugs is dependent on the stage of the disease at which they are administered, we further searched using our network model for potential prognostic biomarkers for the different stages by comparing the steady state levels of each node in mild, early and late severe COVID-19. We observed that a characteristic signature of the early disease is lower activity of cytokines, such as TNF-α, IL-6 and IL-10 and Interferons type I-III, compared to the mild case (Supplementary Figure S5). These will subsequently rise in the late stage of the severe case, with some such as IL-6 exceeding the level seen in the mild case. This is in line with the evidence that severe disease arises, in part, due to an insufficient initial innate immune response, followed by a maladaptively strong and persistent inflammatory response and the prior identification of TNF-*α*, IL-6, IL-8, IL-10 and CXCL10 as prognostic markers for COVID-19 disease severity in hospitalised patients^63–67^. This suggests that measuring cytokine levels could help distinguish mild from severe cases in the early stages of disease, but we also note the risk of an incorrect prognosis due to similarities between late severe and mild cases.

### Camostat and Apilimod combined have significantly greater effect suppressing viral entry and replication

To validate our predictions, we selected the most promising combination treatments affecting viral replication identified from modelling severe, early disease. Our model predicts that Camostat and Apilimod would have an enhanced effect on viral replication in combination compared to monotherapy (Supplementary Table S7). We tested this in Caco-2 cells and found that, as has been previously reported^68^, Camostat is effective at limiting viral entry leading to a reduction in SARS-CoV-2 nucleocapsid protein-positive cells in culture, but cannot completely suppress viral entry and replication (Figure 6A). Addition of Apilimod significantly reduced SARS-CoV-2 infection further, even at the maximum dose of Camostat (Figure 6B), in an additive manner (Supplementary Figure S6). We further investigated whether the ERK inhibitory effects of Dexamethasone^48^ were sufficient to synergise with Akt inhibition by Miltefosine (Supplementary Figure S7), as this may provide extra benefit to Dexamethasone application early in disease progression. This builds on earlier findings that the combination of MAPK and PI3K pathway inhibitors is effective for MERS-CoV^69^, and inhibitors of the PI3K/Akt/mTOR pathway are effective against SARS-CoV-2^70,71^. However, this combination was ineffective except at doses of Miltefosine that showed toxicity, potentially due to limited ERK inhibition by Dexamethasone in Caco-2 cells (Supplementary Figure S7).

**Figure 6.**
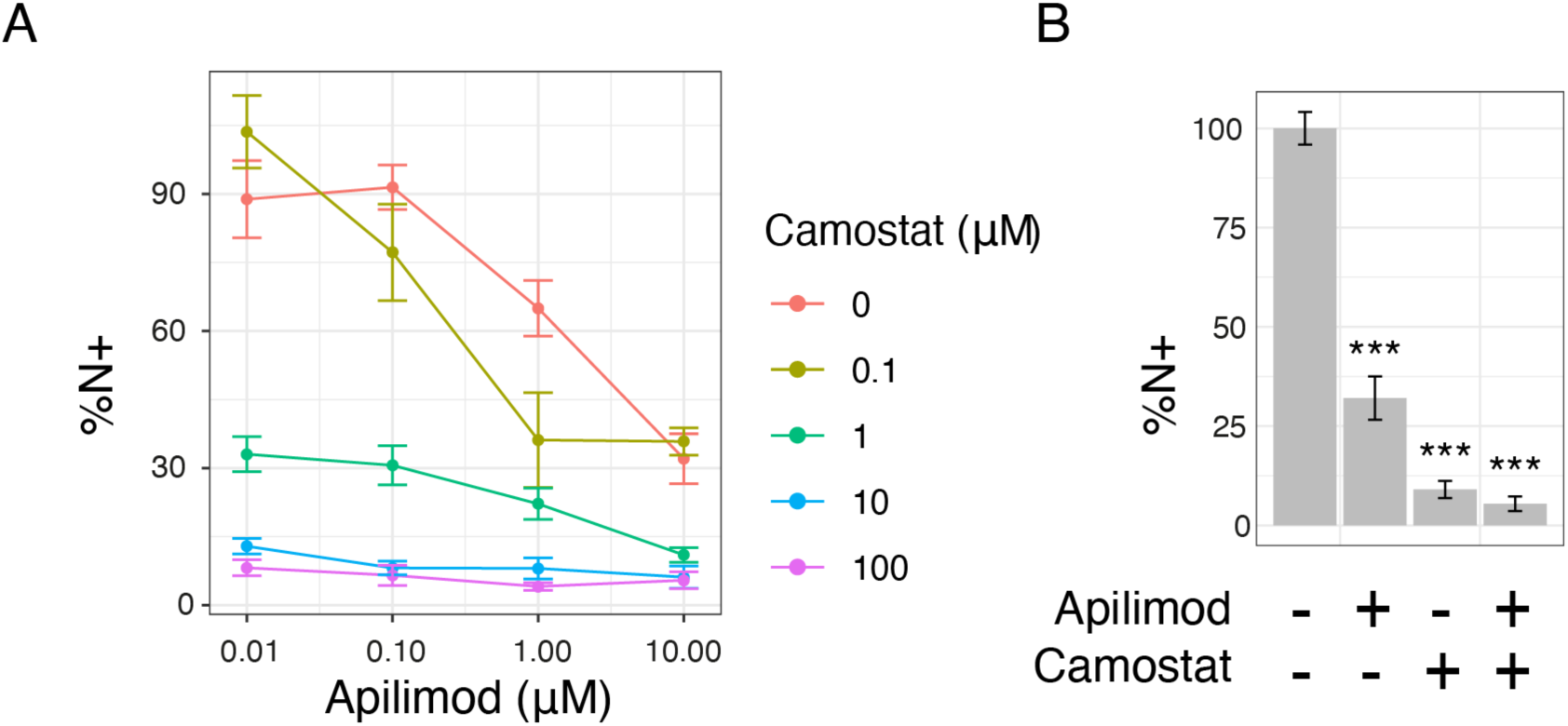
Combination of Camostat and Apilimod suppresses viral entry and replication more effectively than monotherapy. (A) Percentage cells expressing nucleocapsid protein relative to control for Apilimod and Camostat combinations. (B) Percentage cells expressing nucleocapsid protein for control, Apilimod (10µM), Camostat (100µM) or a combination of Apilimod (10µM) and Camostat (100µM). Data show the mean and standard error, from three independent experiments. *** indicates p<0.001 by two-way ANOVA.

## Discussion

Using our computational model, we can predict differential responses to therapy at different disease stages and explore the potential benefits of combination therapies over monotherapies, for treating COVID-19 patients to identify the most effective combinations (Table 2). In this study we have focussed on screening drugs that are clinically approved or in Phase II/III trials as these have the potential to immediately benefit patients in the current crisis and have had their efficacy validated in clinical practice. We further assume all drugs are maximally effective against all their putative targets to find the broadest array of potential therapies to guide experiments. A specific advantage of our computational approach is that we are able to screen a large number of potential drugs and combinations of drugs in a few hours. We have additionally screened for effective mono- and combination treatments for 37 more drugs of interest that are not yet approved or under trial (Supplementary Table S5, Supplementary Figure S8-9). In addition, we have tested all possible hypothetical interventions against all the viral and host proteins in the network model – screening a total of over 9,000 combinations. These targets, if they can be made druggable, suggest potential novel viral or host-directed targets (Supplementary Figure S10-11). Together, the network model and the drug screening algorithm provide a valuable resource that can be leveraged by the scientific community to generate hypotheses about the effect of potential therapies on cellular processes and complement existing screens of monotherapies^72^.

**Table 2.** Summary of predicted effective treatments for early and late severe COVID-19. Drug treatments predicted by the computational model to be effective in early (orange) or late (red) stage severe COVID-19.

|  | Predicted treatments to reduce viral replication in early-stage severe COVID-19 | Predicted treatments to reduce inflammation in advanced-stage severe COVID-19 |
| --- | --- | --- |
| Single drugs | • Umifenovir<br>• Baricitinib<br>• Lopinavir<br>• Rapamycin<br>• Remdesivir<br>• Roferon-A | • Dexamethasone (anti-inflammatory)<br>• 4-PBA (ER stress inhibitor) |
| Drug combinations | • Camostat (TMPRSS2 inhibitor) + Apilimod (PIKfyve inhibitor)<br>• ACE1 inhibitor + 4-PBA (ER stress inhibitor)<br>• ACE1 inhibitor + Miltefosine (Akt inhibitor)<br>• Miltefosine (Akt inhibitor) + Ulixertinib (MAPK inhibitor) | • Ruxolitinib (Jak inhibitor) + Recombinant IL-10<br>• Ruxolitinib (Jak inhibitor) + Canakinumab (IL-1B inhibitor)<br>• Ruxolitinib (Jak inhibitor) + Infliximab (TNF-α inhibitor)<br>• Ruxolitinib (Jak inhibitor) + SAR113945 (IKK inhibitor)<br>• Ruxolitinib (Jak inhibitor) + Rapamycin (mTOR inhibitor)<br>• Ruxolitinib (Jak inhibitor) + Lopinavir (3CLpro inhibitor)<br>• Anifrolumab (IFNAR inhibitor) + Lopinavir (3CLpro inhibitor)<br>• Anifrolumab (IFNAR inhibitor) + SAR113945 (IKK inhibitor) |

There are several computational approaches that have been applied to drug screening that are being applied to COVID-19, each with their own strengths and weaknesses. For example, as the structures of SARS-CoV-2 proteins have been determined, existing databases of small molecules have been leveraged to survey thousands of candidates to bind to these proteins^73^. Such studies helped identify 3CLpro as a common target, as well as use of drugs such as Lopinavir and Remdesivir^25^. Another approach is the use of protein-protein interaction networks, which are particularly suited to identifying host-directed therapies^74,75^. These networks can be analysed for their structure alone and can cover a broad range of potential targets. These have been used to identify, for example, the potential for cancer drug repurposing^45,46^. However, static-network analyses rely on the assumption that certain features of the network structure can identify the best targets, e.g., proximity to disease-associated nodes, but they cannot predict specific effects. Moreover, they rely on pre-existing network databases, or require additional curation^76,77^ or machine learning^78^ for new network types. Mathematical models such as Ordinary Differential Equations can interrogate the dynamics of the disease, but require precise data to fit model parameters, and so can only handle smaller set of variables^79,80^.

By contrast, our approach combines scalable, executable modelling with transparent, biologically plausible explanations^29,81^ since each of our predictions is derived from biological interactions that can be explained in the context of the overall model, which itself is derived from, and consistent with, experimentally verified observations (Supplementary Table S1). The mechanistic explanations from our model advance the fight against the COVID-19 pandemic by increasing our understanding of why some treatments work and others fail. Our model and screening are readily accessible and are built using the BMA open-source and freely available toolset (www.biomodelanalyzer.org/) and can be updated as new SARS-CoV-2 variants emerge that may exploit different mechanisms to enter cells and replicate or suppress host defence mechanisms^15^.

We deliberately chose to focus on the innate, rather than the adaptive immune response, and specifically in lung epithelial cells. We further prioritised predicting the best targets for optimised drugs, rather than attempting to model the full pharmacodynamics of specific compounds. This allowed us to survey a broad selection of potential therapies for a critical cell type in the severe form of COVID-19. It also allowed us to focus specifically on two scenarios for the innate immune response: first, a low response seen in many patients^9–14^ that we believe characterises the early stage, and second, a persistent and excessive response in the late phase of the disease^16,82^.

We considered this broad scope appropriate at this stage of the pandemic, when there is a need for rapid development of new treatments, but without compromising safety. In the absence of computational screening of the kind we advocate, there has been a focus on only a small number of drugs^83,84^, many screened through expensive Phase III trials^84^ with high-profile safety concerns in the case of Hydroxychloroquine^85^, one of the most studied^83^. Our computational approach can help accelerate the process of screening more drugs and rejecting those with safety concerns, rapidly guiding clinicians to the most likely candidates.

Exemplifying this, amongst the most promising targets for early severe COVID-19 predicted by our model were Camostat and Apilimod, targeting TMPRSS2 and PIKfyve respectively. We show that these drugs are significantly more effective together than alone in suppressing viral entry and replication in Caco-2 cells (Figure 6). Combining these drugs effectively blocks the two key pathways for viral entry, via the cell membrane exploiting TMPRSS2 and via the endosomal route, without which the virus is severely limited in routes into the cell^68^. This novel combination builds on prior work blocking Cathepsin directly^68,86^ and demonstrates its effectiveness with two phase II tested drugs. Host-directed therapies such as this and others suggested by the model may be more resistant to future mutations in SARS-CoV-2^30^.

Our computational model could be developed further to address patient stratification and correct dosing. Many treatments, including those we suggest, depend upon being administered at a specific stage of the disease, making it vital to accurately stratify patients, and necessarily includes factors outside the scope of our model, such as when it is practical to administer a treatment. As an example, Remdesivir and Interferon *α*-2a (Roferon-A) have similar effects in the model, but Interferon *α*-2a may be significantly easier to administer as it is inhaled, and so may be more appropriate in the early case, compared to Remdesivir that requires intravenous administration. The likely side-effects of treatments must also be considered; Miltefosine with Ulixertinib is likely to produce diarrhoea and vomiting, which may be a severe burden in an unwell patient and pose aerosol risks to medical staff attending to them. Our network model helps address stratification by identifying potential biomarkers for different stages and severities of disease. It could also be extended to address dosing for both mono- and combination therapies. Finally, as we understand more about the progression of the disease and the set of traits that determine mild versus severe disease, the model could be expanded to better and, in more detail, explore the stages of severe COVID-19.

In this study we have demonstrated that computational modelling has the ability to rapidly screen and predict new and improved therapies for previously unknown and life-threatening conditions. In particular, through this approach, we have listed several new combination therapies, based on existing and approved drugs, with the potential to improve outcomes for COVID-19 patients. The flexibility and transparency of mechanistic computational modelling will allow this work to be further developed as the virus evolves and demands further changes in therapeutic practice.

## Methods

### Qualitative Networks

We model the viral-host interaction network as a discrete qualitative network^31^. Qualitative networks are an extension of Boolean networks^87^ in which each node may take multiple finite values in a fixed range (e.g., 0,1,2) rather than only ON and OFF. We build and analyse this network model using the open-source (MIT License) and freely available BioModelAnalyzer (BMA) tool^88^ http://biomodelanalyzer.org.

The network consists of nodes representing viral proteins and RNA; host genes and proteins; processes such as viral entry; and phenotypes such as inflammation (see Supplementary Text). The interactions between nodes are represented as edges (see Supplementary Table S1). The level of activity of a node is represented as a positive integer within a fixed range specific to that node. The level of activity changes in response to other nodes, as determined by a mathematical function associated with the node called a target function. The target function takes as input the level of neighbouring nodes that have an incoming edge to the node the target function controls. The default target function is *avg(pos)-avg(neg)*. This takes the mean of the level of activity of all the nodes with a positive edge (represented by an arrow (→)) and subtracts the mean of the level of activity of all the nodes with a negative edge (represented by a flat head (⊣)). More complex functions are needed to describe nodes with behaviour such as only activating when an input rises above a threshold. These target functions, and their rationale, are described in Supplementary Table S2. If node *X* has a granularity of *a-b* and is used in the target function of another node *X’* with range *a’-b’*, then it is scaled to the range of *X’* using the equation:

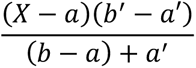

When presenting the values of nodes of differing range in the same plot (Figure 3, 5, Supplementary Figure S1, S5, S12) we normalise the value of each node as a percentage of the maximum value of the node.

For a given set of initial values for all nodes, the network model will update all node values synchronously, and so there will be a single stable attractor the network will tend to. This attractor may be of any number of states, if one we refer to it as a fixed-point attractor, if greater than one we refer to it as a loop. We can test for whether the network reaches a fixed-point attractor for all possible initial states^89^. If there are different attractors for different initial states, we refer to this as a bifurcation. In the case of a fixed point, we take the level of every node in the attractor as the prediction of biological behaviour, in the case of a loop or a bifurcation we use the mean of the tightest bound that can be placed upon node behaviour using the algorithm described in Cook et al^89^. Details of the procedure of network simulation can be found in Schaub et al^31^.

### Model testing

We build the network model using experimental evidence for each edge, described in Supplementary Table S1. We then collate a separate set of testing data used to evaluate the model; this is summarised in Supplementary Table S4. These data describe experiments that do not define a point-to-point interaction between genes and/or proteins, but rather the overall behaviour of the system that emerges from the sum of the interactions.

In order to reproduce an experimental condition in the model, we set the target function of the relevant node to a constant value. For example, the presence of SARS-CoV-2 is represented by changing the target function of *Virion* to 1. We then find the stable states of the model in this case and compare the predicted stable value of relevant phenotype nodes (e.g., inflammation) to that observed in the experiment. We iteratively test and refine the model until it matches all the observed behaviours, as described in Supplementary Table S4.

### *In silico* drug combination screens

In order to find the most effective treatments, we inactivate (set target function to minimum) or activate (set target function to maximum) all nodes, or all sets of nodes targeted by a corresponding drug (Supplementary Table S5) either singly or in a pair-wise combination, using the BMA Command Line tool BioCheckConsole. We consider all combinations of nodes and all combinations of drugs, where a single drug may target multiple nodes. This assumes that drugs are able to act on all their putative targets and does not account for other pharmacodynamic effects that may reduce their efficacy. We evaluate these perturbations by the stable state of the network for the phenotypes of interest, or in the case that the steady state could not be found, the mid-point of the upper and lower bounds BMA could place on the behaviour of that node using the algorithm described in Cook et al^89^. We compare different stages of the disease by using different backgrounds; setting certain nodes to a constant level to reproduce the mild, early severe and late severe forms of the disease (see Supplementary Text, Supplementary Table S3).

### Cell and virus cultures

Caco-2 cells were a kind gift Dr Dalan Bailey (Pirbright Institute). Cells were cultured in Dulbecco’s modified Eagle Medium (DMEM) supplemented with 10% heat-inactivated FBS (Labtech), 100U/ml penicillin/streptomycin. For infections, cells were seeded at 0.2×10^6^ cells/ml. Stocks of SARS-CoV-2 strain BetaCoV/Australia/VIC01/2020 (NIBSC) were generated by propagation on Caco-2 cells and virus titres determined by RT-qPCR for viral E RNA copies as described previously^16^.

### Inhibitor treatment and infections

Caco-2 cells were pre-treated with Camostat Mesilate (ApexBio, B2082), Apilimod (Selleckchem, S6414), Miltefosine (Cayman Chemicals, 63280), Dexamethasone (EMD Millipore, 265005) or DMSO (Sigma) vehicle control at 2x the final concentration for 2h prior to infection in 50μl culture medium. Cells were infected with 1000 E copies/cell SARS-CoV-2 in 50μl, bringing the total culture volume to 100μl and inhibitors to 1x final concentrations as indicated. Cytotoxicity of inhibitors was determined by tetrazolium salt (MTT) assay. 10 % MTT was added to culture media and cells were incubated for 24 h at 37°C. Cells were lysed with 10% SDS, 0.01M HCl and the formation of purple formazan was measured at 570nm.

### Flow Cytometry

Infection levels were measured at 24h by flow cytometry. Caco-2 cells were trypsinised, stained with fixable Zombie UV Live/Dead dye (Biolegend) and fixed with 4% PFA before intracellular staining for nucleocapsid protein. For intracellular detection of SARS-CoV-2 nucleoprotein, cells were permeabilised for 15 min with Intracellular Staining Perm Wash Buffer (BioLegend). Cells were then incubated with 1μg/ml CR3009 SARS-CoV-2 cross-reactive antibody (a kind gift from Dr. Laura McCoy) in permeabilisation buffer for 30 min at room temperature, washed once and incubated with secondary Alexa Fluor 488-Donkey-anti-Human IgG (Jackson Labs). All samples were acquired and analysed using a NovoCyte (Agilent) and NovoExpress 1.5.0 software (Agilent).

### Calculation of drug combination indices

The expression of intracellular SARS-CoV-2 nucleocapsid protein was measured by flow cytometry at 24h post infection and used as a measure of drug effect on viral entry and replication. The data were normalized to the value of the control group and averaged across 9 replicates. For each treatment, the combined effect of drug 1 at concentration *a* and drug 2 at concentration *b* was calculated as *E_a,b_* = (100 – *N_a,b_*)/100, where *N_a,b_* is the average percentage of cells expressing nucleocapsid protein at this concentration. The combination index (CI) was then calculated according to the Bliss independence model ^90,91^:

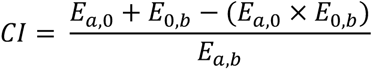

When *CI* < 0, this indicates synergy between the drugs at this concentration, while *CI ≈* 1 indicates additivity and *CI* > 0 indicates antagonism.

## Acknowledgments

The authors wish to thank A.J. Herbert for critical reading and sub-editing of the manuscript. The authors acknowledge support by the National Institute for Health Research University College London Hospitals Biomedical Research Centre to J.F. and D.M.L., Cancer Research UK grant (C17918/A28870) to J.F, Cancer Research UK project grant (28334) to C.B, Wellcome Trust Investigator Award (108079/Z/15/Z) to C.J and grant from LifeArc (COVID0005) for a clinical trial of early antiviral treatment in COVID-19 to D.M.L.

## Supplementary Materials

## Supplementary Text

We developed an executable model of SARS-CoV-2 infection in lung epithelial cells in order to explore combination therapies targeting the virus and host proteins. In this supplementary information we will begin by explaining how the model was constructed and the disease states it models. We will then describe in detail how the host and viral pathways were modelled. Next, we describe how the model was validated against *in vitro* experiments from the literature and clinical trials. Finally, we will discuss how, using the model, we performed *in silico* screening to identify drug combinations to treat SARS-CoV-2 infection.

### Modelling the SARS-CoV-2 infection regulatory network

Reviewing 100 publications describing SARS-CoV-2, SARS-CoV-1 and pan-viral disease mechanisms, we constructed a Qualitative Network model^1^ using the BioModelAnalyzer^2^ tool (Figure 2A, Supplementary Table S1, Supplementary References). This computational network model represents the regulatory network underlying SARS-CoV-2 infection in lung epithelium. The network consists of 175 nodes representing 36 viral proteins and complexes, 102 host proteins and complexes, 31 cellular processes, 5 RNA species and a node controlling the strength of innate immune response (Supplementary Table S6). Of the 31 cellular processes, 8 are considered to be clinically relevant biological processes, which are used to monitor COVID-19 patients’ response to treatment. These nodes are connected by edges that can be either activating (→) or inhibiting (⊣) (Supplementary Table S1).

Each node in the network holds an integer value between 0 and its maximum range, which is 2 for most nodes, and 4 for some symptom nodes and cytokines. The value of each node represents its level of activity, for example when *CCL2* takes a value of 0 it is inactive and when it is at 4 it is at its most active. The level of each node is determined by the activity of its activators and inhibitors, as specified by a target function which expresses mathematically how the node is regulated by its activators and inhibitors (Supplementary Table S2). A generic target function, given by the average of the activators minus the average of the inhibitors, is used when a specific target function is not specified.

There are eight phenotypic readout nodes in the model: *Viral Entry*, *Viral Replication*, *Inflammation*, *Fibrosis*, *Cell Death*, *T-cell Infiltration*, *Syncytia Formation* and *Host Protein Synthesis*. The *Viral Entry* node represents the ability of the virus to enter the cell, while *Viral Replication* represents the production of new virions which can go on to infect other cells. *Inflammation* represents the summed effect of pro-inflammatory cytokines, minus the anti-inflammatory cytokines. *Fibrosis* represents the production of *α*-collagen in the lungs, leading to lung scarring. *Cell Death* represents the level of death in a population of cells via apoptosis, necroptosis or pyroptosis. *T-cell Infiltration* represents the movement of T-cells into the lungs, which is suspected to be the cause of T-cell lymphopenia seen in COVID-19 patients. *Syncytia Formation* represents the formation of cell-cell fusions seen in populations of infected cells. *Host Protein Synthesis* represents the ability of the cell to produce its own proteins.

There are three key input nodes to the model, which are used to describe the condition modelled. The infected state of the model is controlled by the *Virion* node, representing the presence of viral particles in the lungs. A node called *Immune Cells* is used to distinguish the presence of immune cells in tissue from epithelial cell lines, and so allows the comparison of the model to results from pure epithelial cell lines, but also *in vivo* data where immune cells and their reaction to the epithelial cells is recorded. The strength of the innate IFN-I and IFN-III response is thought to be critical to the progression and severity of COVID-19^3^. The *Sensitivity of IFN Response* node is used to convey both the response to, and magnitude of, IFN-I/III production in the range low, medium, high (0, 1, 2). All other unregulated nodes in the model represent constitutively active proteins.

### States of the model

The SARS-CoV-2 infection model can stabilise in one of four states (Figure 2B):

- Uninfected
- Mild COVID-19 disease
- Early stage, severe COVID-19 disease
- Late stage, severe COVID-19 disease

Each of these states is specified by a combination of constraints applied to the model. We describe each of these states and their constraints below.

The **uninfected** state is controlled by setting the *Virion* node to be 0. In this state, all symptom nodes take value 0 (with the exception of *Host Protein Synthesis*, which is 2), representing the levels seen in healthy cells. This state is used as a control to compare with the infected states.

The **mild COVID-19** state represents a patient that is able to mount a robust innate immune response to SARS-CoV-2 infection, and as a result is able to limit Viral Replication in the lungs. This is controlled by setting the *Sensitivity of IFN Response* node to be 1, allowing a normal level of IFN induction in response to infection. In this state, both *Viral Replication* and *Viral Entry* are reduced relative to the early stage of the severe case, as some Interferon Stimulated Genes (ISGs)^4^, such as ZAP and OAS, cause degradation of the viral genome^5^, while others, such as CH25H^6^ and LY6E^7^ are capable of inhibiting entry of the virus into the cell. The *Inflammation* node reaches level 2, driven by both IFN-dependent and virus-dependent pathways. This is considered to be a control case, as such patients do not require medical intervention. However, it is not currently possible to distinguish between mild and severe cases in the early stages of the disease, so any treatment proposed to treat severe cases must be shown not to be damaging to the mild response.

The **early stage, severe COVID-19** case represents patients who are unable to mount a robust IFN response. In this state viral replication proceeds rapidly in the lungs. This state is controlled by setting the *Sensitivity of IFN Response* to 0. In this state *Viral Replication* and *Viral Entry* reach their maximum value of 4. *Inflammation* reaches level 1 in the absence of IFN-dependent inflammation. In this case we search for treatments to limit the spread of infection by reducing the *Viral Replication* node.

There is a growing body of evidence that, although SARS-CoV-2 initially suppresses the IFN response, late stages of infection are characterised by high levels of IFN signalling^8^, corresponding to the **severe COVID-19, late stage** case. This condition is modelled by setting the *Sensitivity of IFN Response* node and the *Viral Genome* node to 2. Setting *Sensitivity of IFN Response* to 2 leads to high levels of IFN induction, which totally dampen both *Viral Replication* and *Viral Entry* but also lead to Inflammation reaching 3. We set *Viral Genome* to 2 to represent that significant viral replication has already occurred by this stage of the disease and so there is a burden of viral RNA present even though replication has peaked. In this case we look for treatment to lower the *Inflammation* node, as this is the most dangerous aspect of this stage of the disease.

### Key cellular processes and signalling represented in the model

Our model focuses on the following key aspects of SARS-CoV-2 replication and pathogenesis:

#### Viral entry

A cell becomes infected when a SARS-CoV-2 virion binds to the cell through interaction between membrane-bound ACE2 and the spike protein^9–11^. Once bound, the virus enters the cell using host factors TMPRSS2, CathepsinL and TPC2. Upon viral entry, the virus introduces the viral genome, wrapped in the nucleocapsid (N) protein, to the cell^12^.

#### Viral translation

The viral proteins can be divided into 3 main categories: structural, non-structural and accessory^13^. There are 4 structural proteins: the spike (S), nucleocapsid (N), envelope (E) and membrane proteins (M), which together with the viral genome form the virion^12^. There are 16 non-structural proteins (nsps 1-16), which perform key tasks such as viral protein processing and replication of the viral genome^13^. There are also a number of accessory proteins, although they are less well understood^12^. Only a subset of the ORFs in this region of the genome are actually expressed as proteins. On the basis of transcriptomic and proteomic evidence we include p3a, p3b, p6, p7a, p7b, p8, p9b and p9c^14–16^.

Two large ORFs: *ORF1a* and *ORF1ab* are translated directly from the viral (RNA) genome into polyproteins pp1a and pp1ab^13^. Two proteases, 3CLpro (nsp5) and PLpro (nsp3) cleave these polyproteins into the constituent nsps. The structural and accessory proteins are translated from small genomic RNAs (sgRNAs) which are transcribed from the genome by RNA-Dependent RNA Polymerase (RDRP). The RDRP activity is performed by a complex formed of nsp7, nsp8 and nsp12^17^. The sgRNAs are then capped in a mechanism involving nsp10, nsp14 and nsp16, allowing their translation into proteins^18^.

#### Viral replication

The virus reproduces by the formation of new virions in the ERGIC (ER-Golgi Intermediate Compartment)^19^. These particles are assembled from the four structural proteins surrounding the viral genome. After formation, the virions exit the cell through exocytosis.

#### Innate immune detection of the virus

Two innate virus sensing pathways are thought to be activated by SARS-CoV-2 in epithelial cells^20^. The first relies on cytosolic Pattern Recognition Receptors (PRRs), namely RIG-I and MDA5 which detect viral RNA, leading to activation of the kinase TBK1^21^. TBK1 phosphorylates the transcription factors IRF3 and IRF7, leading to expression of Type I and Type III IFNs. The second pathway involves the TLR3 receptor, which recognizes dsRNA species that form during replication of the viral genome^21^. This pathway also activates TBK1, as well as promoting inflammation by activating NF-*κ*B, through IKK and IkB*α*^22^.

#### IFN signalling pathway

Type I and Type III IFNs are expressed in lung epithelial cells^3^ but Type II IFNs are only produced in some immune cells^23^. However, all types of IFNs are recognized by epithelial cells. There are multiple IFNs of each type as well as multiple receptors, but to simplify the model we consider only the different types of IFN and a single receptor for each type. Upon activation of the receptor, a signal is passed through JAK-STAT signalling leading to expression of Interferon-Stimulated Genes (ISGs)^4^. In this model we include the ISGs CH25H and LY6E, which inhibit viral entry^6,7^; ZAP and OAS which lead to degradation of viral RNA^4,24^; IFITM which has been shown to inhibit the formation of cell-cell fusions^25^; PKR which can inhibit translation^26^; and iNOS which is a cellular producer of Nitrous Oxide^27^.

#### Cytokine secretion

Alongside the IFNs, the model includes a number of cytokines expressed both directly by epithelial cells and indirectly via immune cells. NF-*κ*B is responsible for the expression of a wide range of cytokines and is a signalling hub on which multiple lines of signalling converge^22^, including the Unfolded Protein Response (UPR)^28^ and IFN signalling^29^. The cytokines represented in the model were chosen to represent those commonly observed in COVID-19 patients and correlated with worse outcomes^30,31^.

#### Cell death

The model includes the pathways regulating apoptosis, pyroptosis and necroptosis. The model contains Caspases-1, 3, 8 and 10, with the activity of Caspase-3 controlling the level of apoptosis^32^. Pyroptosis is controlled by the activity of Gasdermins D and E, which are regulated by the inflammasome and PANoptosome complexes respectively^33^. Necroptosis is regulated by MLKL which is also controlled by the PANoptosome^32^.

#### Renin-angiotensin system

ACE2 is the key adaptor used by SARS-CoV-2 to enter the cell, but it is also an enzyme which is important for regulation of angiotensin, a hormone that regulates blood pressure^34^. In healthy cells, ACE2 converts AngII into Ang1-7, helping to shift the balance in favour of Ang1-7^35^. Interaction of ACE2 with spike proteins is thought to prevent its catalytic activity, leading to an increase in AngII levels. AngII is detected by the AT1R, while Ang1-7 is detected by MasR. An imbalance between the activities of these two receptors leads to ER stress and activation of MAPK and ROCK kinases^36^.

#### Translation of host proteins

Translation of host proteins can proceed via both cap-dependent and cap-independent pathways^37^, while viral proteins rely on cap-dependent pathways^38^. Both pathways rely on ribosomal subunits, however they have different requirements in terms of eIFs. The viral protein nsp1 binds to the 40S ribosomal subunit and causes preferential translation of viral transcripts^39^.

#### Endoplasmic Reticulum (ER) stress

ER stress is caused both by imbalance of the Ang1-7:AngII ratio and also by the S and p3a proteins^35,40,41^. ER stress leads to activation of the Unfolded Protein Response (UPR)^28^. The UPR has 3 pathways: ATF6 which acts to decrease protein translation; and IRE1 and XBP1 which cause inflammation.

### Separation of initial infection and IFN response in the model

In order to distinguish between initial viral entry and viral entry after the response induced by the innate immune system, we use two separate nodes. The *Viral Entry1* node represents the initial entry of the virus into the cell and is uninhibited by the IFN response, while the *Viral Entry* node shows the effects of this IFN response. If the model stabilises in a state where *Viral Entry1* and *Viral Replication* are active but *Viral Entry* is fully inhibited, we interpret this as showing that a small population of cells was initially infected but that the subsequent IFN response prevented a chain reaction of infection throughout the wider population of cells. This effect is required to explain the fact that cells treated with high does of IFN-I are resistant to SARS-CoV-2 infection despite the presence of ZAP siRNA^24^. Similarly, we include a *Viral Genome Degradation* node to represent the effects of ISGs that restrict viral RNA. This node inhibits the *Viral Replication* node, representing the requirement of the virion for intact viral RNA, but not the *Viral Genome* node, which represents the initial proliferation of viral RNA.

Since our model runs from all possible initial states, IFN-I and NF-*κ*B could self-activate each other^22,29^. To prevent this undesired behaviour, we model the induction of IFNs through the innate immune sensing machinery alone and discount positive feedback occurring through NF-*κ*B, as the former interaction is more relevant in the early stage of the severe disease. Conversely, in the late stage, IFN activity is high due to many mechanisms, some of which remain unknown, driving a persistent and maladaptive inflammatory response. This is represented by the high level of the *Sensitivity of IFN Response* node. Taken together, these choices allow the model to represent ongoing feedback alongside the initial stimulus, giving greater insight into the underlying process.

### Validating the model against publicly available data

In order to establish whether the model faithfully reproduces experimental data, we tested it against data from scientific literature. We collected 81 experimental observations derived from 21 published and pre-print studies using SARS-CoV-2, specific viral proteins, SARS-CoV-1 and relevant host factors. The full list of experimental observations we used to test the model is summarised in Supplementary Table S4. We reproduced the conditions described in each experiment as constraints on the model by fixing the nodes representing the experimental variables to a specific value. For example, the effect of an inhibitor was modelled by setting the relevant node to the constant value of 0. We then confirmed whether the resulting steady state of the model matched the experimental results. For example, Hoffmann, M et al.^42^ used the protease inhibitor E-64d to inhibit CathepsinL in E6 Vero cells transformed with TMPRSS2. To reproduce this, we set *TMPRSS2* to 1, and *CathepsinL* to 0. Hoffmann, M et al.^42^ showed that this treatment reduced viral entry; indeed our model predicts that E-64d reduces the level of *Viral Entry* from 2 to 1, matching the experiment.

Our model reproduced 74 experimental observations out of the 81 tested. We have explanations for five of the seven cases that diverge. Zhang, X. et al.^41^ shows that IFN-I is induced by expression of p3a, which the model cannot recapitulate, most likely as in our model NF-*κ*B does not induce IFNs. Similarly, Thorne L. G. et al.^43^ found that inhibition of NF-*κ*B with the IKK inhibitor TPCA-1 reduced IFN expression, a finding that is also not recapitulated by the model. Furthermore, Zhang, X. et al.^41^ also found that expression of p3a leads to production of TNF-*α* in A549. The model assumes that TNF-*α* is produced only by macrophages^44^, meaning it will not be produced in an epithelial cell line. This is also the reason that the model does not reproduce the findings of Hsu, A. C.-Y. et al.^40^, that TNF-*α* is induced by expression of S protein in a minimally immortalized epithelial cell line, BCi-NS1.1. The model is also unable to reproduce the findings of Hsu, A. C.-Y. et al.^40^ that S-induced induction of various cytokines is suppressed by the ERK inhibitor, Ulixertinib. Banerjee, A. K. et al.^39^ show that nsp8 and nsp9 inhibit IFN release by binding to the 7SL RNA component of the Signal Recognition Particle (SRP), which is represented in the model. They then show that nsp8 and nsp9 inhibit the ISG response of cells treated with IFN. Inhibition of the SRP by nsp8 and nsp9 is represented in the model but this mechanism is not sufficient to explain the experiment described so we cannot fully reproduce the experiment. Thorne L. G. et al.^43^ found that IL-6 production is not affected by inhibition of MDA5 or Jak1/2, a disagreement with the model we cannot explain at this point.

Having tested the model against experimental data, we then checked the behaviour of commonly used drugs using the model. Remdesivir is a nucleoside analogue that blocks RDRP and has shown promise against SARS-CoV-2 in cell lines^45^ and as a treatment for COVID-19^46^. We tested the effects of Remdesivir by setting the RDRP complex nsp12-nsp7-nsp8 to 0 and checked the steady state of the model in each of the clinically relevant conditions (Figure 3A). We specifically checked each of the eight biological processes nodes and compared their value to the untreated state. In this case, the model recapitulates the activity of Remdesivir, showing a total loss of Viral Replication. We repeated this analysis for Lopinavir, Tocilizumab, Dexamethasone, Roferon-A and Ruxolitinib.

Having established that the model captures the behaviour of commonly used drugs, we were interested to see if the model can predict the impact of genetic risk factors associated with increased risk of severe COVID-19. Zhang, Q. et al.^47^ found that mutations in the IFN-I pathway, including IFNAR, lead to an increased risk of severe disease. Pairo-Castineira, E. et al.^48^ also identified mutations in IFNAR, as well as OAS. We tested the impact of these genetic background in the same way that we tested the drugs effect (Figure S12). Notably, the model predicts that loss of function mutations in IFNAR or OAS would increase viral replication in mild COVID-19 (Figure S12A, B), suggesting that people with these mutations are at a higher risk of developing severe disease. We also tested the findings of Boudewijns, R. et al.^49^, who used a hamster model of severe SARS-CoV-2 infection to show that in the late stages of disease STAT2-/- hamsters had increased viral load but reduced inflammation, which was in agreement with our model (Figure S12C).

### *In silico* screens of new and improved COVID-19 drug combination treatments

We used the model to simulate the effect of applying drugs individually and in all combinations of two drugs, in each of the four states of the model. The network includes many nodes which are the targets of existing selective therapies, such as Miltefosine, an Akt inhibitor^50^. We identified 75 drugs targeting nodes in the network (Supplementary Table S5), of which 38 drugs had passed phase II clinical trials (Table 1). For each treatment tested and condition we recorded the steady state value of the 8 symptom nodes, or in the case that the steady state could not be found, the mid-point of the upper and lower bounds BMA could place on the behaviour of that node using the algorithm described in Cook et al ^51^. We focussed our analysis on two specific parameters and conditions: viral replication in the early stage of the severe case and inflammation in the late stage of the severe case.

First, we considered the effect of each of the 38 drugs individually on viral replication in the early severe case (Figure 4A). We plotted the results as a heatmap, showing the level of *Viral Replication* after each treatment. This showed that many drugs in consideration as treatments for COVID-19, such as Baracitinib^52^, were effective in our model. However, in order to effectively treat patients at this stage of disease, it may be necessary to administer these drugs as post-exposure prophylaxis. We then turned our attention to the combination therapies (Figure 4B). Here we again plotted the results as a heatmap, with each square showing the value of *Viral Replication* upon treatment with the drugs shown on the x and y axis. We looked specifically for combinations that improved upon the single therapies, for example the combination of Dexamethasone and Ulixertinib. We repeated this analysis for Inflammation in the severe, late infection condition (Figures 4C, D).

We applied the above analysis to all eight phenotypic readout nodes in the mild, severe early and severe late conditions (Supplementary Figures S2-4). We also extended this analysis to the full list of 75 targeted drugs (Supplementary Figures S8, 9). Furthermore, we tested the impact of inhibiting all nodes individually and in combination, even where no drugs targeting these proteins currently exists (Supplementary Figures S10, 11). The column “Number_of_perturbations” indicates whether a single or pairwise combination was applied; the column “Disease_State” indicates mild, early severe or late severe disease state; the column “Perturbation_A” indicates the first perturbation applied; the column “Level_A” indicates the level the first node is set to, or not applicable (NA) as the effect is defined in Supplementary Table S5; the column “Perturbation_B” indicates the second perturbation applied or not applicable (NA); the column “Level_B” indicates the level the second node is set to or not applicable (NA); the column “Lower bound” shows the lower bound for the phenotype measured determined by the algorithm; the column “Upper bound” shows the upper bound for the phenotype measured determined by the algorithm and the column “Midpoint” shows the midpoint between the upper and lower bound.

In order to examine the effect of each drug combination on all the biological processes modelled, we plotted these values for each combination as a radar chart (Figure 5, Supplementary Figure S1). In these plots, the biological processes nodes are arranged in a circle and the value reached with a given treatment is plotted as point, which are then joined together between all biological processes. A representative monotherapy is plotted in grey with the combination therapy plotted in colour (orange for treatments identified for severe, early cases and red for treatments identified for severe, late cases).

### *In silico* simulation of Caco-2 cell drug treatments

We validated the model’s predictions that Apilimod and Camostat; and Dexamethasone and Miltefosine act together to reduce viral replication. We test these predictions using Caco-2 cell cultures as they do not mount a significant IFN response to SARS-CoV-2 infection^53^, providing a model of the severe, early COVID-19 condition. We modelled Caco-2 cells by setting the *Immune Cells* node to 0 as this is a clonal colorectal epithelial cell culture, and the *Sensitivity Of IFN Response* node to 0 to reflect the lack of significant IFN response. These simulations predict that combinations of Apilimod and Camostat, and Dexamethasone and Miltefosine, have a greater effect in combination than alone in Caco-2 cells (Supplementary Table S7).

### Differential characteristics of different states of COVID-19

Individuals with COVID-19 display a wide range of symptoms and disease outcomes, with severity ranging from asymptomatic to fatal^54^. Therefore, there is a need for clinical biomarkers to aid in triage of COVID-19 patients. We investigated the steady states of our model representing the different states of disease to look for differences that could potentially distinguish between mild and severe cases. We applied hierarchical clustering to the level of all nodes in each state (Supplementary Figure S5A). The resulting heatmap reveals clusters of nodes that differ between states of the disease (Supplementary Figure S5B).

Hierarchical clustering identifies IFN-I, IFN-II, IFN-III, TNF-*α*, IL-2, IL-1B, IL-6, IL-8, IL-10, IL-18, CCL2, CXCL9, CXCL10 and CXCL11. All of these factors are raised in the late versus early severe stage of disease, in line with observed increases in IFN-I, IFN-II, IFN-III, IL-2, IL-10 and IL-18 reported in longitudinal studies^8,30^. The model also predicts that these factors are higher in the mild state compared to the early severe state of disease, reflecting the effective immune response in these patients. Many of these factors, especially TNF-*α*, IL-6, IL-8, IL-10 and CXCL10 have been identified as prognostic markers for severity of COVID-19 in hospitalised patients^30,31,55–57^.

---

**Figure S1.**
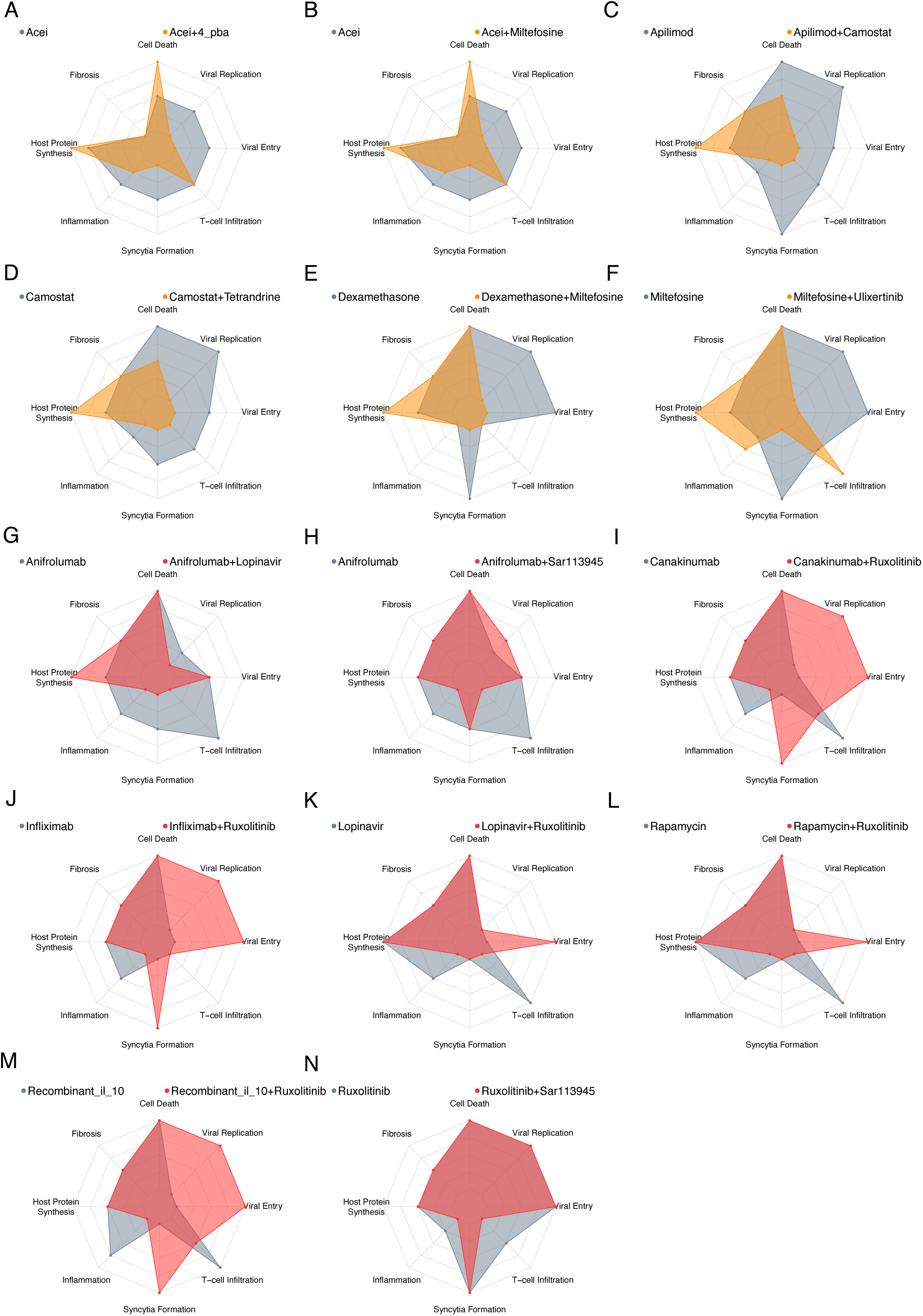
Effect of drug combinations identified in the *in silico* screen on all modelled cellular processes. (A-F) The effect of monotherapy vs drug combinations identified to limit viral replication under the early stage, severe Covid-19. Monotherapy (grey), combination (orange), with the strength of the symptom denoted by radial distance. (D-N) The effect of monotherapy vs drug combinations identified to reduce inflammation under the late stage, severe Covid-19. Monotherapy (grey), combination (Red). All nodes normalized to be between 0 and 100%.

**Figure S2 A-H.**
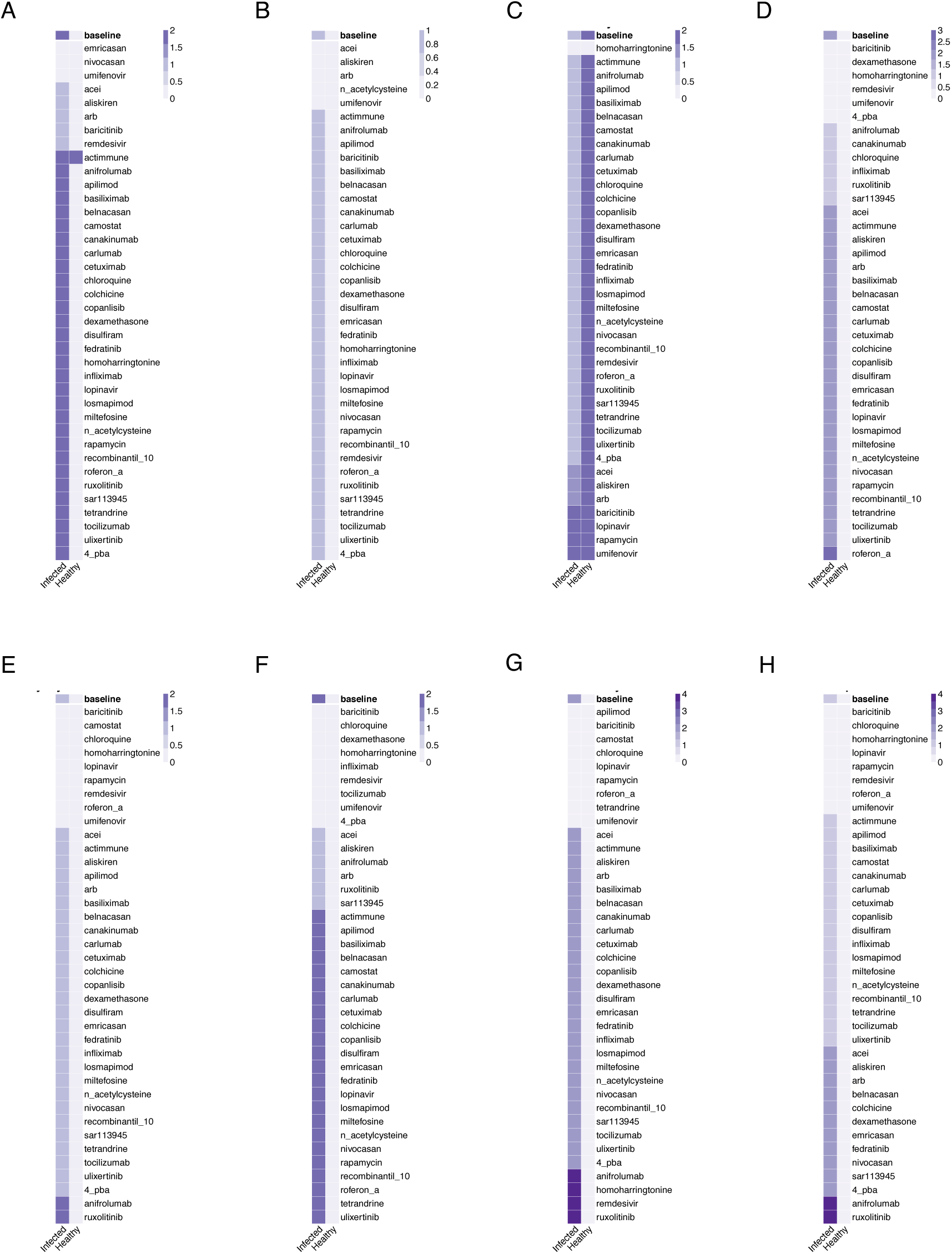
*In silico* screens to identify effects of drugs on mild COVID-19. Columns represent infected and healthy conditions, rows represent drug tested including only those approved for use. Showing the effect on: (A) Cell Death. (B) Fibrosis. (C) Host Protein Synthesis. (D) Inflammation. (E) Syncytia Formation. (F) T-cell Infiltration. (G) Viral Entry. (H) Viral Replication.

**Figure S2I.**
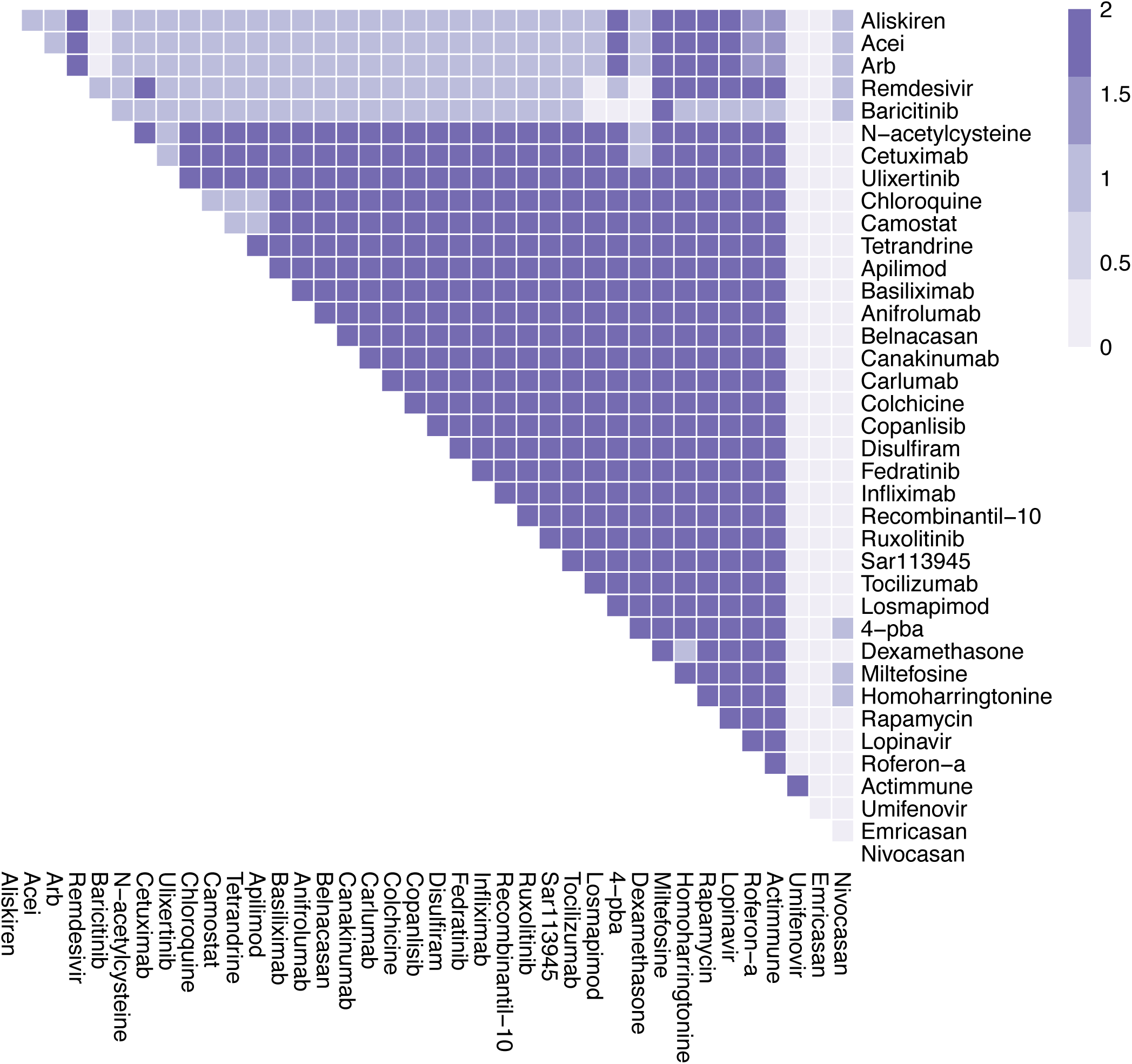
The effects of combination therapy on Cell Death in mild COVID-19. The colour of the square corresponds to the level of the *CellDeath* node at steady state under treatment by the drugs indicated on the x- and y-axis.

**Figure S2J.**
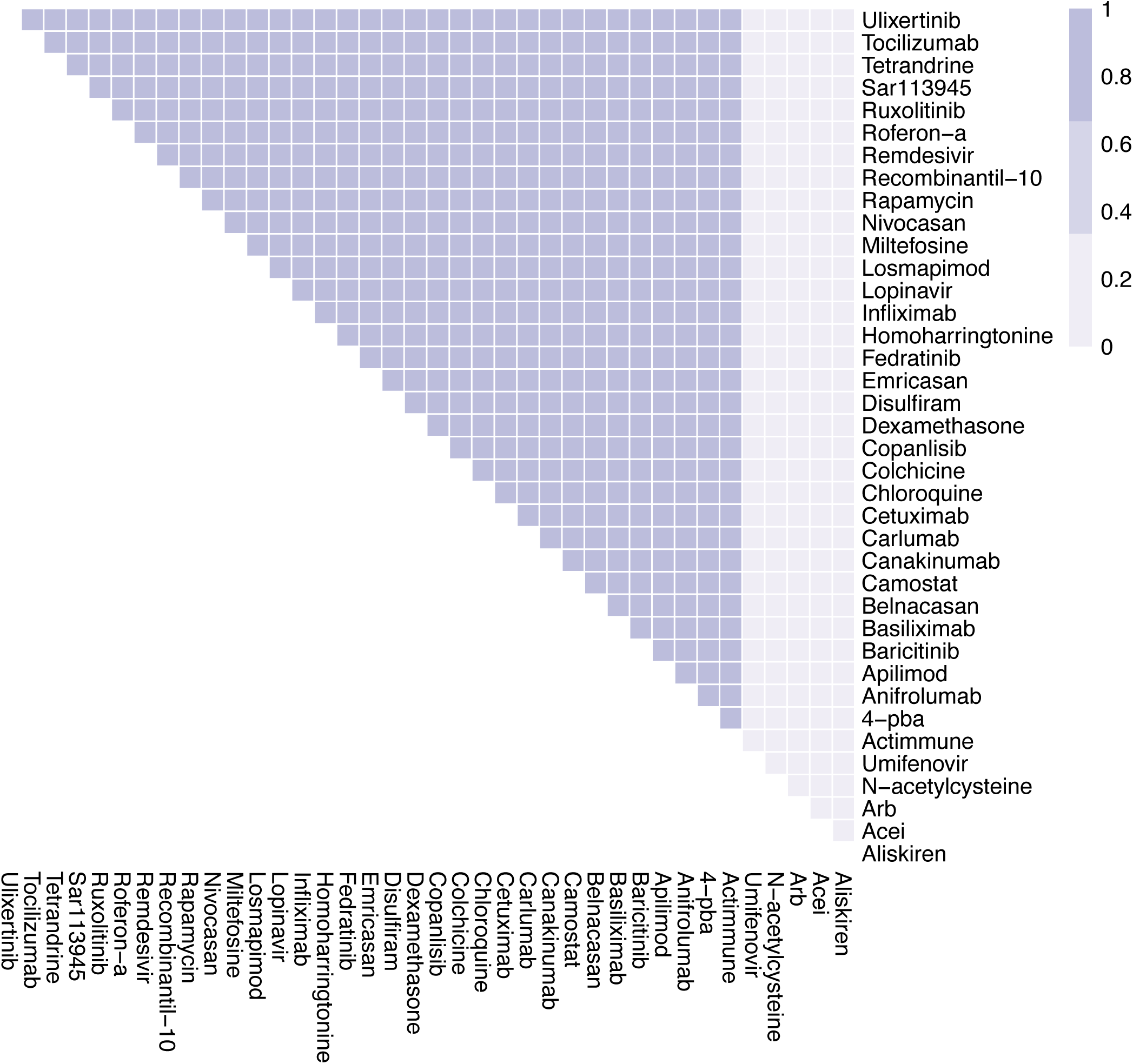
The effects of combination therapy on Fibrosis in mild COVID-19. The colour of the square corresponds to the level of the *Fibrosis* node at steady state under treatment by the drugs indicated on the x- and y-axis.

**Figure S2K.**
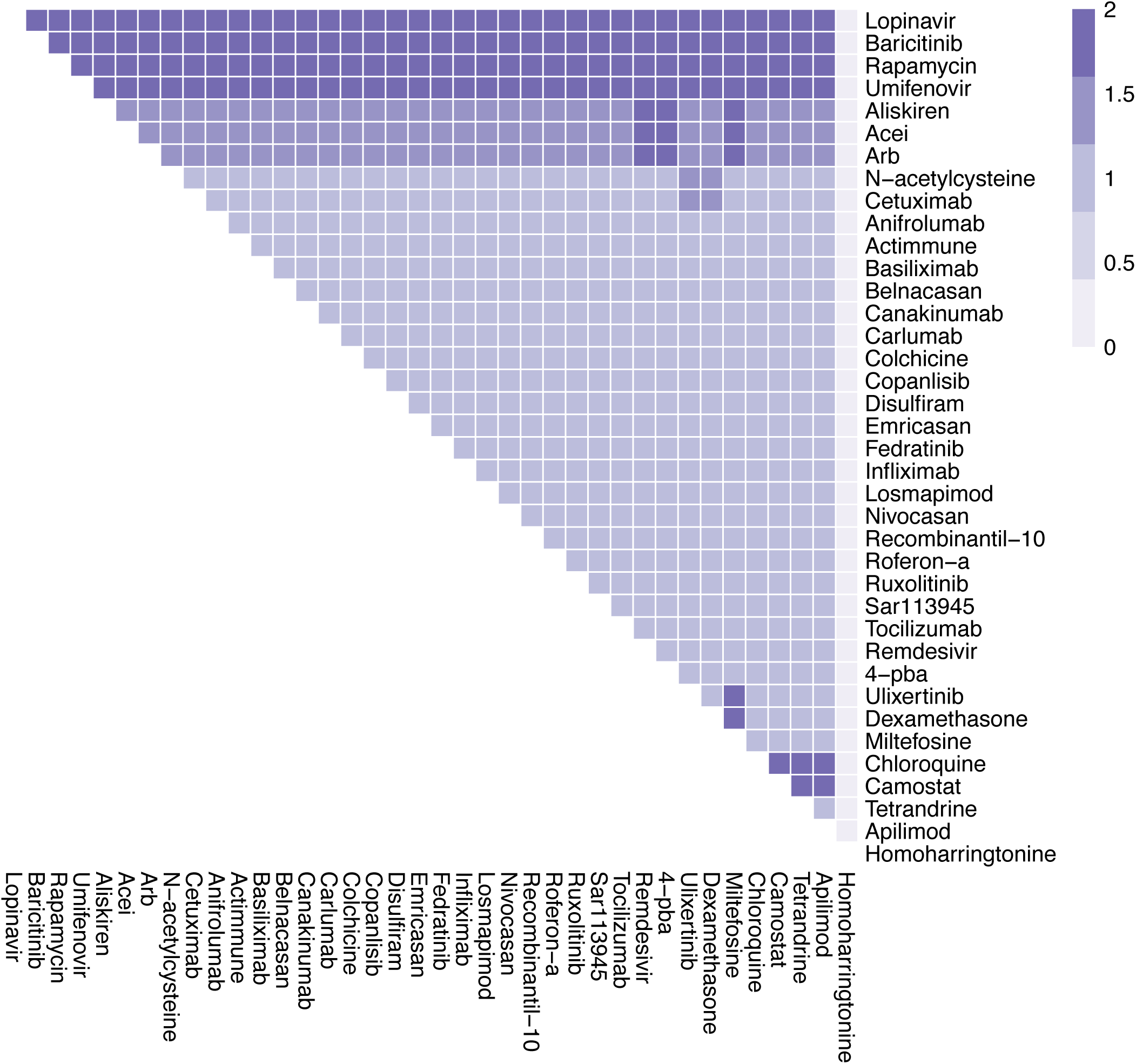
The effects of combination therapy on Host Protein Synthesis in mild COVID-19. The colour of the square corresponds to the level of the *HostProteinSynthesis* node at steady state under treatment by the drugs indicated on the x- and y-axis.

**Figure S2L.**
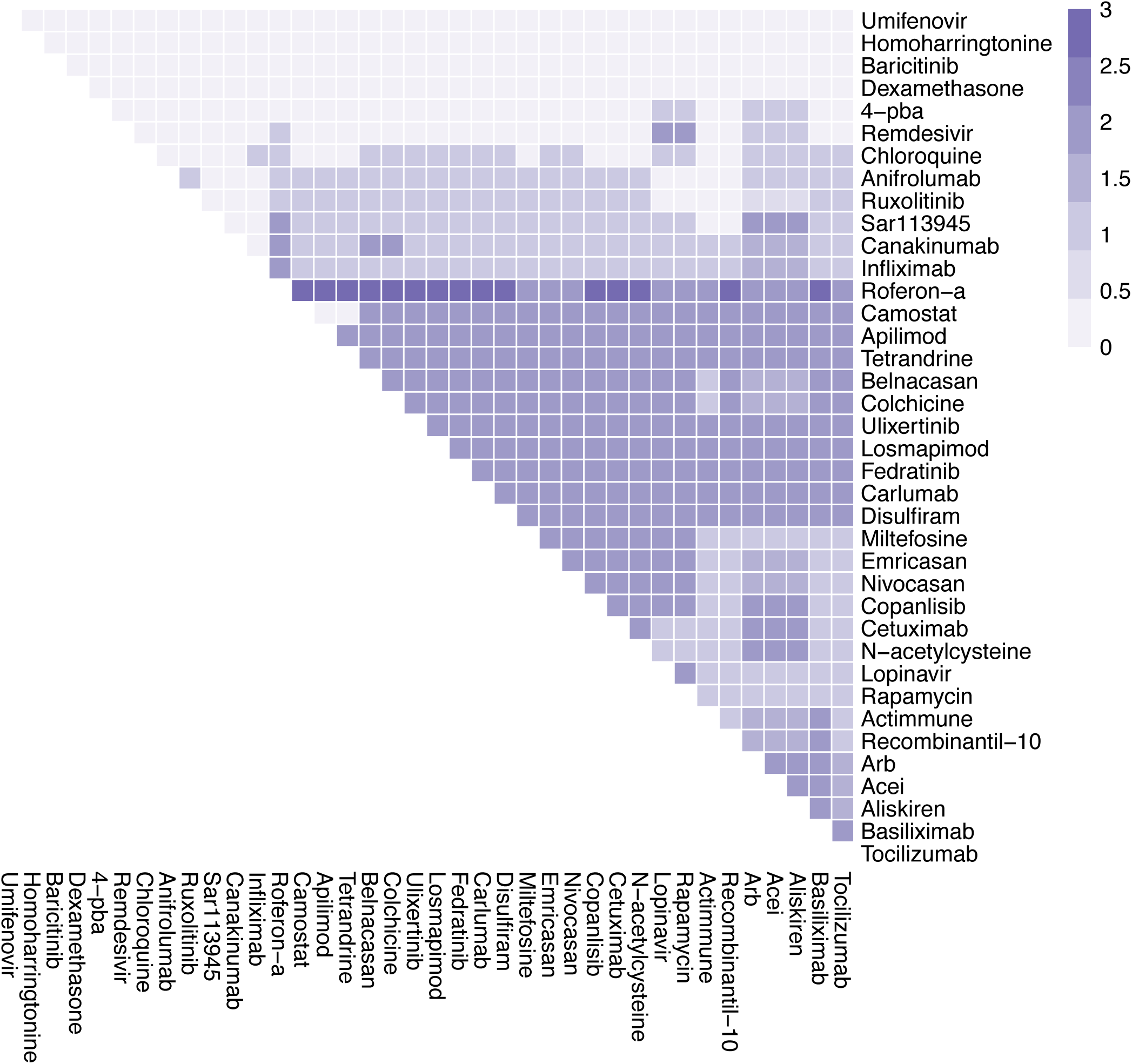
The effects of combination therapy on Inflammation in mild COVID-19. The colour of the square corresponds to the level of the *Inflammation* node at steady state under treatment by the drugs indicated on the x- and y-axis.

**Figure S2M.**
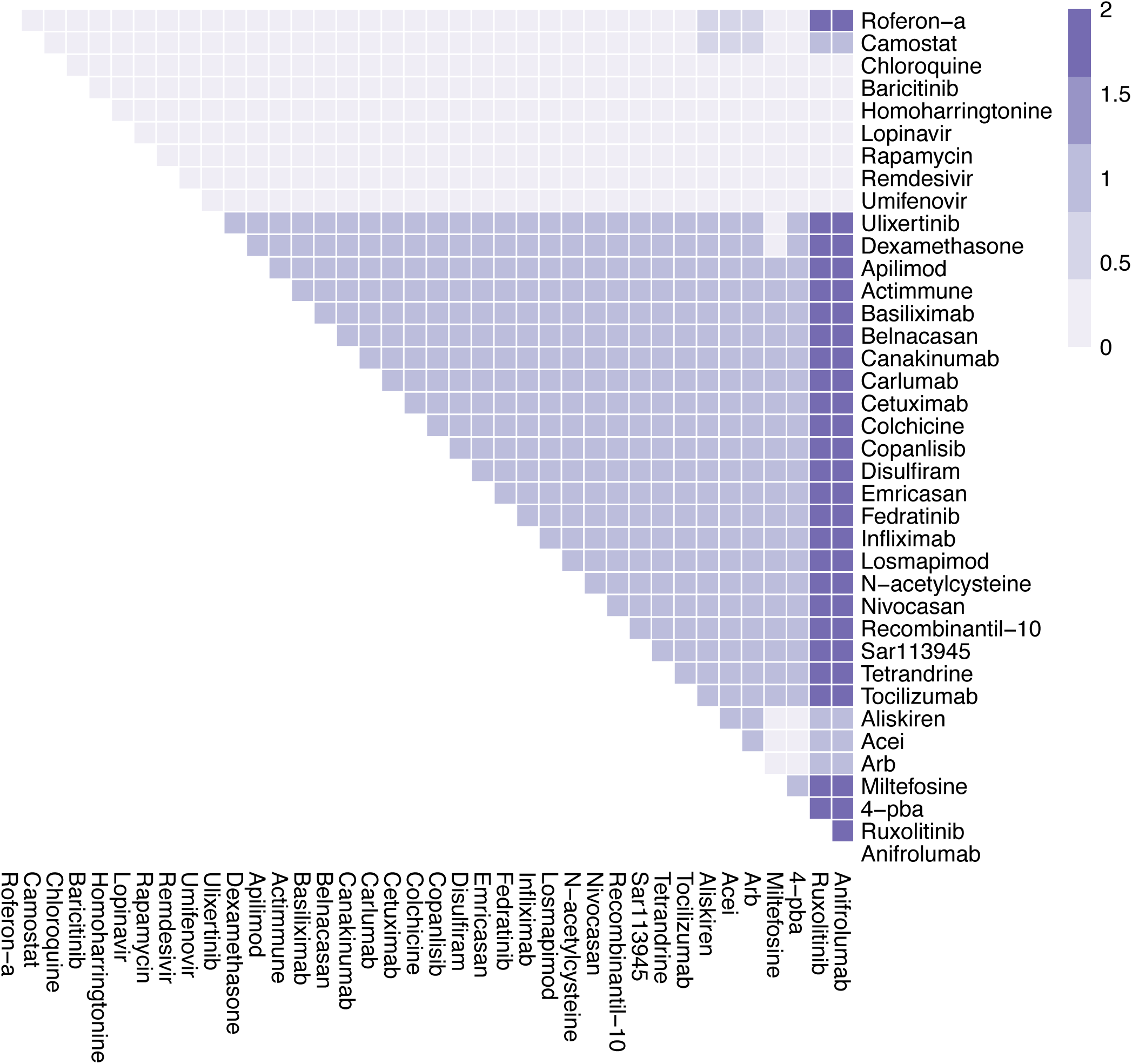
The effects of combination therapy on Syncytia Formation in mild COVID-19. The colour of the square corresponds to the level of the *SyncytiaFormation* node at steady state under treatment by the drugs indicated on the x- and y-axis.

**Figure S2N.**
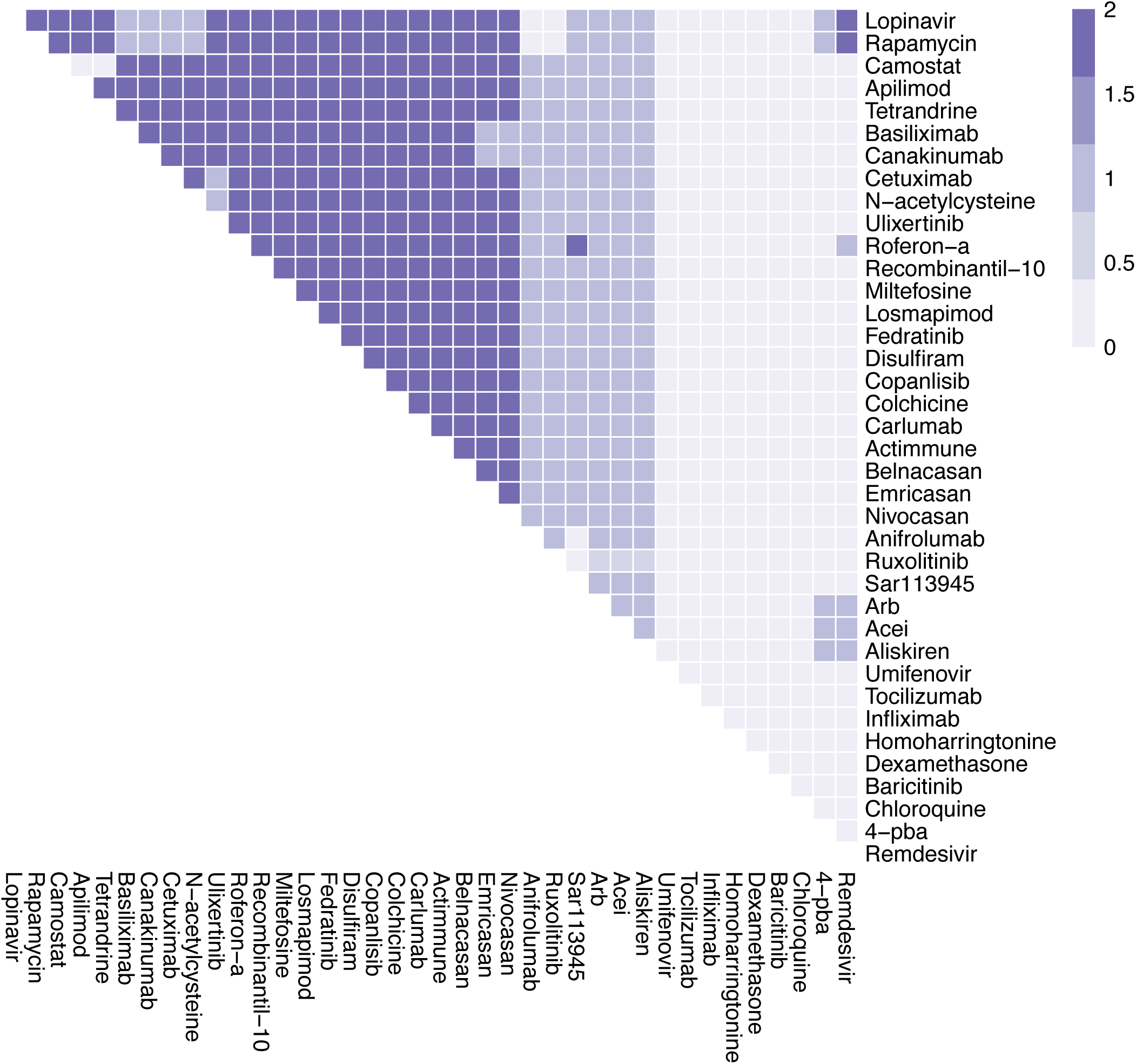
The effects of combination therapy on T-cell Infiltration in mild COVID-19. The colour of the square corresponds to the level of the *T-cellInfiltration* node at steady state under treatment by the drugs indicated on the x- and y-axis.

**Figure S2O.**
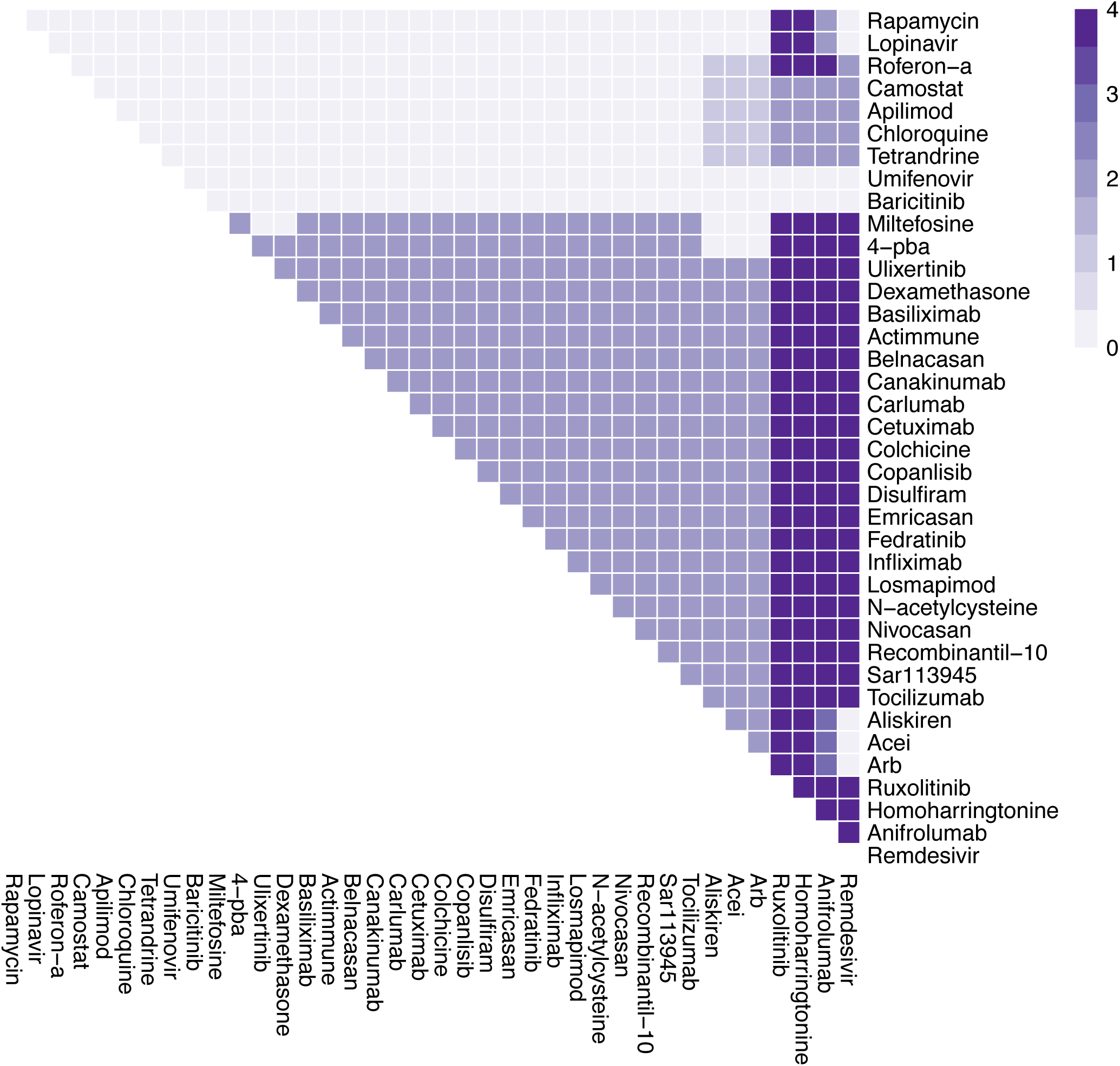
The effects of combination therapy on Viral Entry in mild COVID-19. The colour of the square corresponds to the level of the *ViralEntry* node at steady state under treatment by the drugs indicated on the x- and y-axis.

**Figure S2P.**
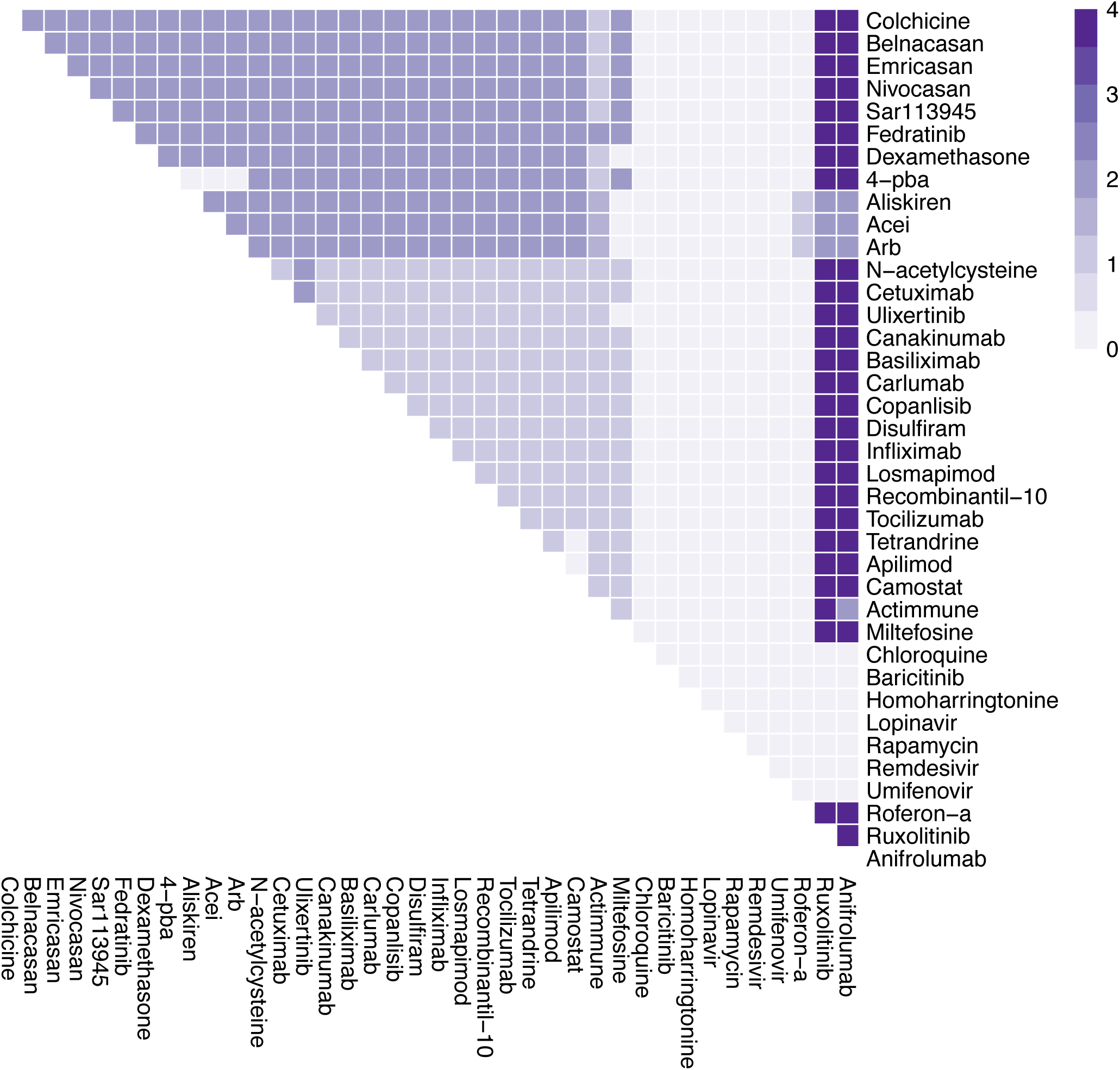
The effects of combination therapy on Viral Replication in mild COVID-19. The colour of the square corresponds to the level of the *ViralReplication* node at steady state under treatment by the drugs indicated on the x- and y-axis.

**Figure S3 A-H.**
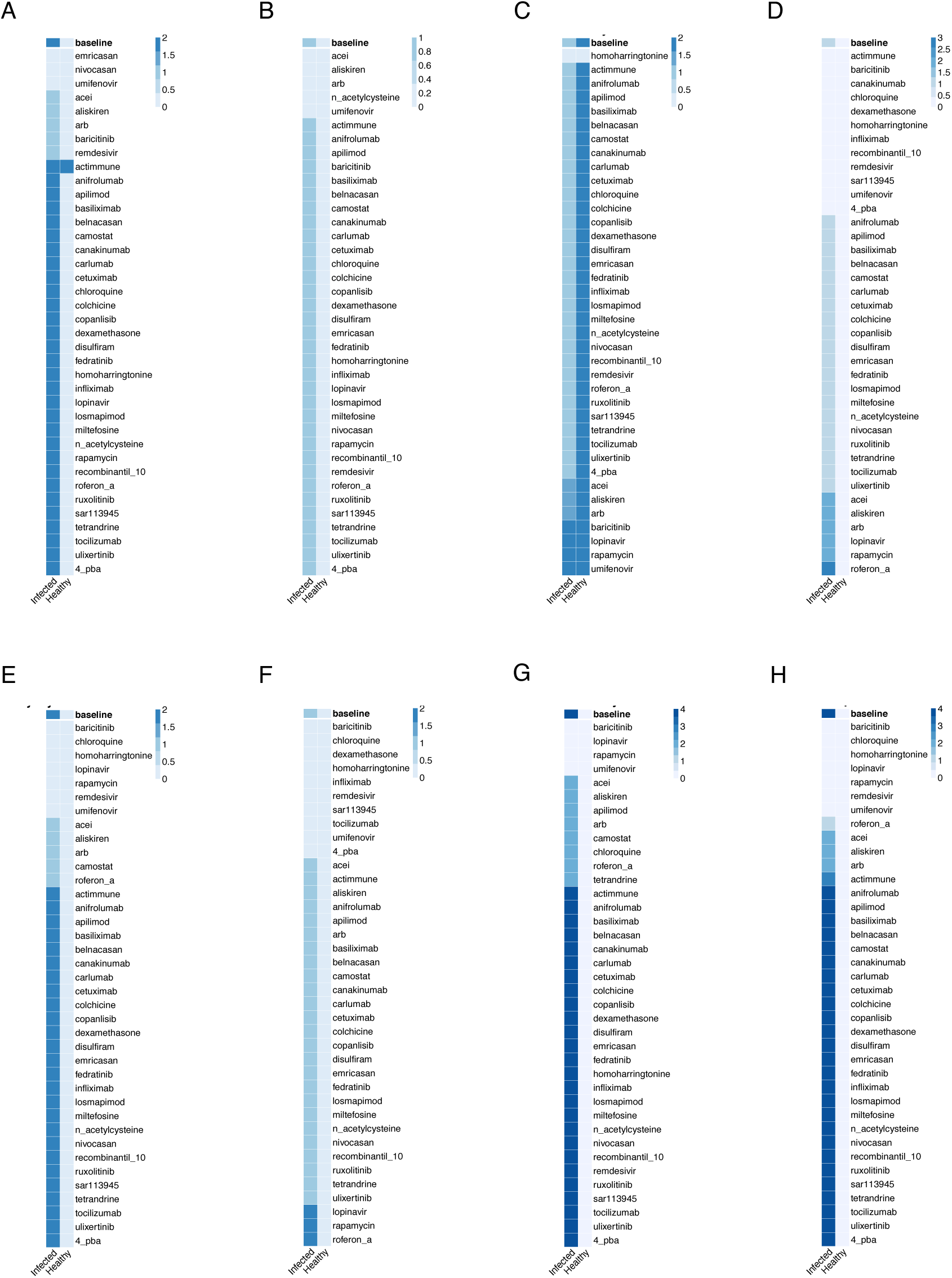
*In silico* screens to identify potential drugs to treat early severe COVID-19. Columns represent infected and healthy conditions, rows represent drug tested including only those approved for use. Showing the effect on: (A) Cell Death. (B) Fibrosis. (C) Host Protein Synthesis. (D) Inflammation. (E) Syncytia Formation. (F) T-cell Infiltration. (G) Viral Entry. (H) Viral Replication.

**Figure S3I.**
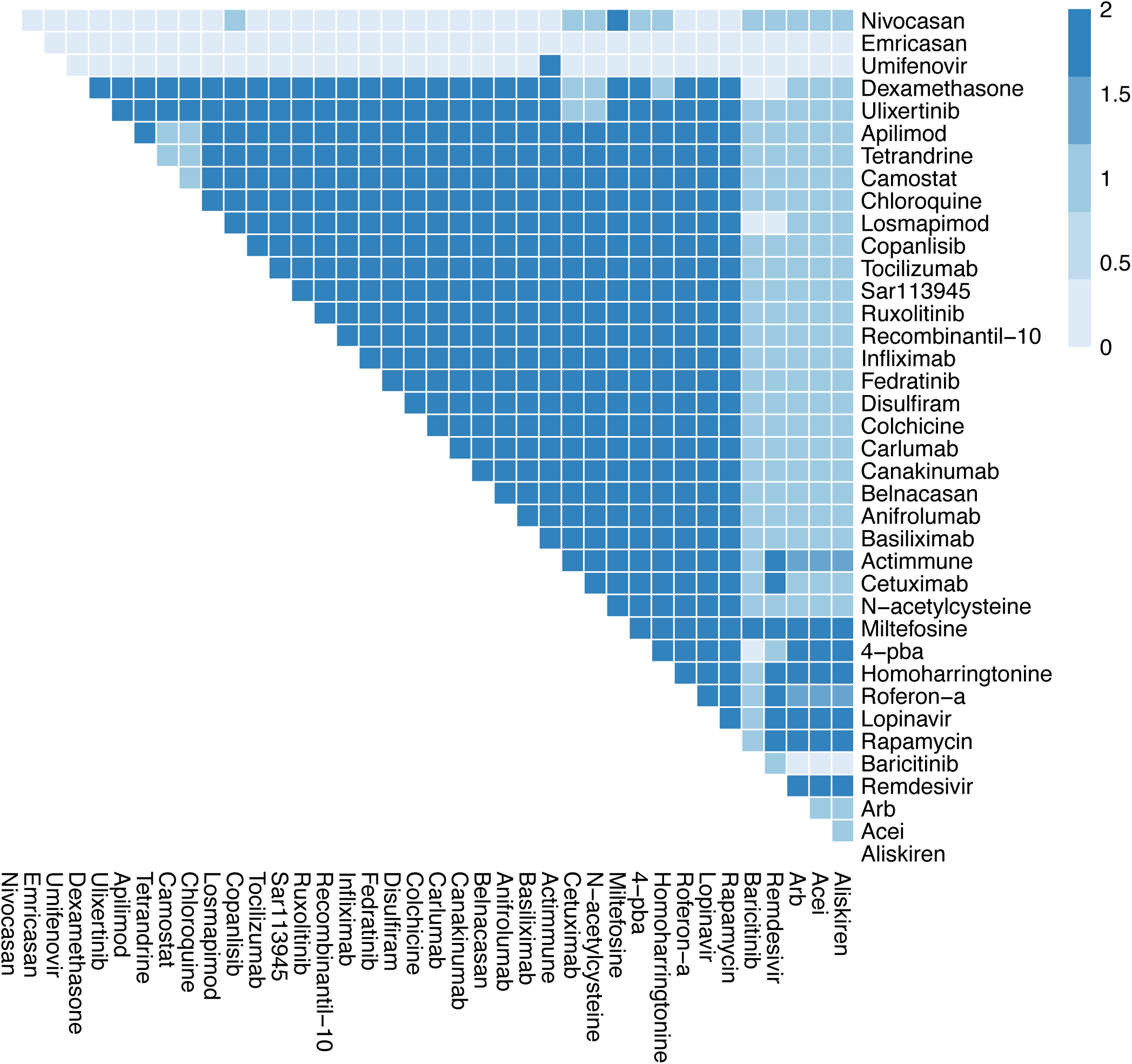
The effects of combination therapy on Cell Death in early stage, severe COVID-19. The colour of the square corresponds to the level of the *CellDeath* node at steady state under treatment by the drugs indicated on the x- and y-axis.

**Figure S3J.**
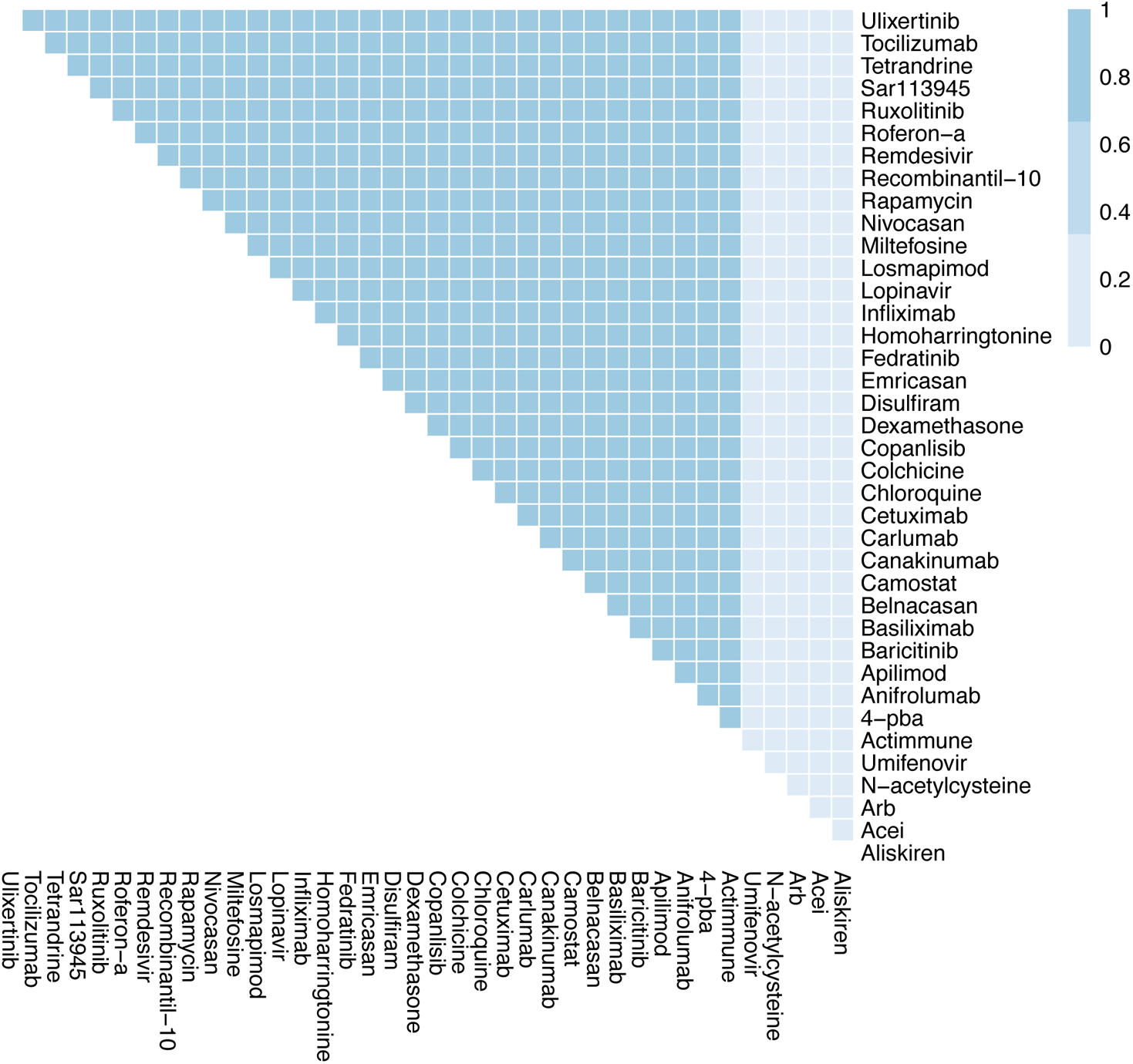
The effects of combination therapy on Fibrosis in early stage, severe COVID-19. The colour of the square corresponds to the level of the *Fibrosis* node at steady state under treatment by the drugs indicated on the x- and y-axis.

**Figure S3K.**
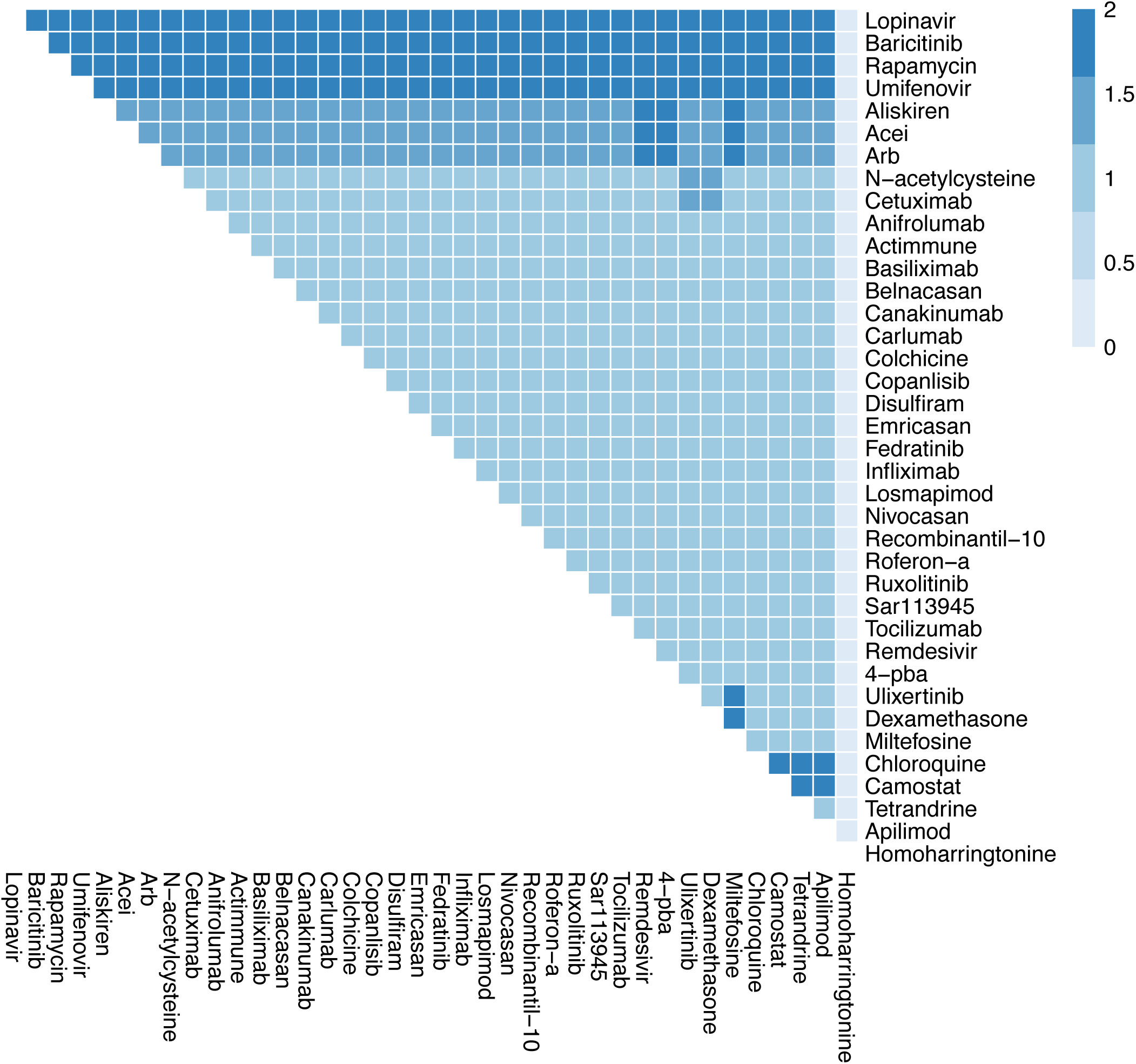
The effects of combination therapy on Host Protein Synthesis in early stage, severe COVID-19. The colour of the square corresponds to the level of the *HostProteinSynthesis* node at steady state under treatment by the drugs indicated on the x- and y-axis.

**Figure S3L.**
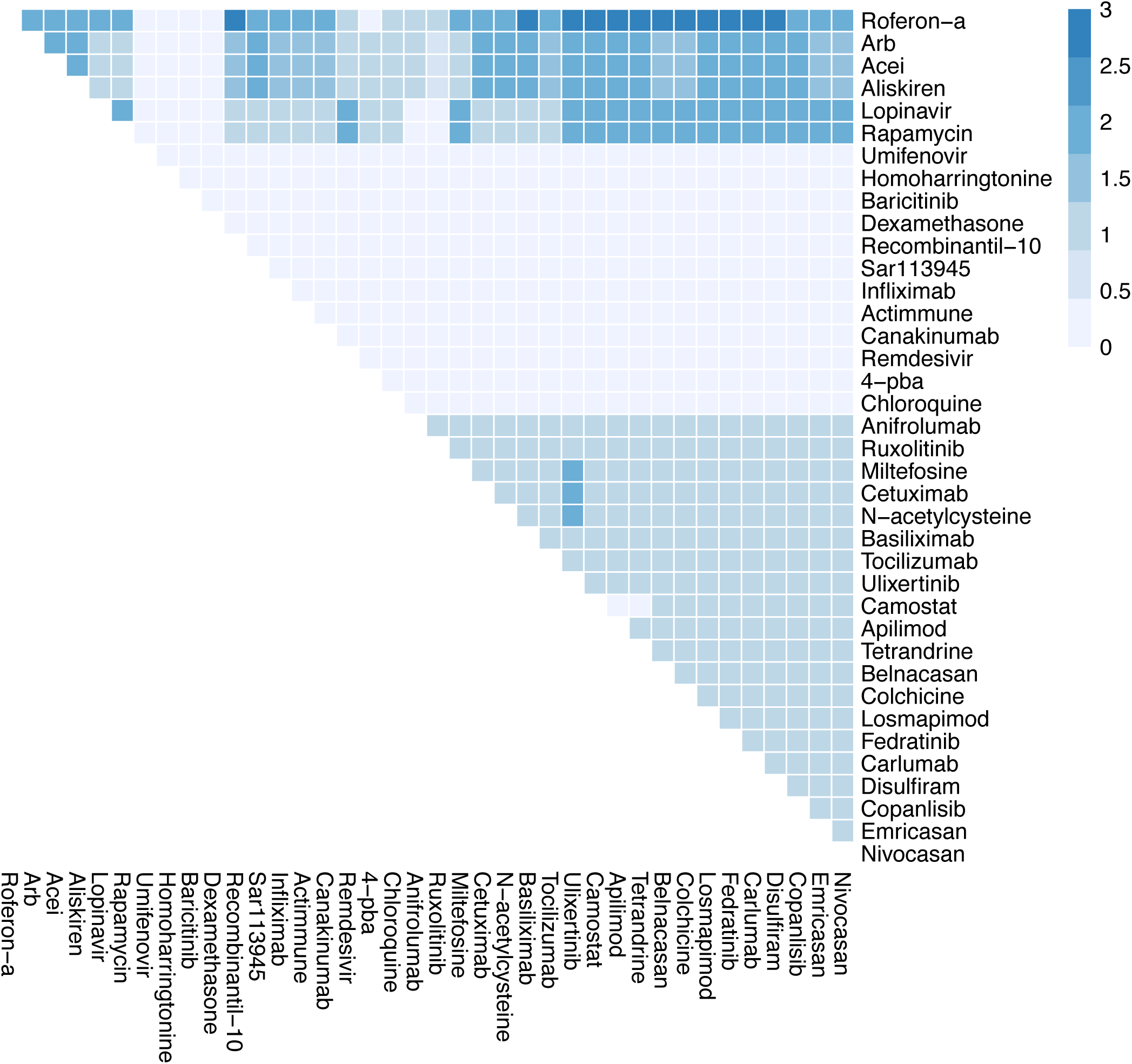
The effects of combination therapy on Inflammation in early stage, severe COVID-19. The colour of the square corresponds to the level of the *Inflammation* node at steady state under treatment by the drugs indicated on the x- and y-axis.

**Figure S3M.**
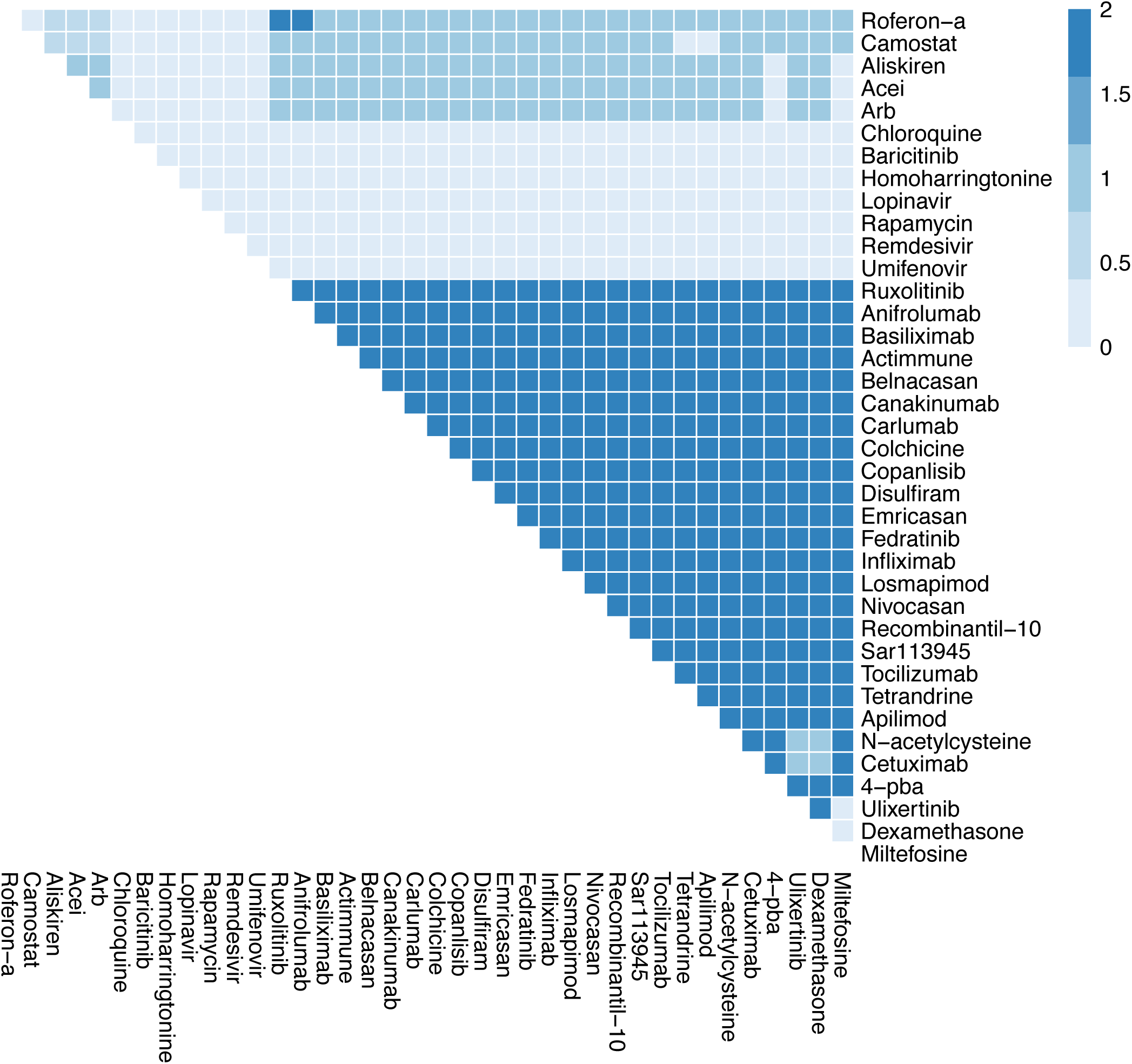
The effects of combination therapy on Syncytia Formation in early stage, severe COVID-19. The colour of the square corresponds to the level of the *SyncytiaFormation* node at steady state under treatment by the drugs indicated on the x- and y-axis.

**Figure S3N.**
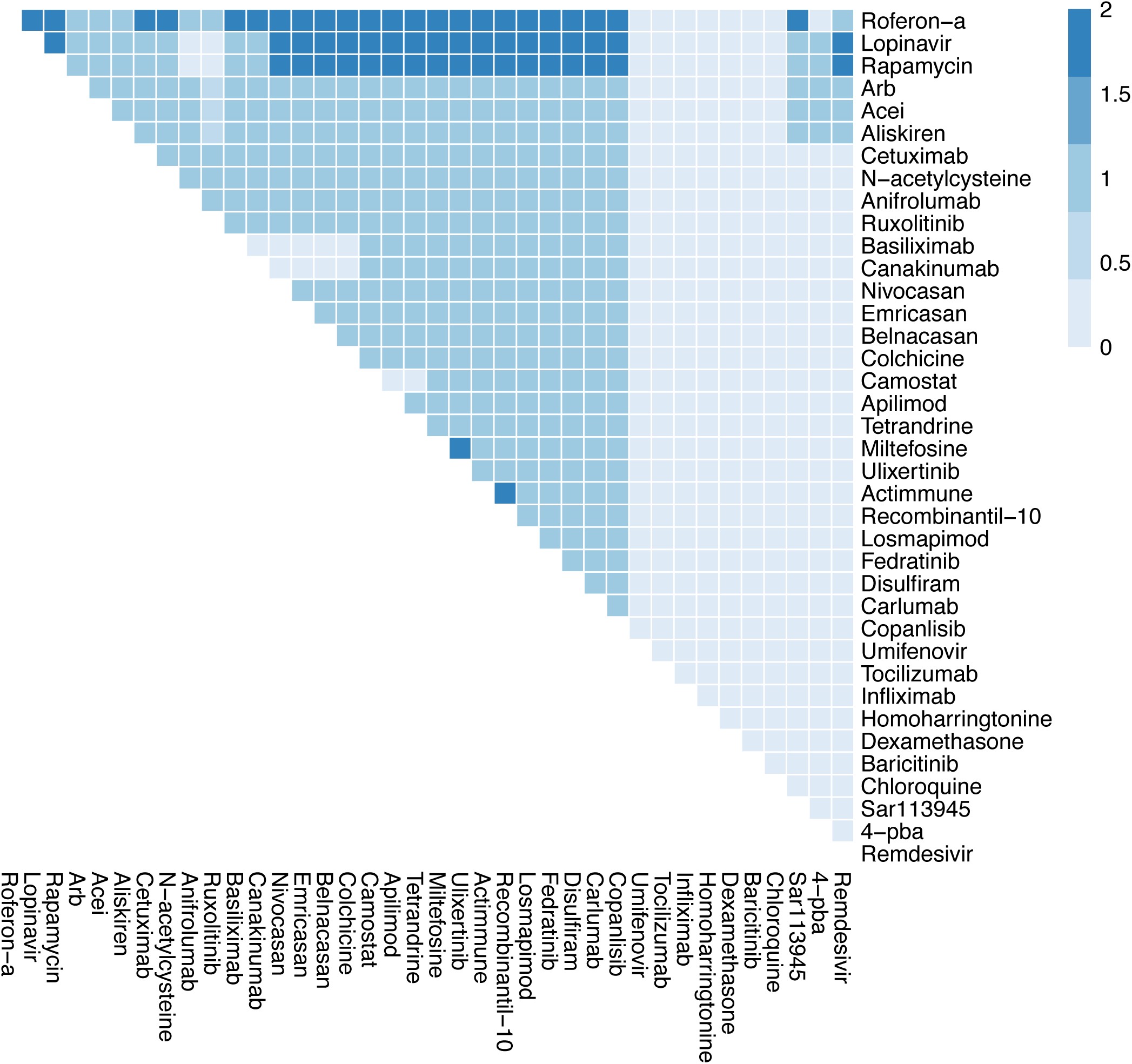
The effects of combination therapy on T-cell Infiltration in early stage, severe COVID-19. The colour of the square corresponds to the level of the *T-cellInfiltration* node at steady state under treatment by the drugs indicated on the x- and y-axis.

**Figure S3O.**
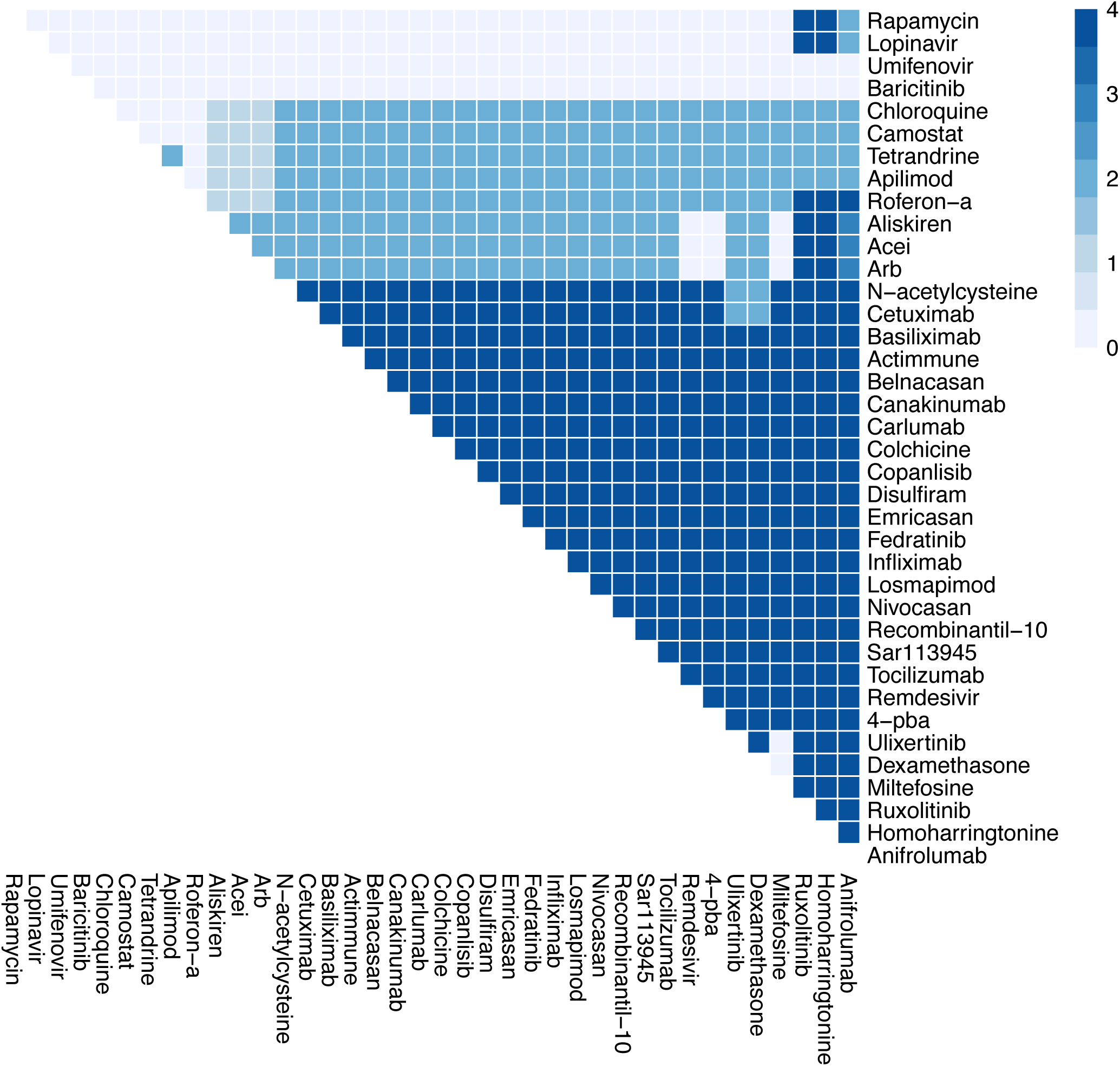
The effects of combination therapy on Viral Entry in early stage, severe COVID-19. The colour of the square corresponds to the level of the *ViralEntry* node at steady state under treatment by the drugs indicated on the x- and y-axis.

**Figure S3P.**
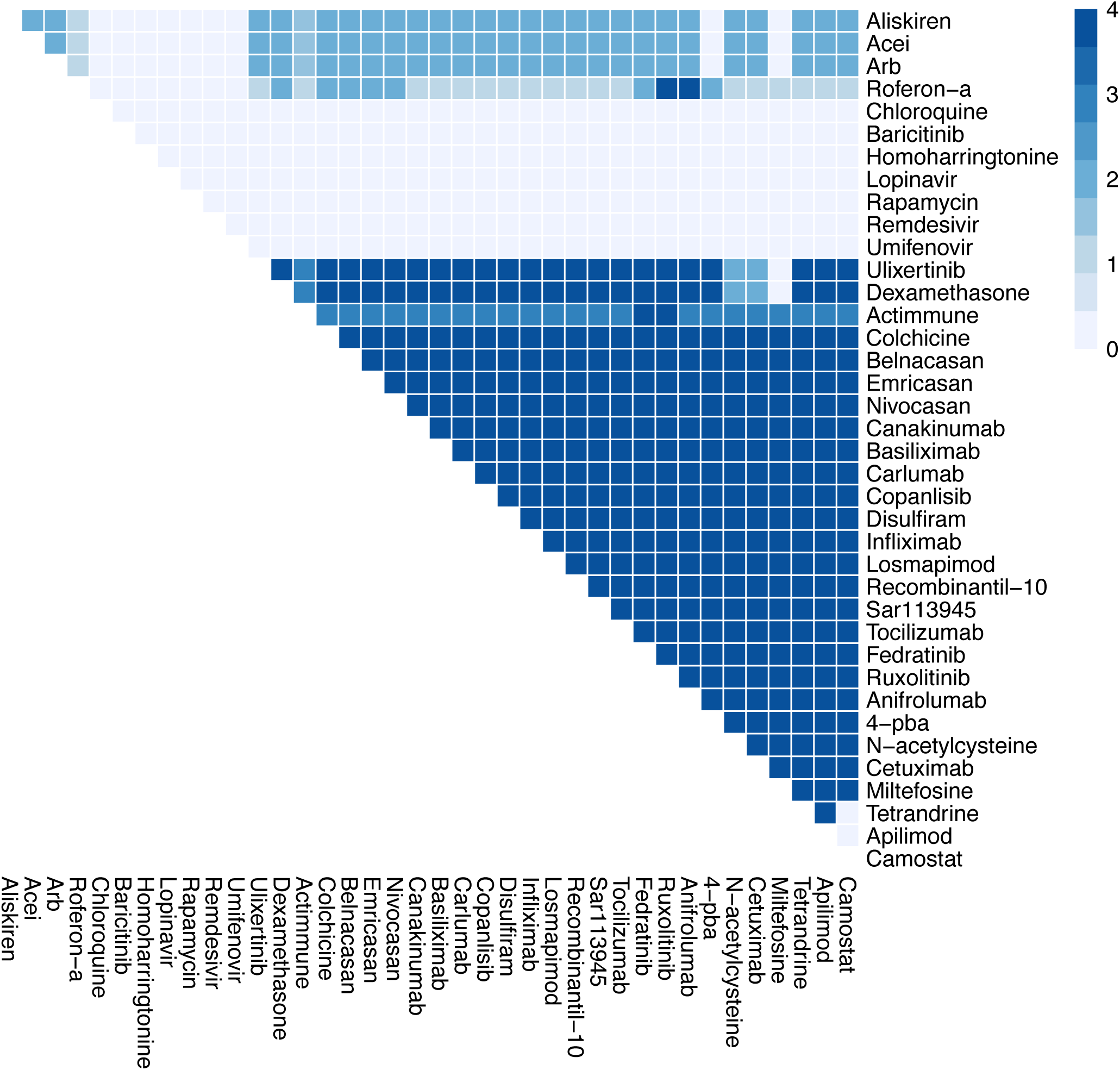
The effects of combination therapy on Viral Replication in early stage, severe COVID-19. The colour of the square corresponds to the level of the *ViralReplication* node at steady state under treatment by the drugs indicated on the x- and y-axis.

**Figure S4 A-H.**
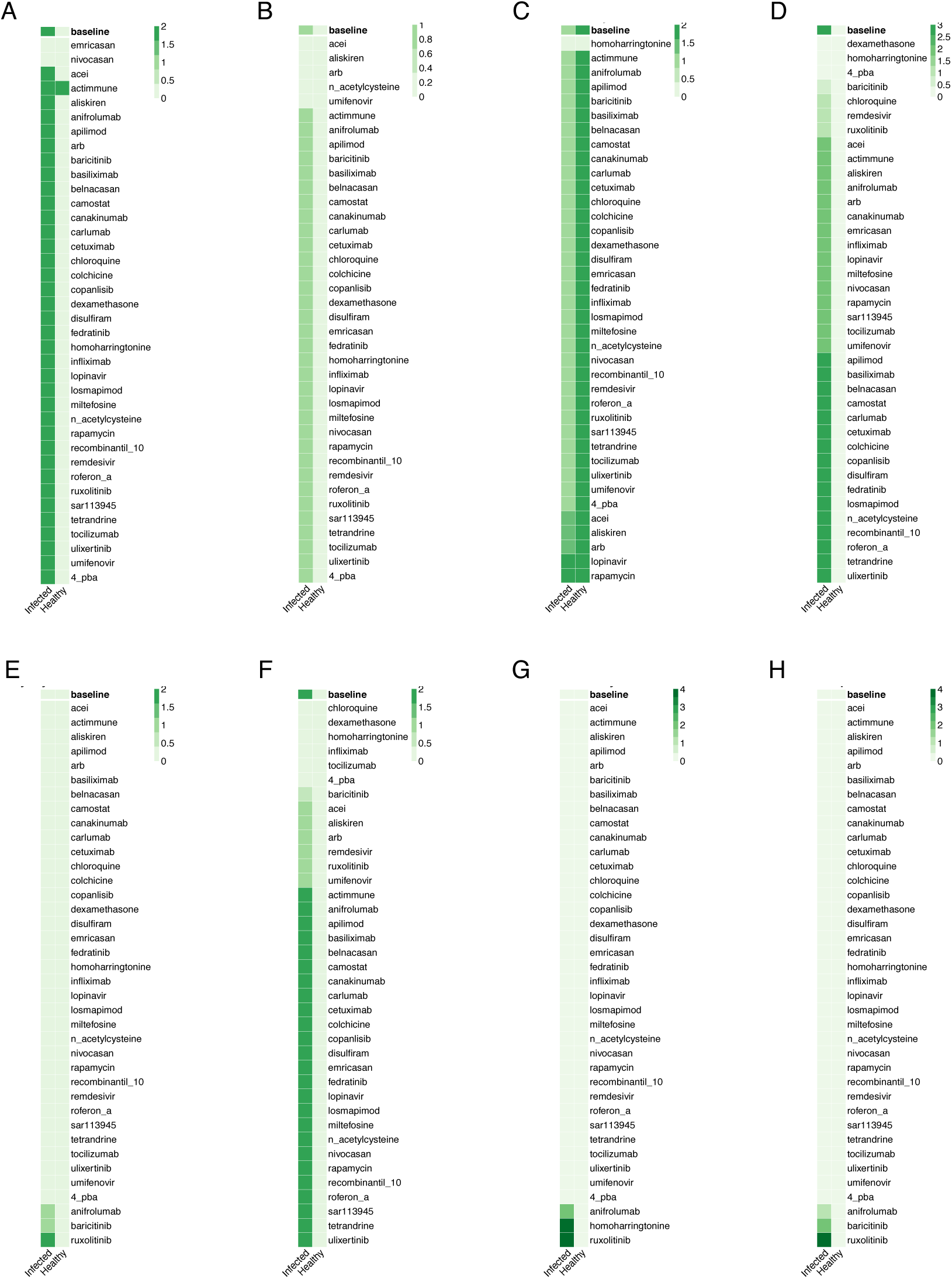
*In silico* screens to identify potential drugs to treat late severe COVID-19. Columns represent infected and healthy conditions, rows represent drug tested including only those approved for use. Showing the effect on: (A) Cell Death. (B) Fibrosis. (C) Host Protein Synthesis. (D) Inflammation. (E) Syncytia Formation. (F) T-cell Infiltration. (G) Viral Entry. (H) Viral Replication.

**Figure S4I.**
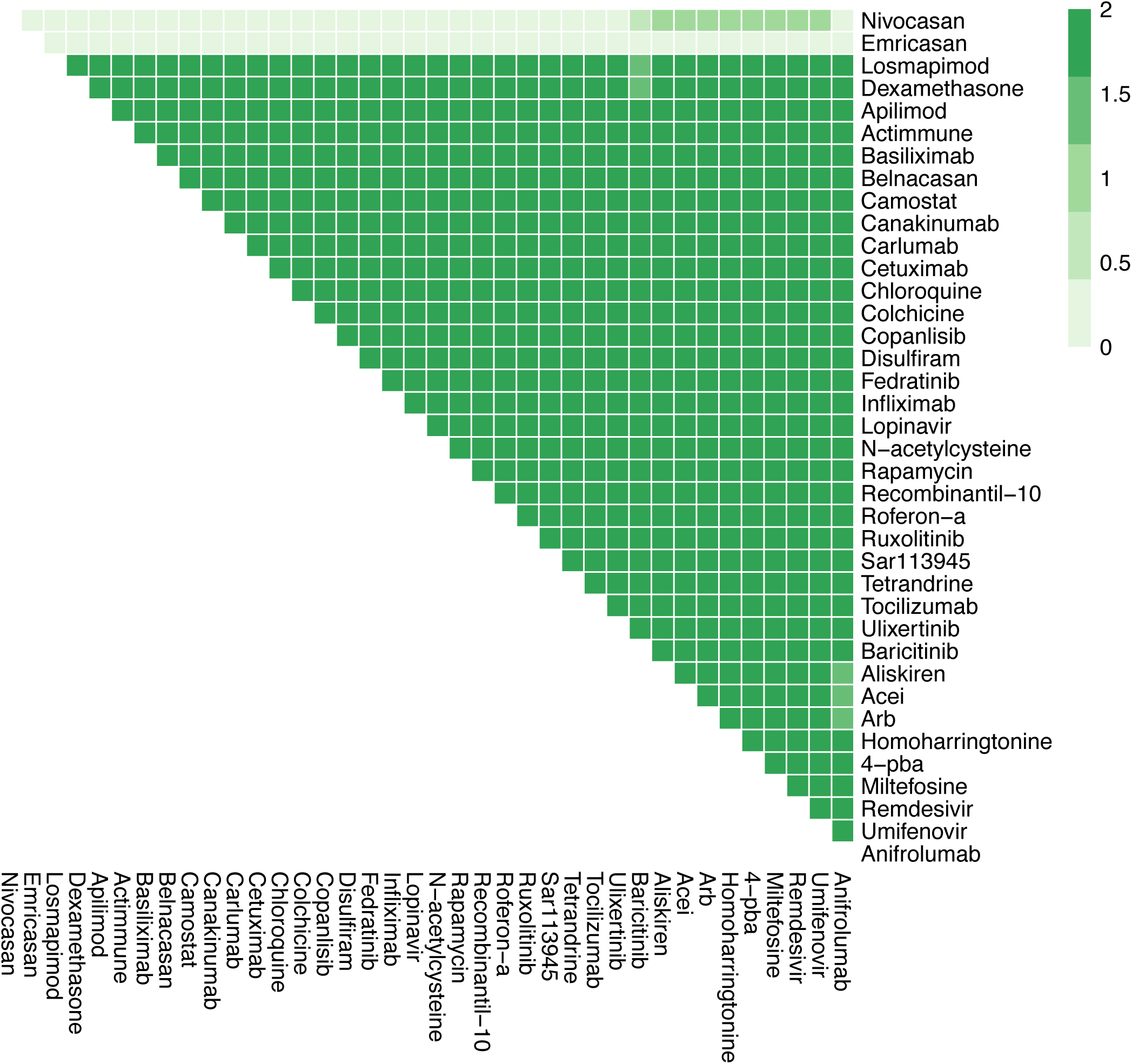
The effects of combination therapy on Cell Death in late stage, severe COVID-19. The colour of the square corresponds to the level of the *CellDeath* node at steady state under treatment by the drugs indicated on the x- and y-axis.

**Figure S4J.**
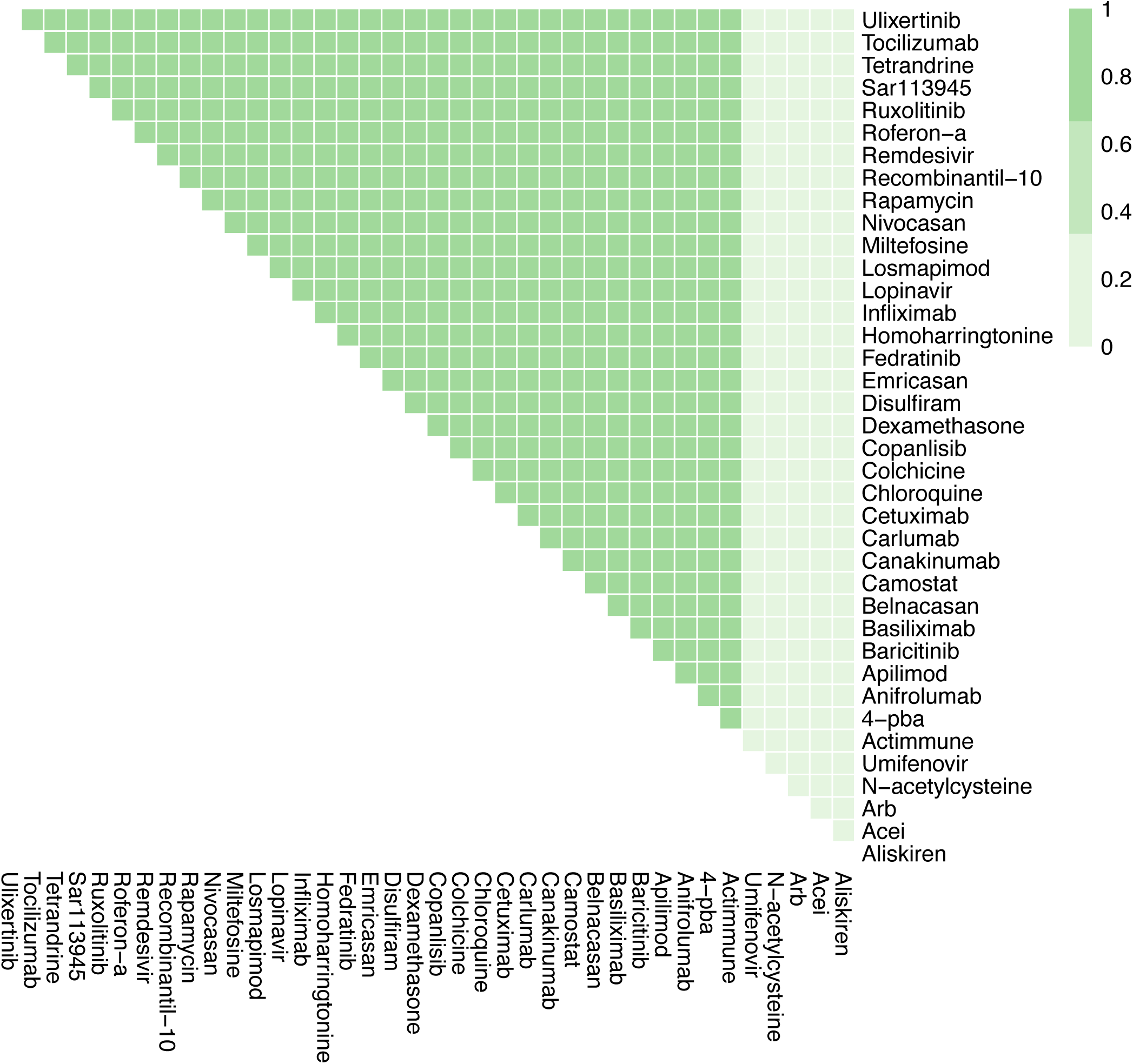
The effects of combination therapy on Fibrosis in late stage, severe COVID-19. The colour of the square corresponds to the level of the *Fibrosis* node at steady state under treatment by the drugs indicated on the x- and y-axis.

**Figure S4K.**
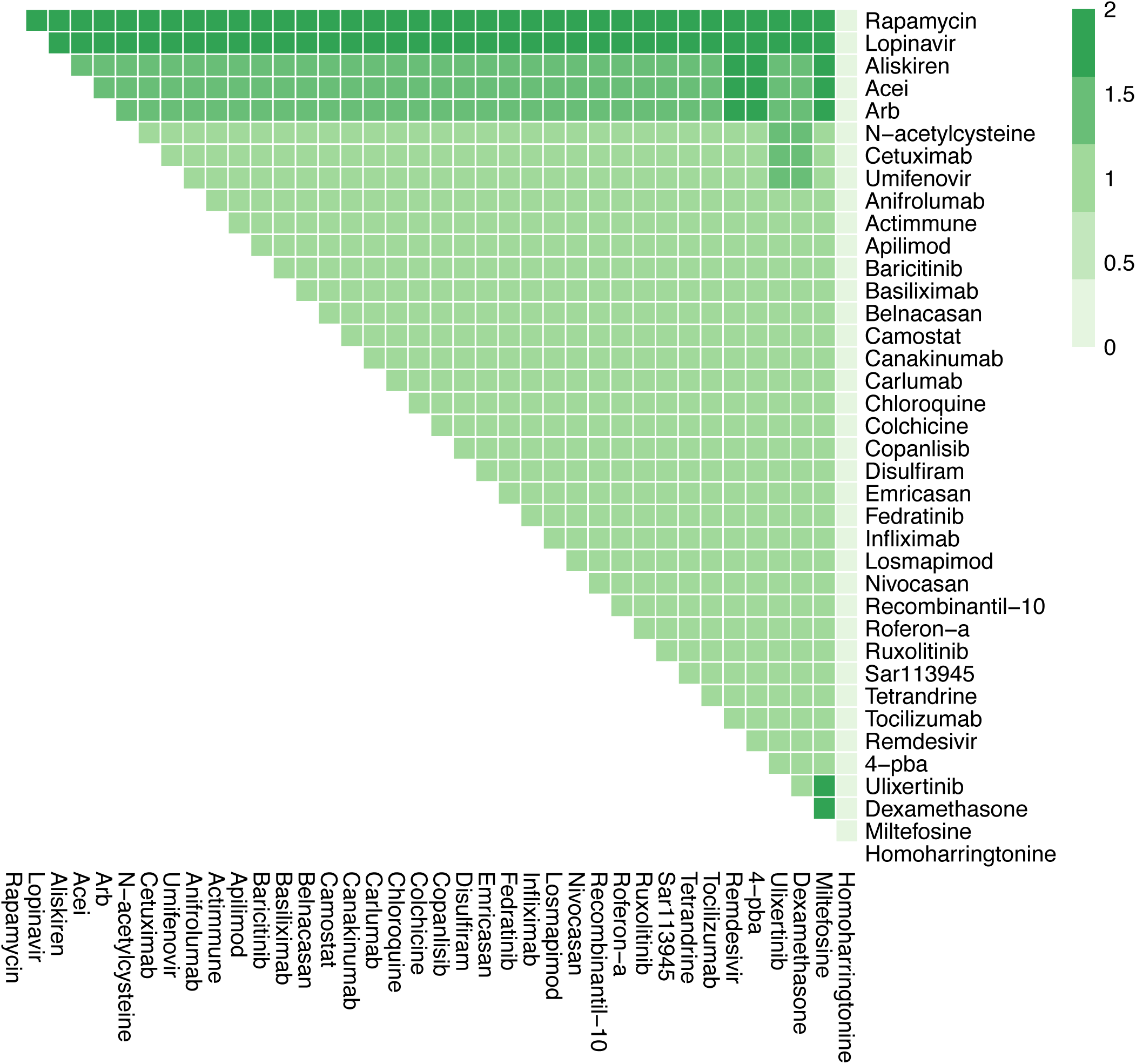
The effects of combination therapy on Host Protein Synthesis in late stage, severe COVID-19. The colour of the square corresponds to the level of the *HostProteinSynthesis* node at steady state under treatment by the drugs indicated on the x- and y-axis.

**Figure S4L.**
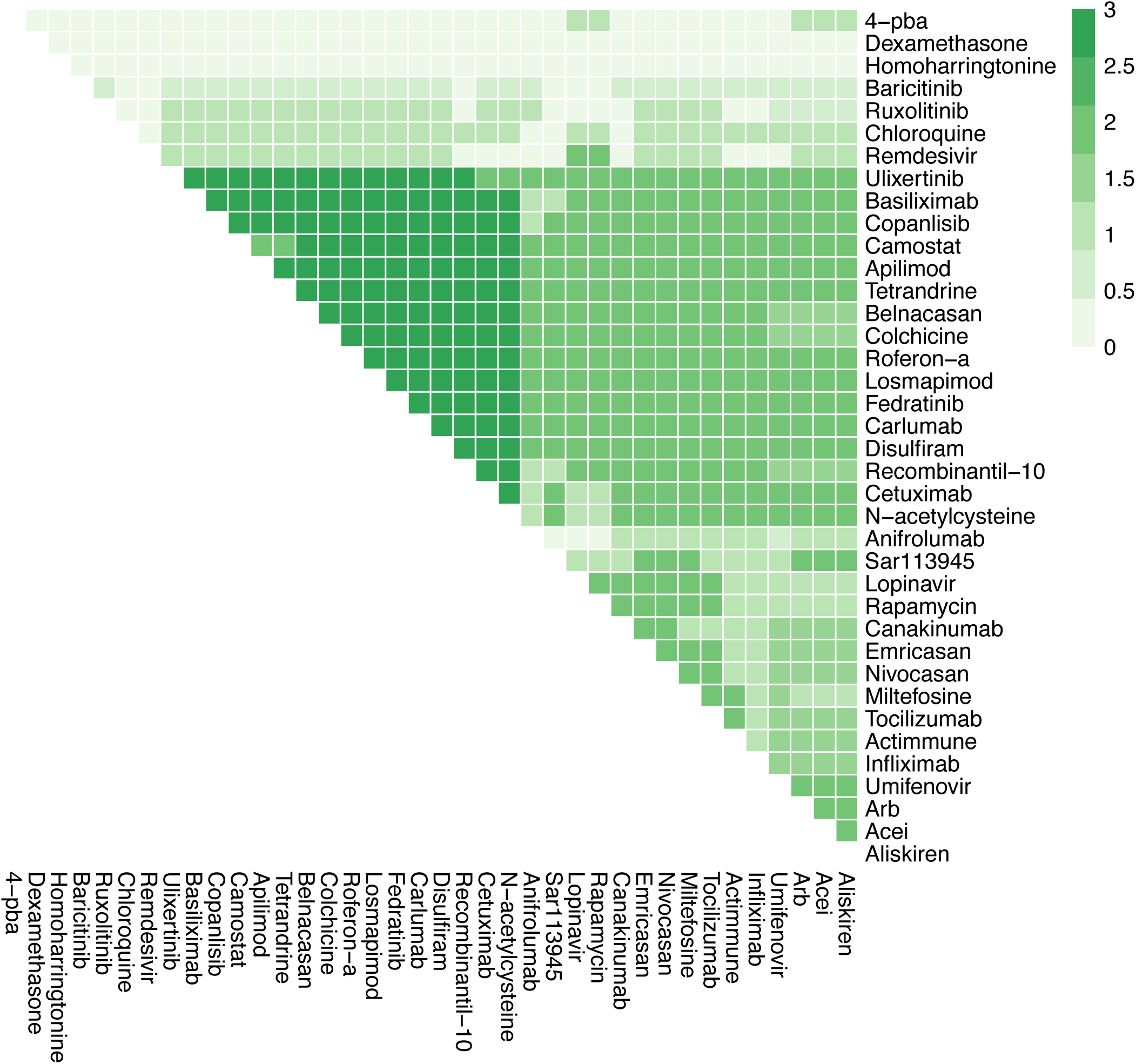
The effects of combination therapy on Inflammation in late stage, severe COVID-19. The colour of the square corresponds to the level of the *Inflammation* node at steady state under treatment by the drugs indicated on the x- and y-axis.

**Figure S4M.**
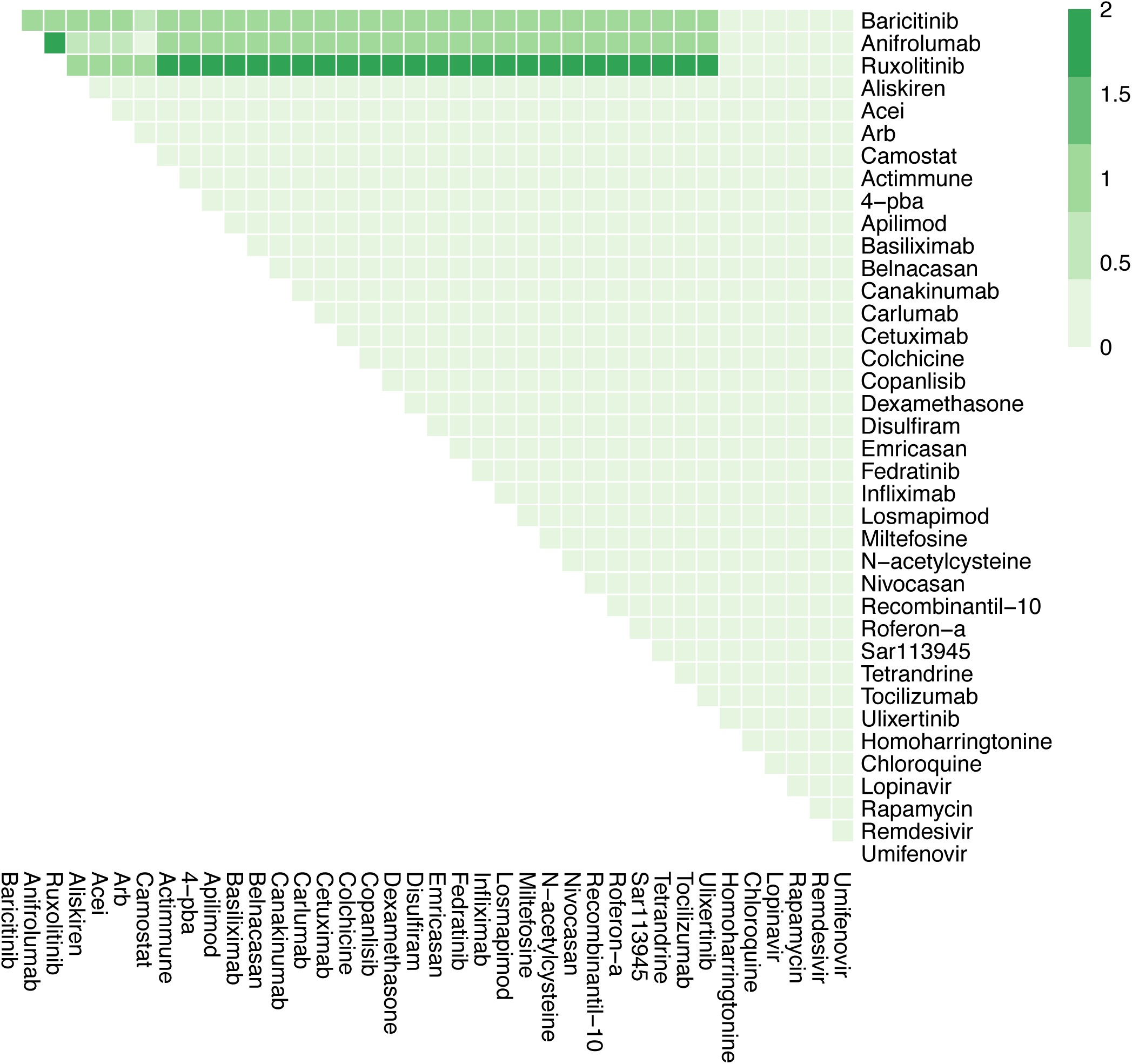
The effects of combination therapy on Syncytia Formation in late stage, severe COVID-19. The colour of the square corresponds to the level of the *SyncytiaFormation* node at steady state under treatment by the drugs indicated on the x- and y-axis.

**Figure S4N.**
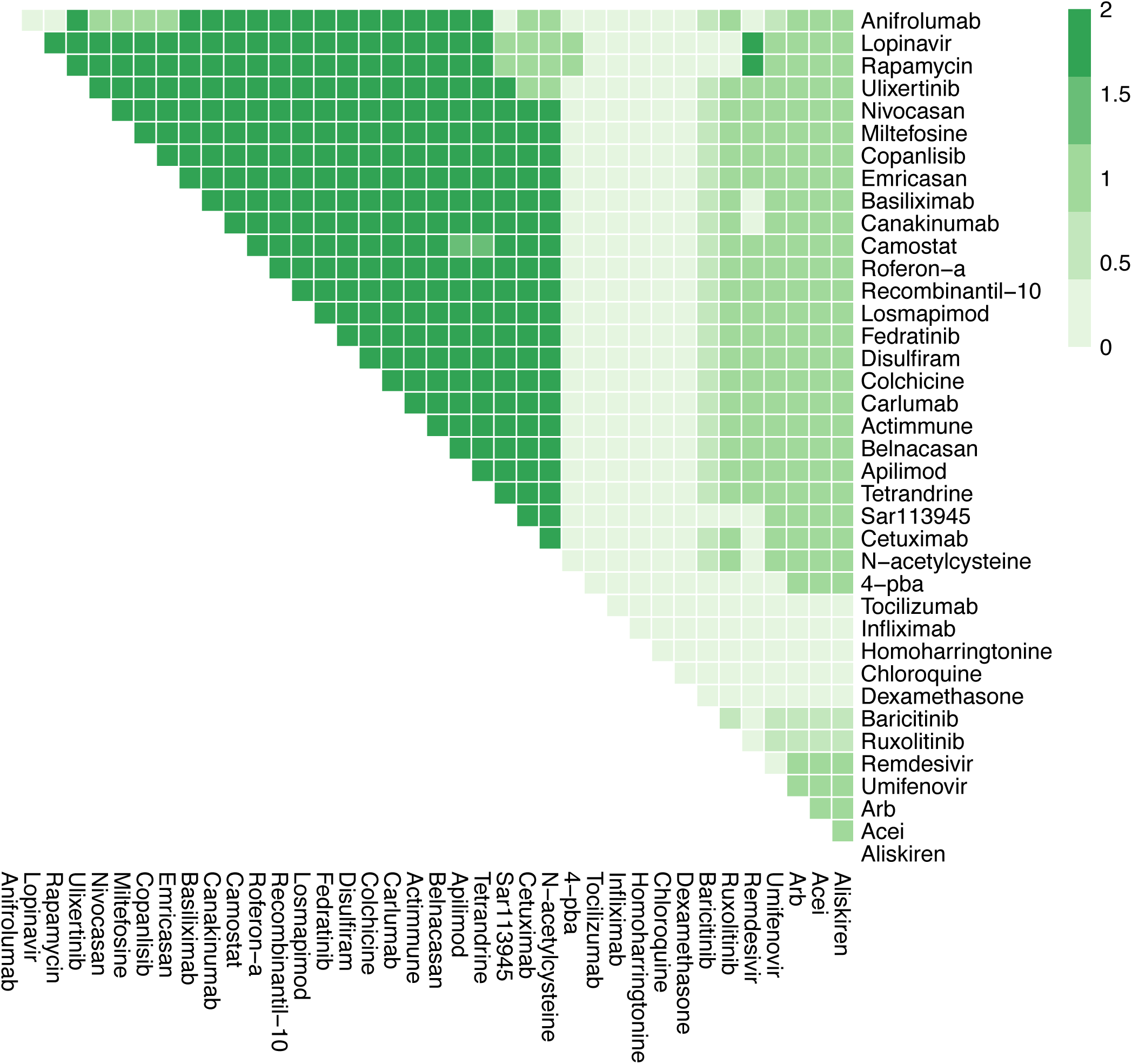
The effects of combination therapy on T-cell Infiltration in late stage, severe COVID-19. The colour of the square corresponds to the level of the *T-cellInfiltration* node at steady state under treatment by the drugs indicated on the x- and y-axis.

**Figure S4O.**
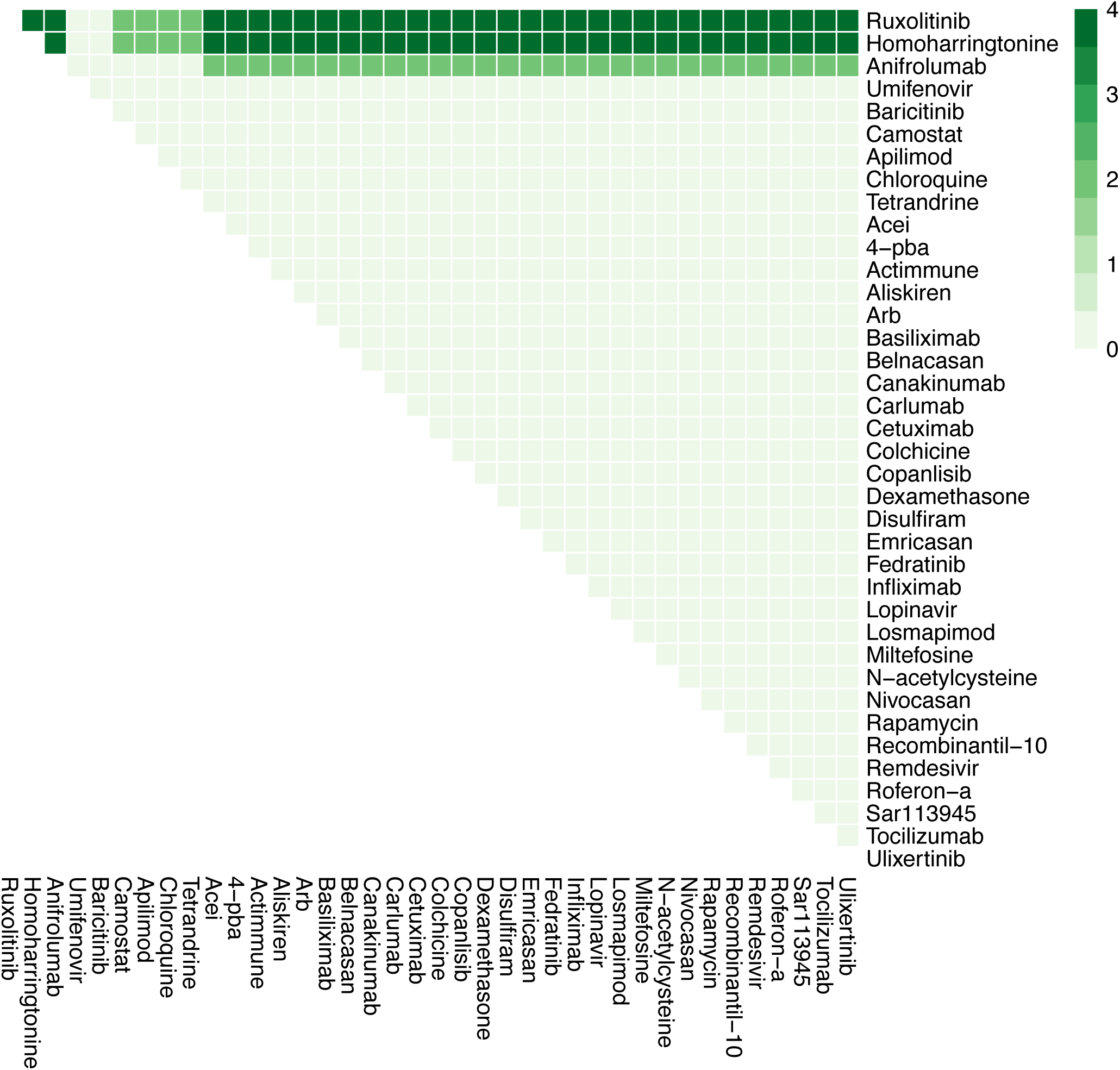
The effects of combination therapy on Viral Entry in late stage, severe COVID-19. The colour of the square corresponds to the level of the *ViralEntry* node at steady state under treatment by the drugs indicated on the x- and y-axis.

**Figure S4P.**
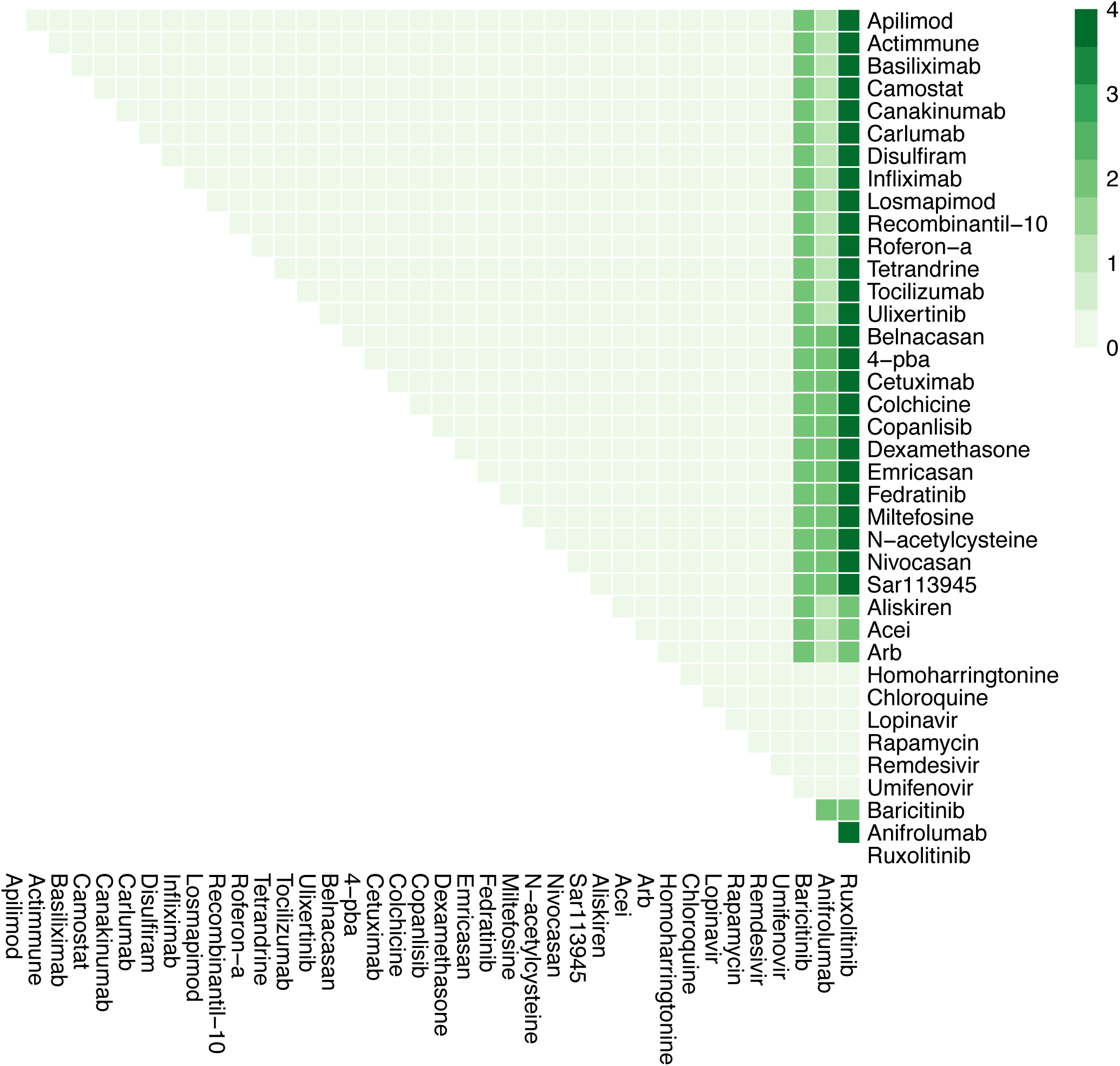
The effects of combination therapy on Viral Replication in late stage, severe COVID-19. The colour of the square corresponds to the level of the *Viral Replication* node at steady state under treatment by the drugs indicated on the x- and y-axis.

**Figure S5.**
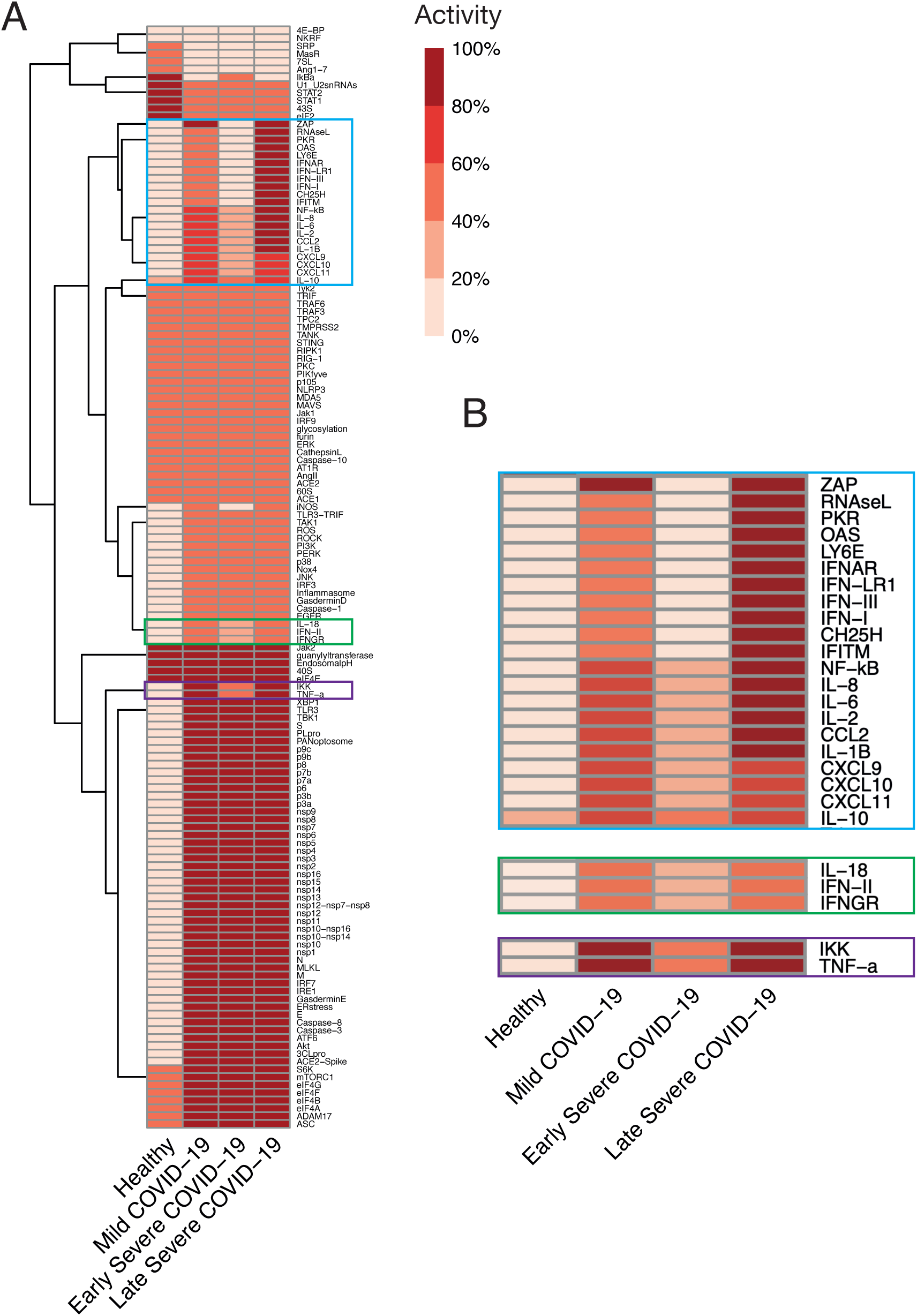
Differential characteristics of different states of COVID-19. (A) Hierarchical clustering heatmap representing the steady state values of each node in the network model for mild disease, and early and late severe disease (healthy controls for comparison). (B) Zoom in plot of areas in (A) that vary between disease states. All nodes normalised to maximal level of respective nodes, and range between 0-100%.

**Figure S6:**
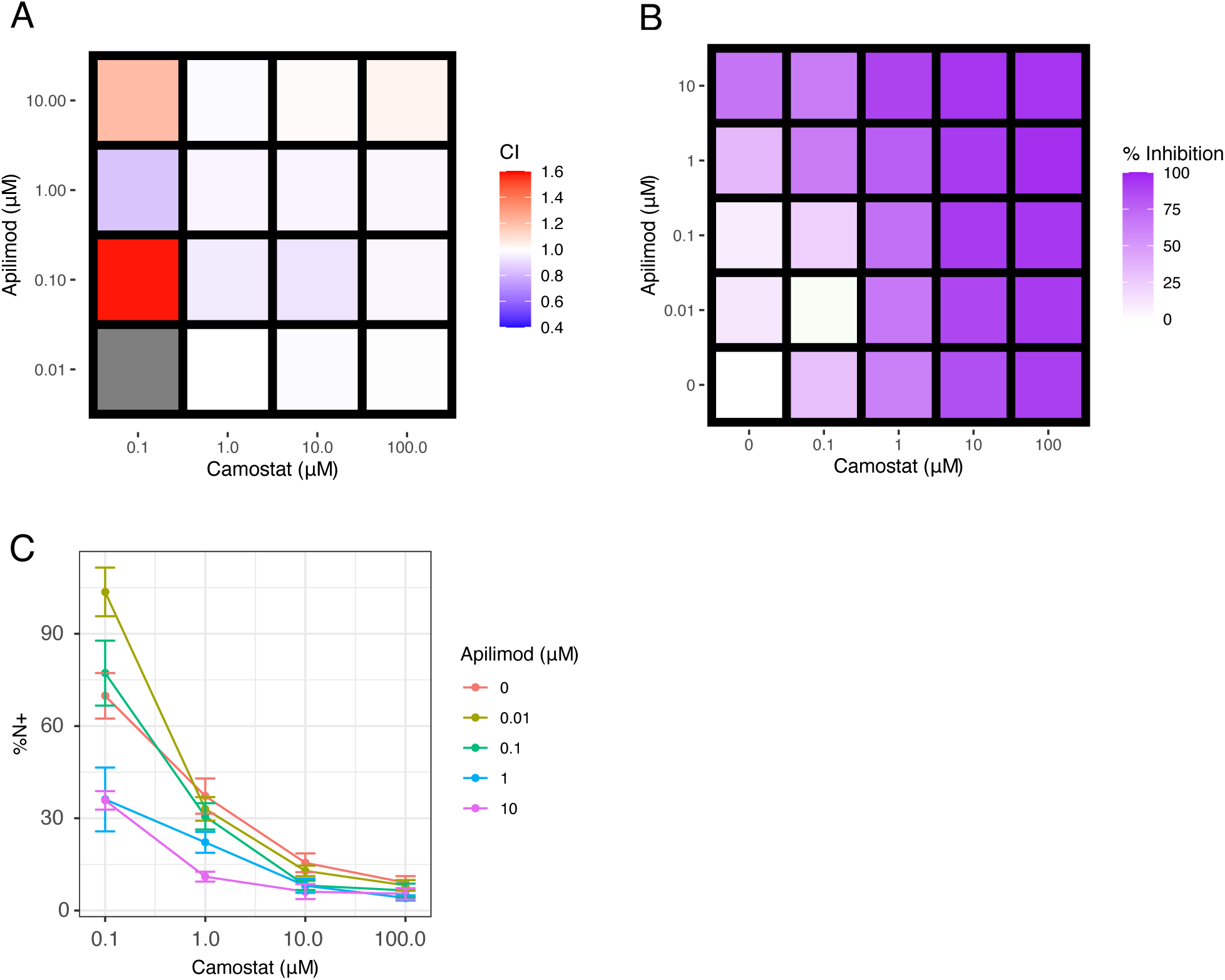
The impact of Camostat and Apilimod on replication of SARS-CoV-2 in Caco-2 cells. A) Heatmap showing the Cooperativity Index (CI) between each dose of Apilimod and Camostat tested. The CI was calculated according to the Bliss independence model. B) The percentage inhibition of cells expressing nucleocapsid protein achieved at each dose of Apilimod and Camostat tested. C) Line graph showing the effect of dose of Apilimod against increasing dose of Camostat. Data show the mean and standard error, from three independent experiments.

**Figure S7:**
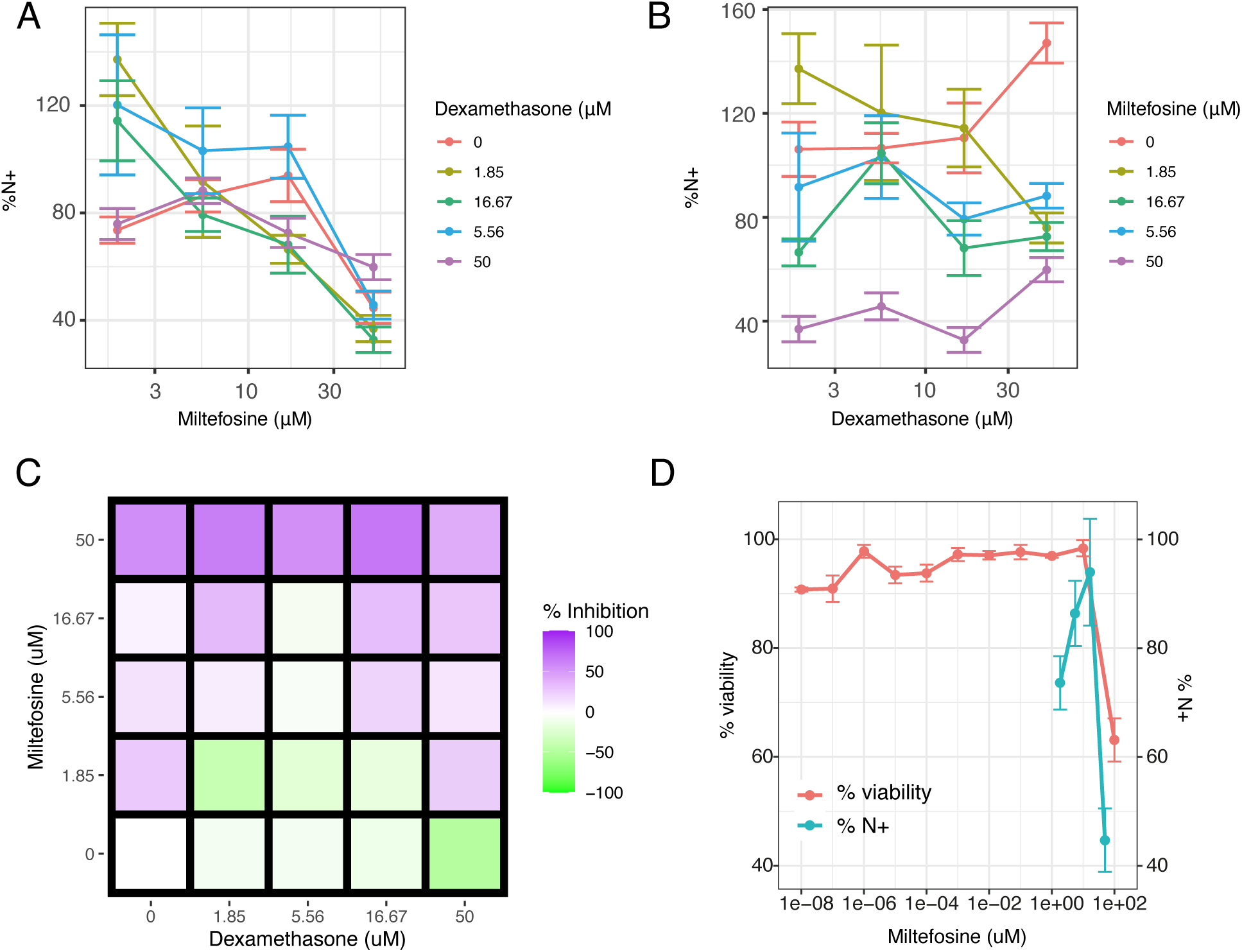
The impact of Dexamethasone and Miltefosine on replication of SARS-CoV-2 in Caco-2 cells. A) Line graph showing the effect of dose of Dexamethasone against increasing dose of Miltefosine. B) Line graph showing the effect of dose of Miltefosine against increasing dose of Dexamethasone. C) The percentage inhibition of cells expressing nucleocapsid protein achieved at each dose of Dexamethasone and Miltefosine tested. D) Percentage viability as determined by MTT assay plotted alongside percentage of cells expressing nucleocapsid protein at varying doses of Miltefosine. Data show the mean and standard error, from three independent experiments.

**Figure S8 A-H.**
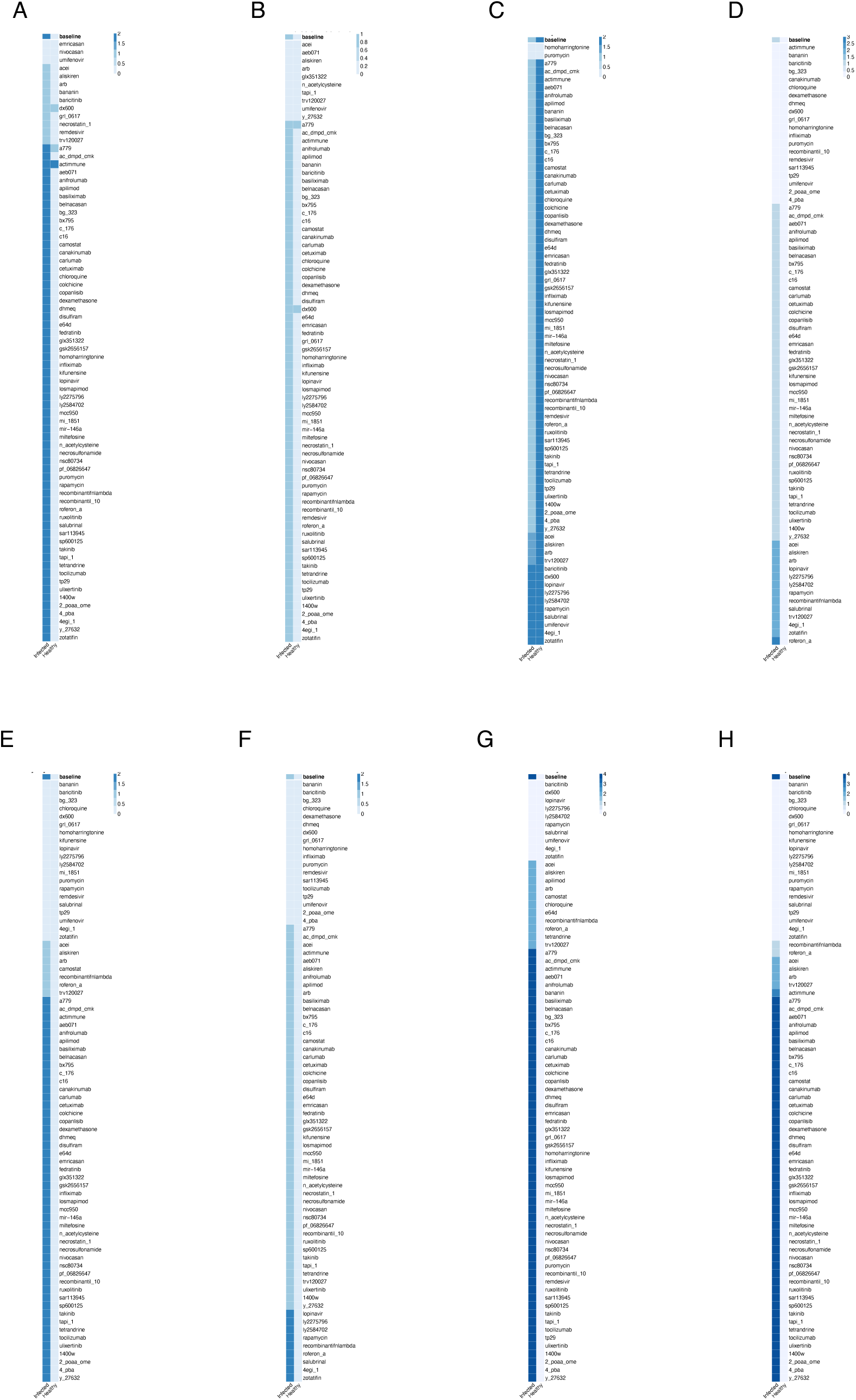
*In silico* screens to identify potential drugs to treat early severe COVID-19. Columns represent infected and healthy conditions, rows represent drug tested including those not yet approved for use. Showing the effect on: (A) Cell Death. (B) Fibrosis. (C) Host Protein Synthesis. (D) Inflammation. (E) Syncytia Formation. (F) T-cell Infiltration. (G) Viral Entry. (H) Viral Replication.

**Figure S8I.**
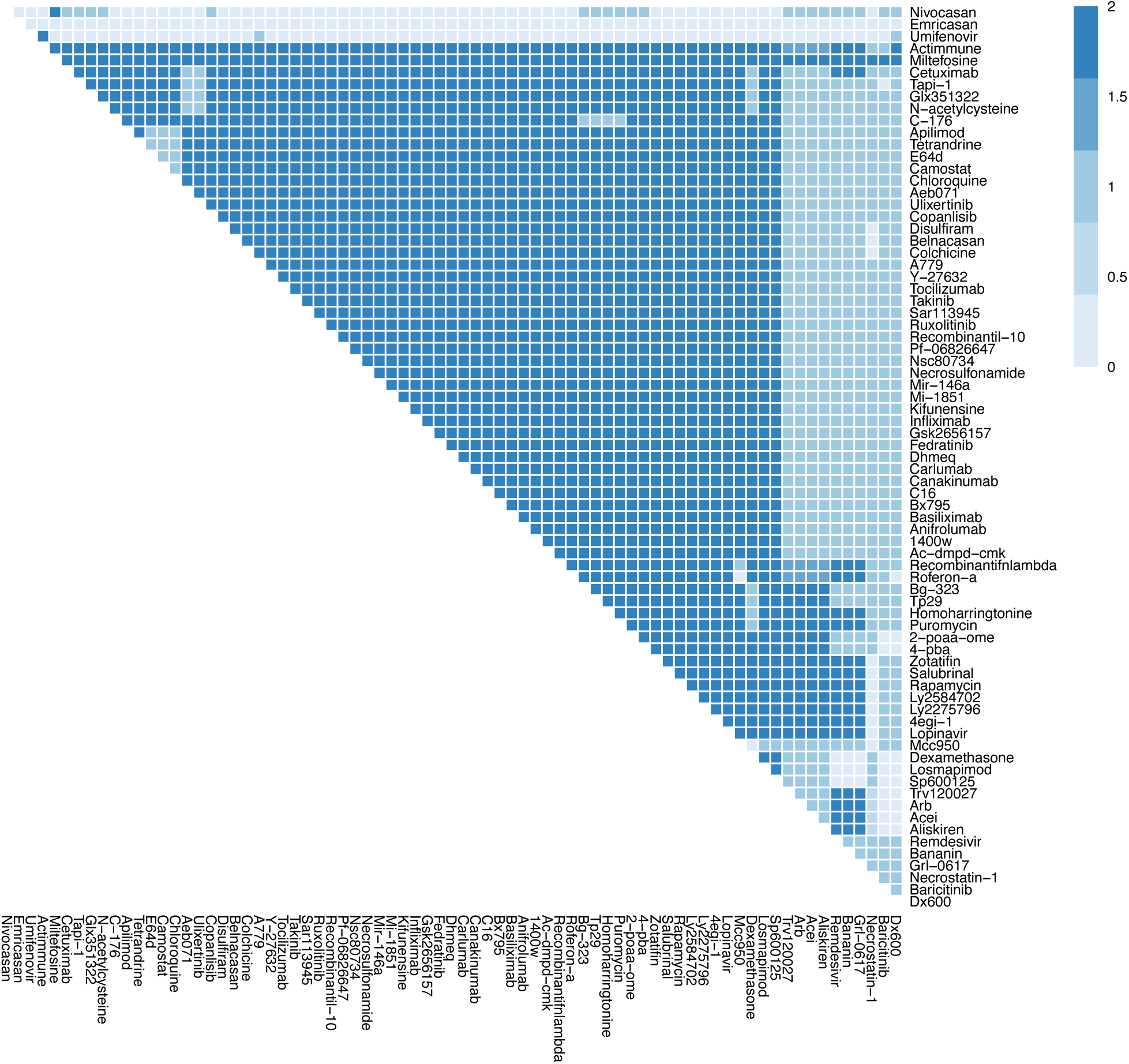
The effects of combination therapy on Cell Death in early stage, severe COVID-19. The colour of the square corresponds to the level of the *CellDeath* node at steady state under treatment by the drugs indicated on the x- and y-axis. This plot shows all drugs, including those at an experimental stage of development.

**Figure S8J.**
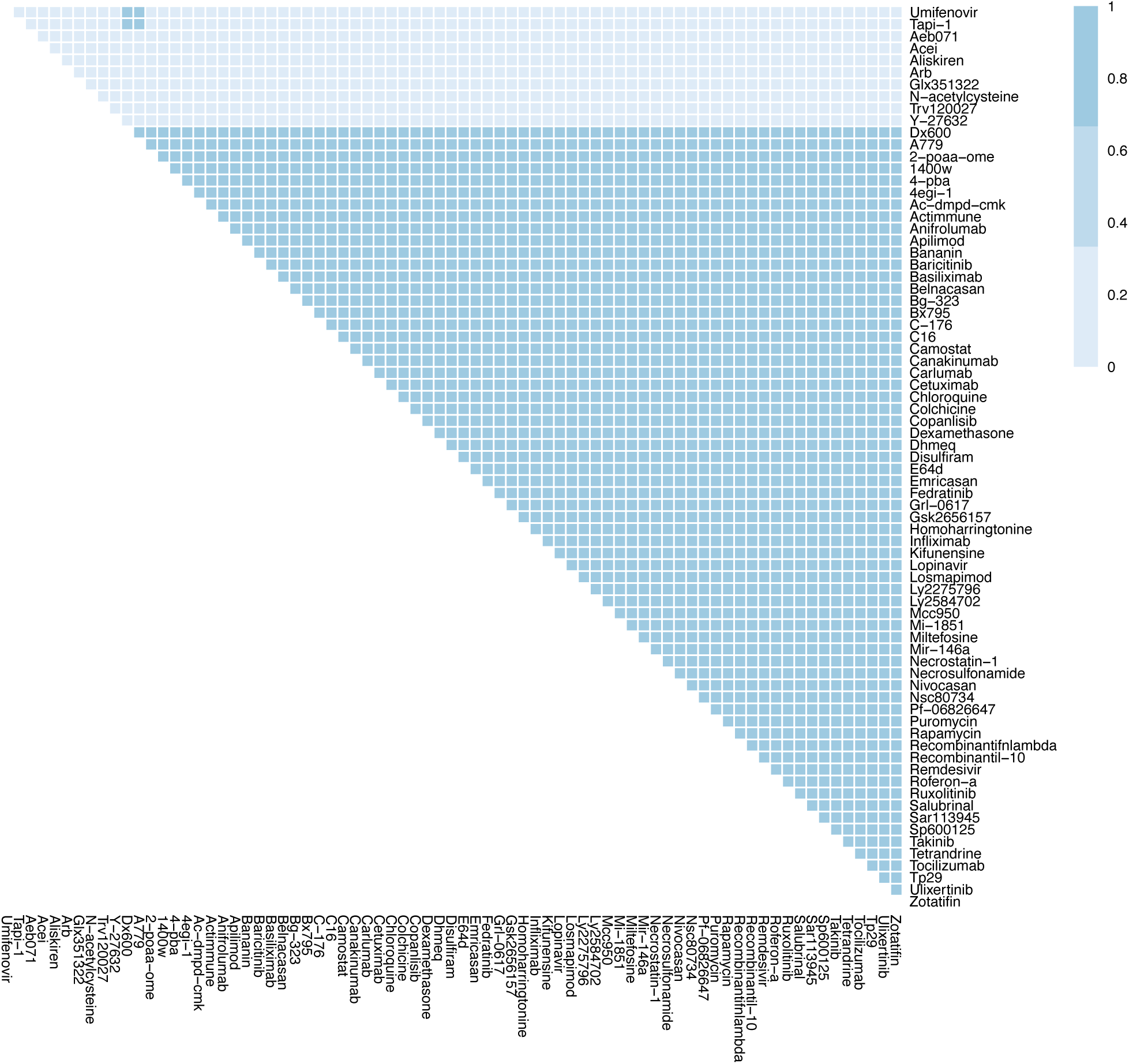
The effects of combination therapy on Fibrosis in early stage, severe COVID-19. The colour of the square corresponds to the level of the *Fibrosis* node at steady state under treatment by the drugs indicated on the x- and y-axis. This plot shows all drugs, including those at an experimental stage of development.

**Figure S8K.**
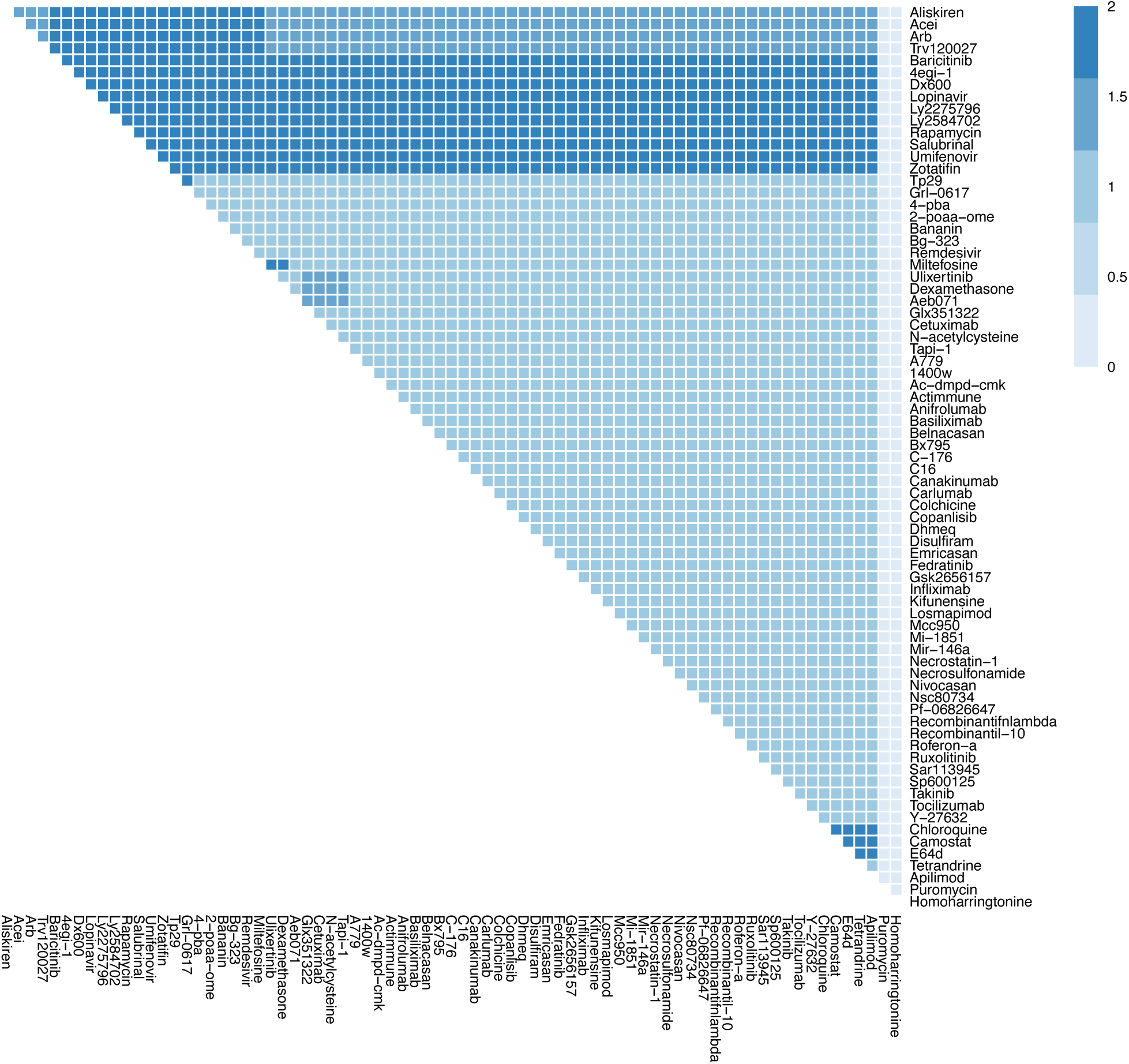
The effects of combination therapy on Host Protein Synthesis in early stage, severe COVID-19. The colour of the square corresponds to the level of the *HostProteinSynthesis* node at steady state under treatment by the drugs indicated on the x- and y-axis. This plot shows all drugs, including those at an experimental stage of development.

**Figure S8L.**
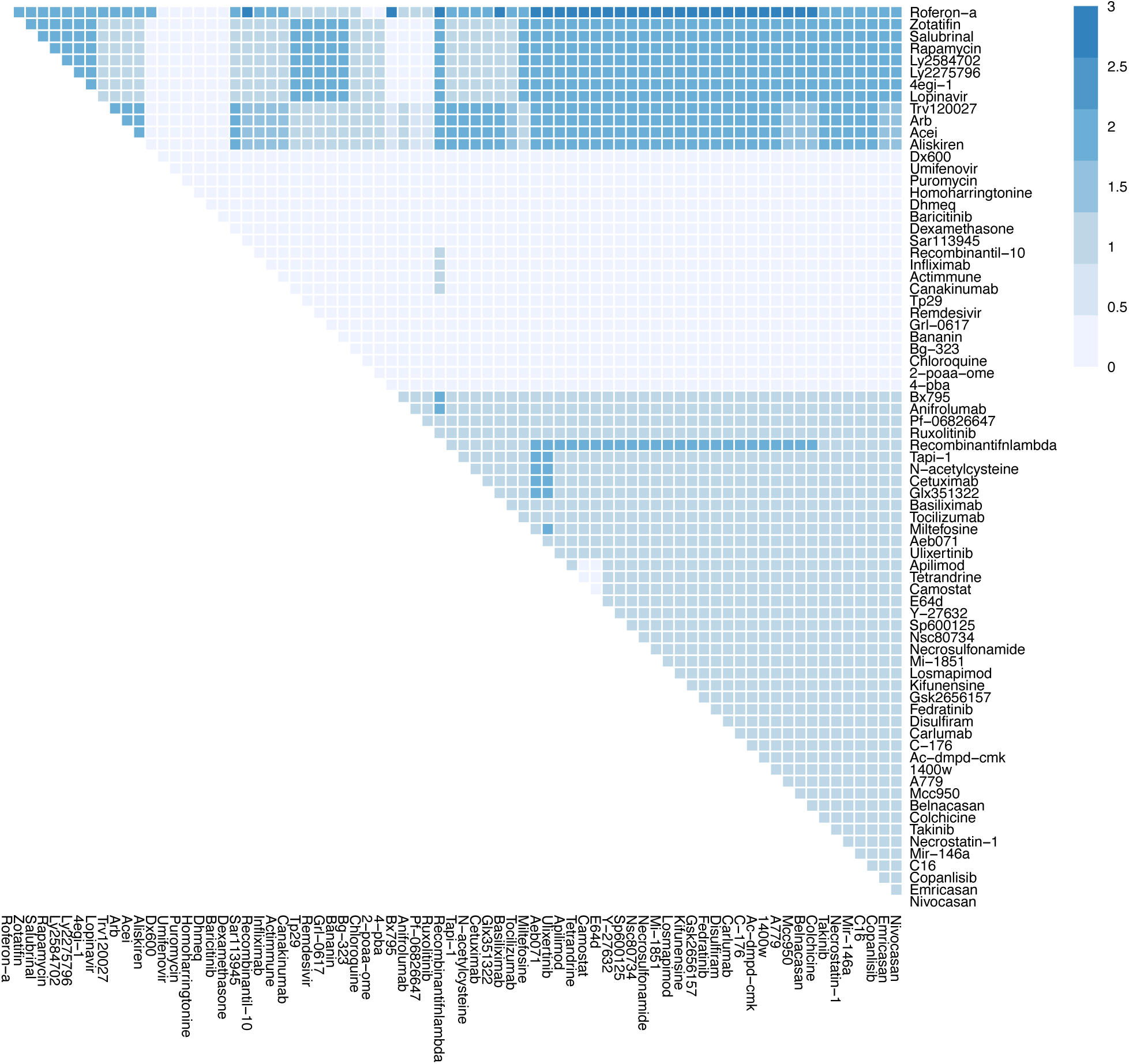
The effects of combination therapy on Inflammation in early stage, severe COVID-19. The colour of the square corresponds to the level of the *Inflammation* node at steady state under treatment by the drugs indicated on the x- and y-axis. This plot shows all drugs, including those at an experimental stage of development.

**Figure S8M.**
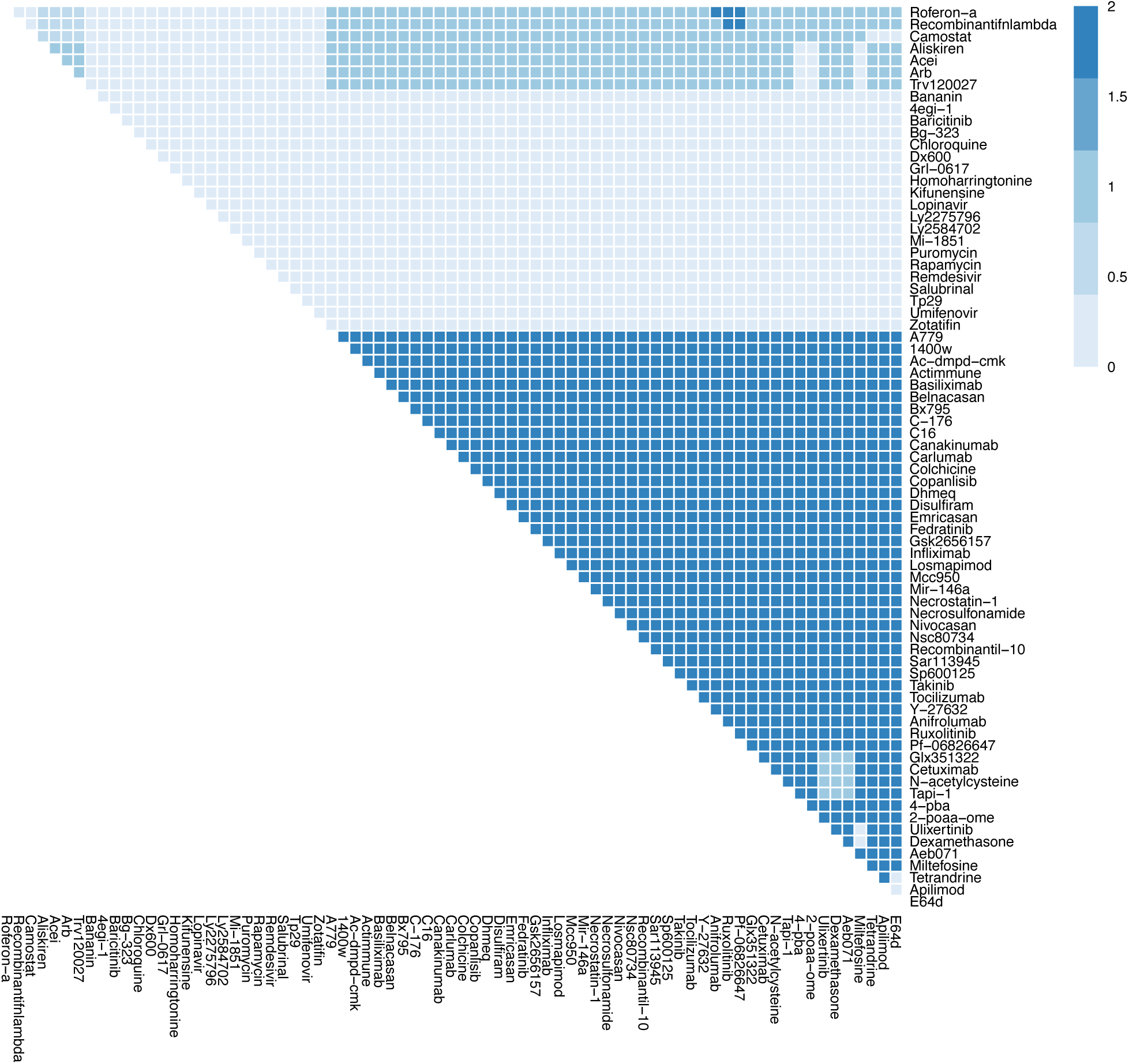
The effects of combination therapy on Syncytia Formation in early stage, severe COVID-19. The colour of the square corresponds to the level of the *SyncytiaFormation* node at steady state under treatment by the drugs indicated on the x- and y-axis. This plot shows all drugs, including those at an experimental stage of development.

**Figure S8N.**
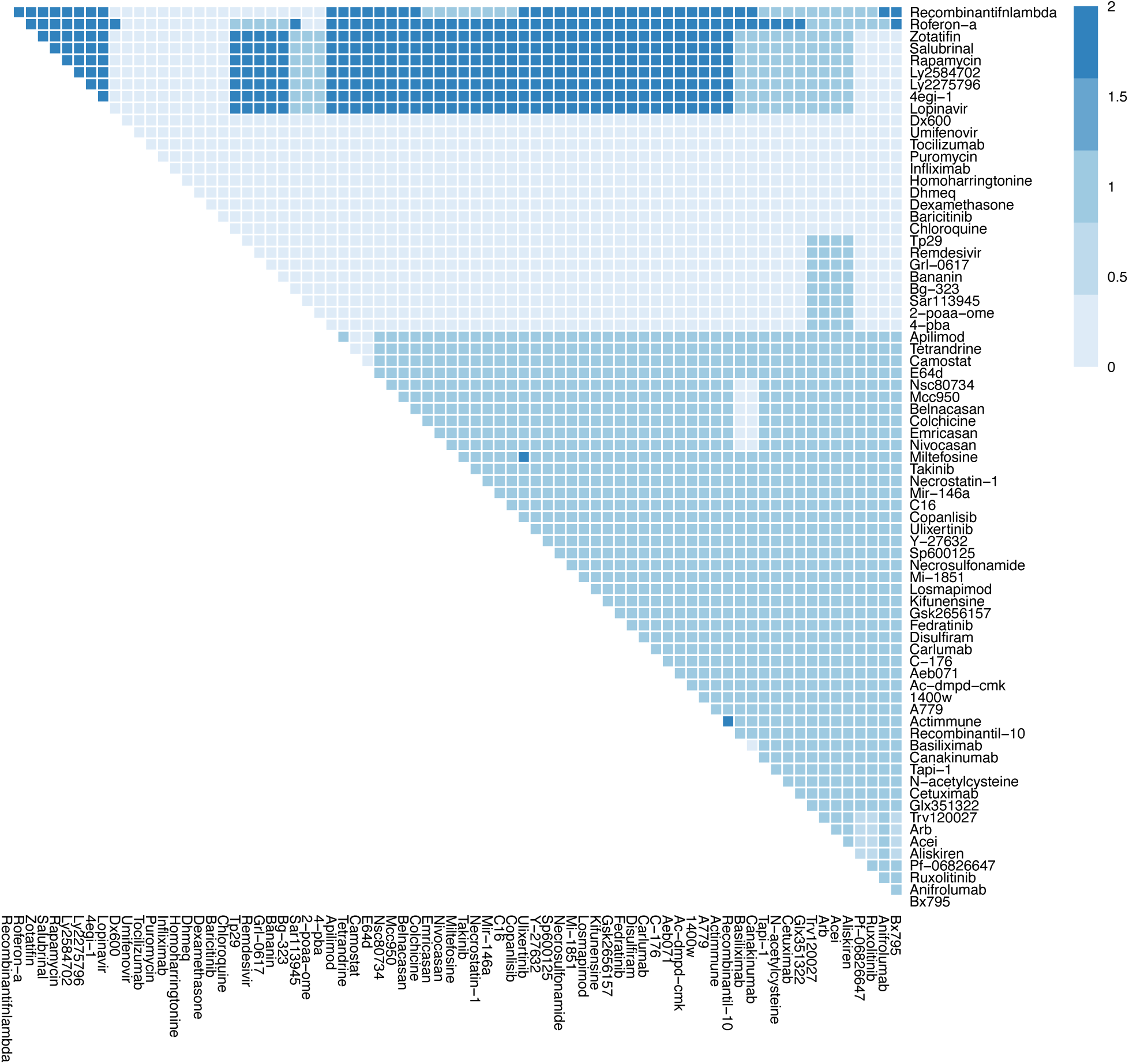
The effects of combination therapy on T-cell Infiltration in early stage, severe COVID-19. The colour of the square corresponds to the level of the *T-cellInfiltration* node at steady state under treatment by the drugs indicated on the x- and y-axis. This plot shows all drugs, including those at an experimental stage of development.

**Figure S8O.**
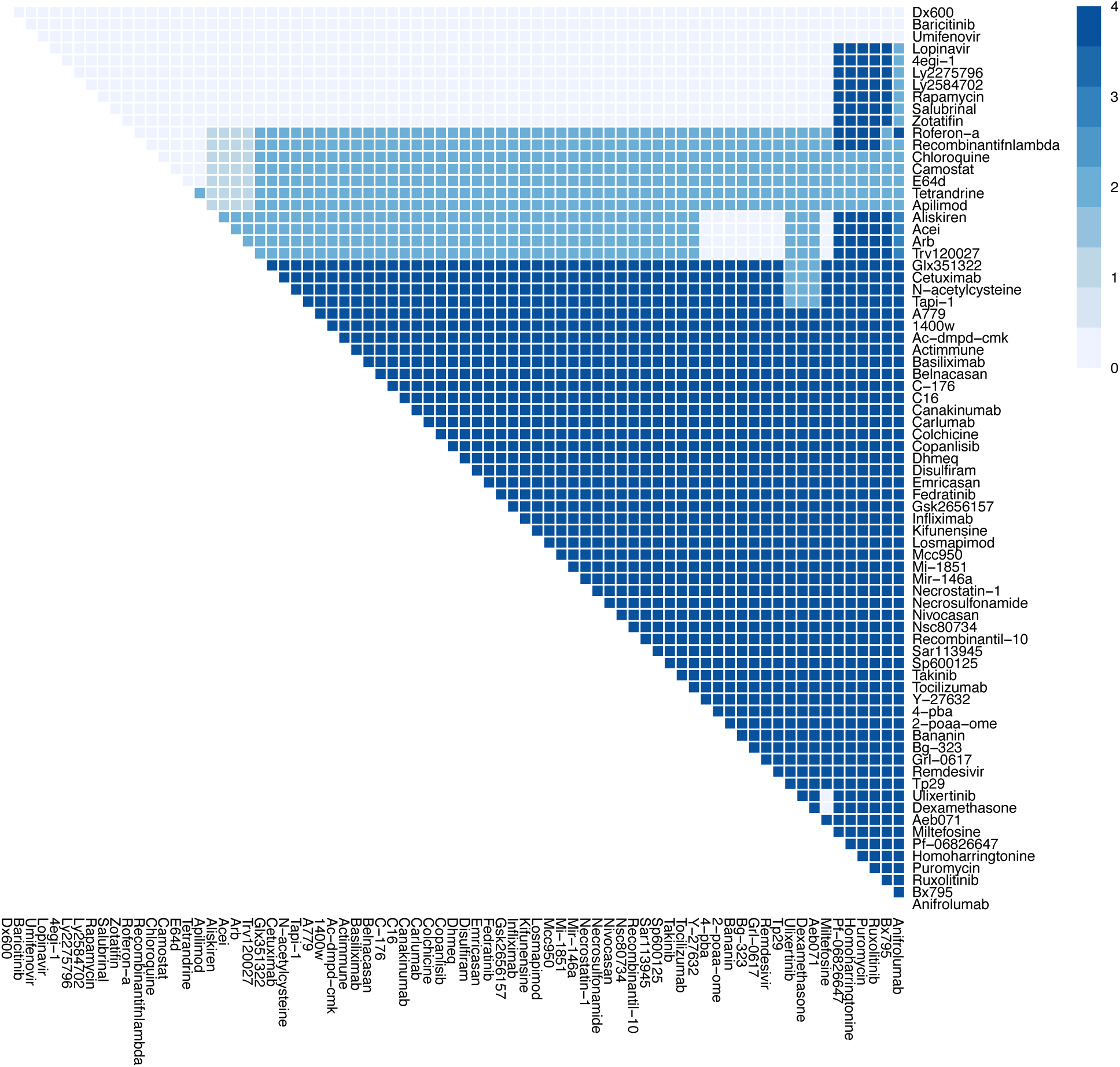
The effects of combination therapy on Viral Entry in early stage, severe COVID-19. The colour of the square corresponds to the level of the *ViralEntry* node at steady state under treatment by the drugs indicated on the x- and y-axis. This plot shows all drugs, including those at an experimental stage of development.

**Figure S8P.**
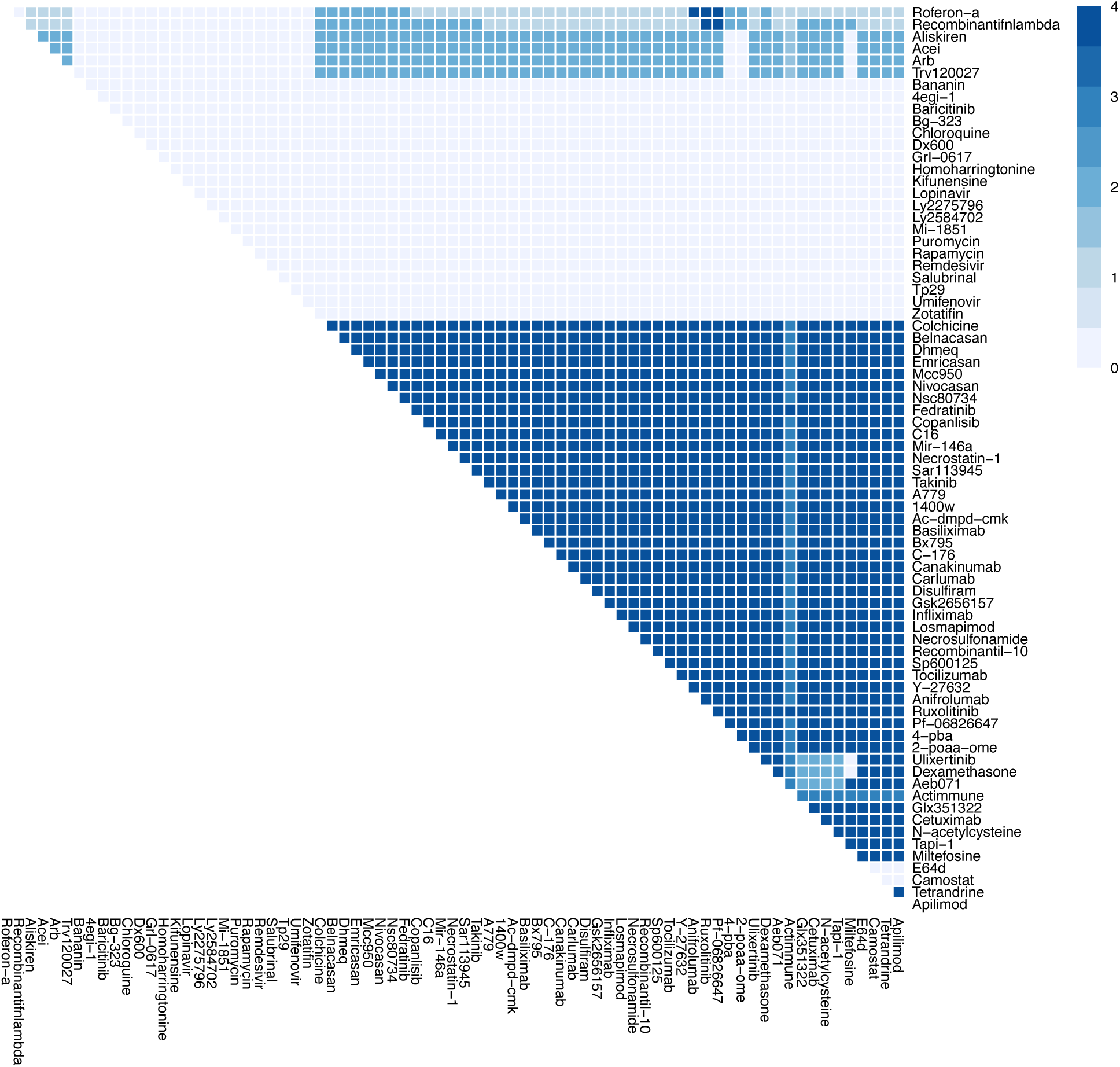
The effects of combination therapy on Viral Replication in early stage, severe COVID-19. The colour of the square corresponds to the level of the *ViralReplication* node at steady state under treatment by the drugs indicated on the x- and y-axis. This plot shows all drugs, including those at an experimental stage of development.

**Figure S9 A-H.**
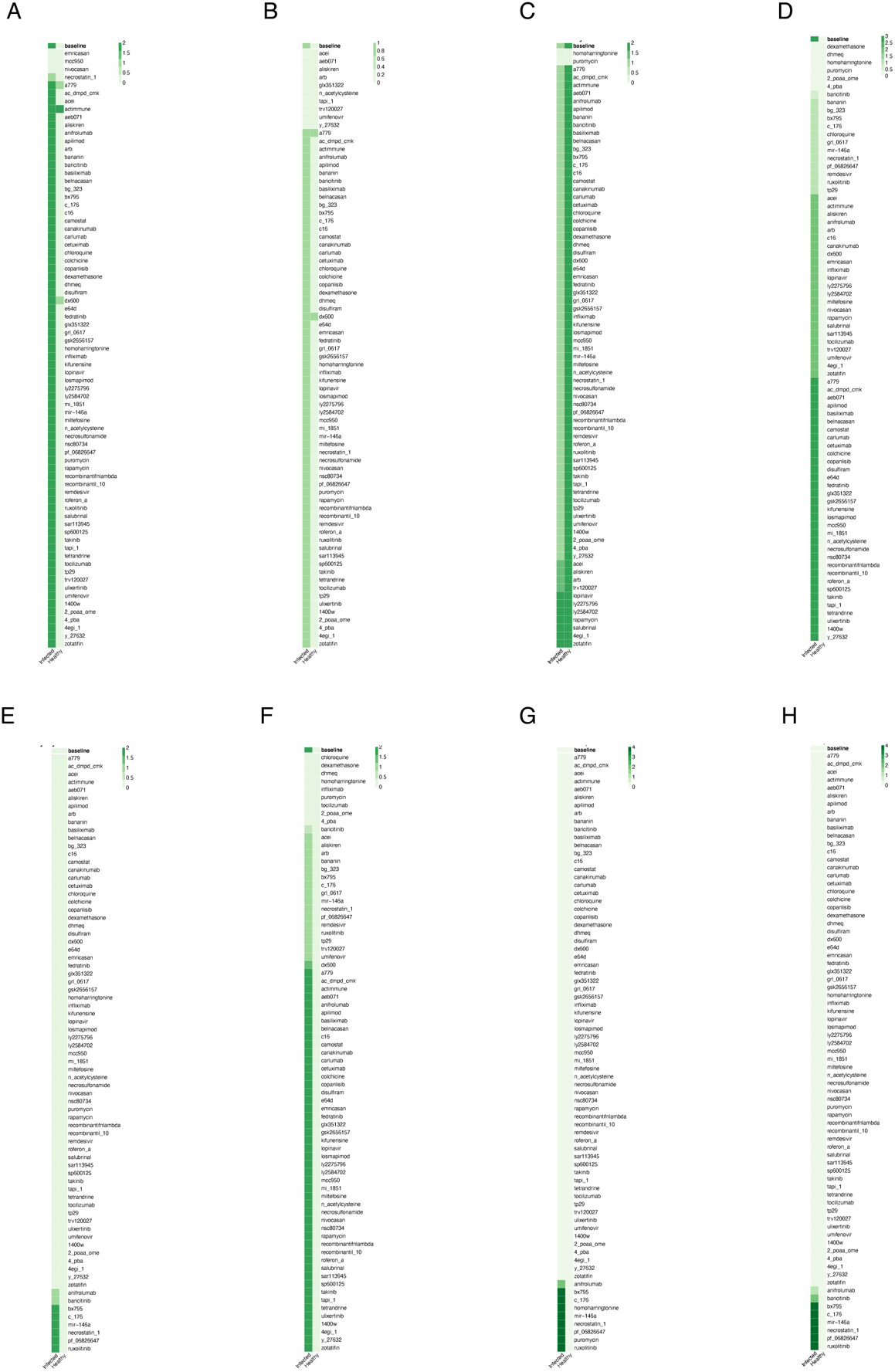
*In silico* screens to identify potential drugs to treat late severe COVID-19. Columns represent infected and healthy conditions, rows represent drug tested including those not yet approved for use. Showing the effect on: (A) Cell Death. (B) Fibrosis. (C) Host Protein Synthesis. (D) Inflammation. (E) Syncytia Formation. (F) T-cell Infiltration. (G) Viral Entry. (H) Viral Replication.

**Figure S9I.**
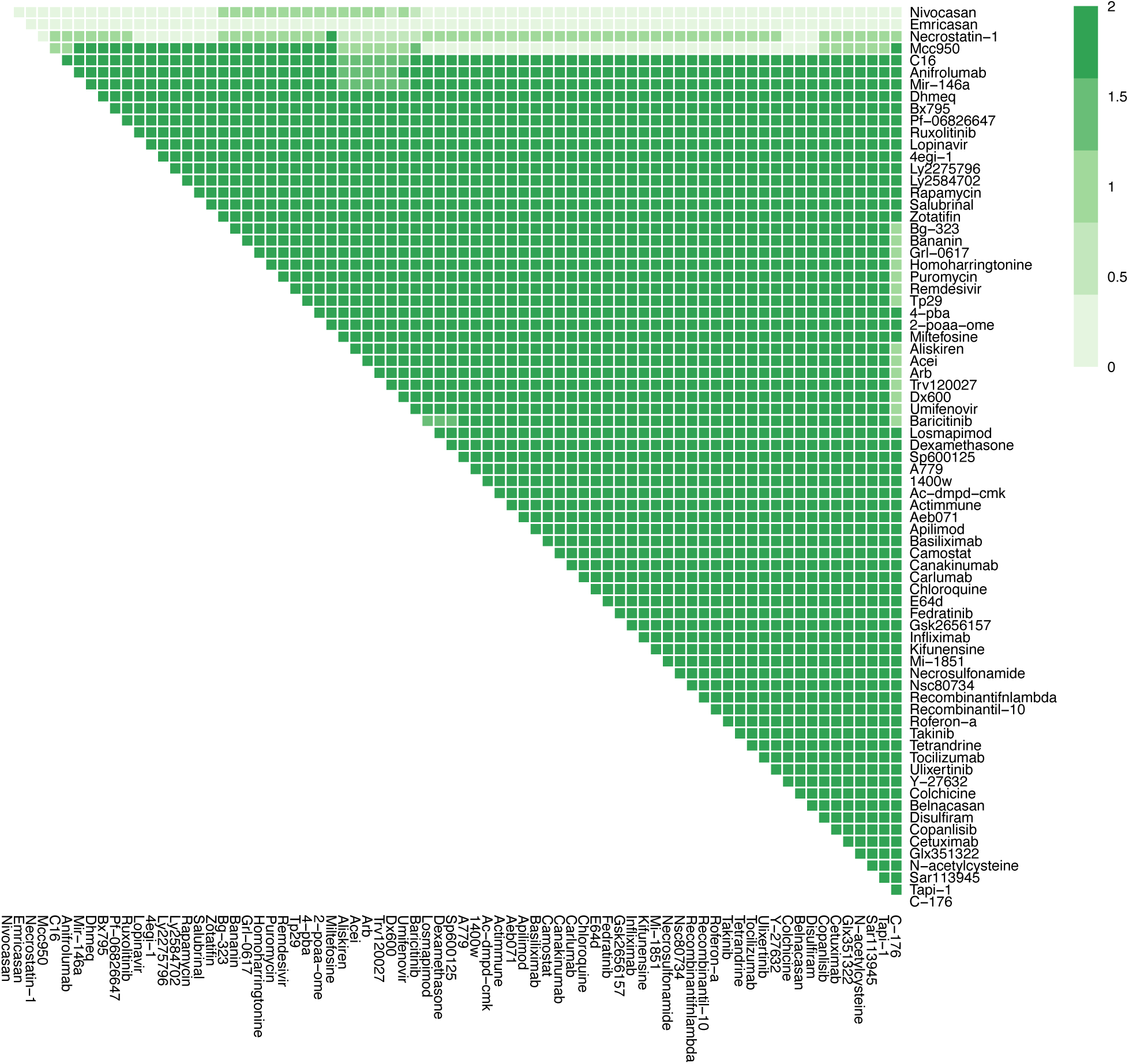
The effects of combination therapy on Cell Death in late stage, severe COVID-19. The colour of the square corresponds to the level of the *CellDeath* node at steady state under treatment by the drugs indicated on the x- and y-axis. This plot shows all drugs, including those at an experimental stage of development.

**Figure S9J.**
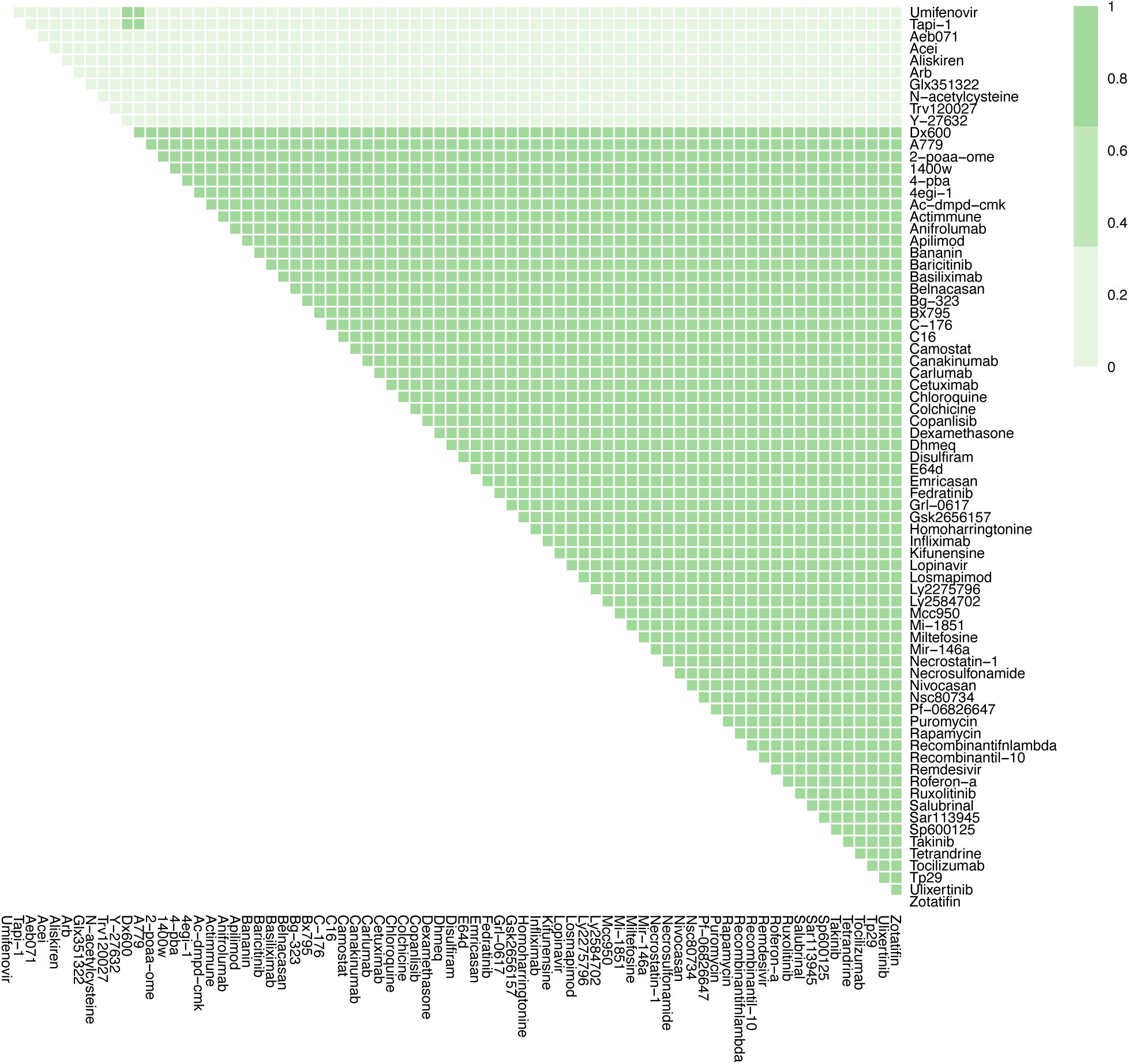
The effects of combination therapy on Fibrosis in late stage, severe COVID-19. The colour of the square corresponds to the level of the *Fibrosis* node at steady state under treatment by the drugs indicated on the x- and y-axis. This plot shows all drugs, including those at an experimental stage of development.

**Figure S9K.**
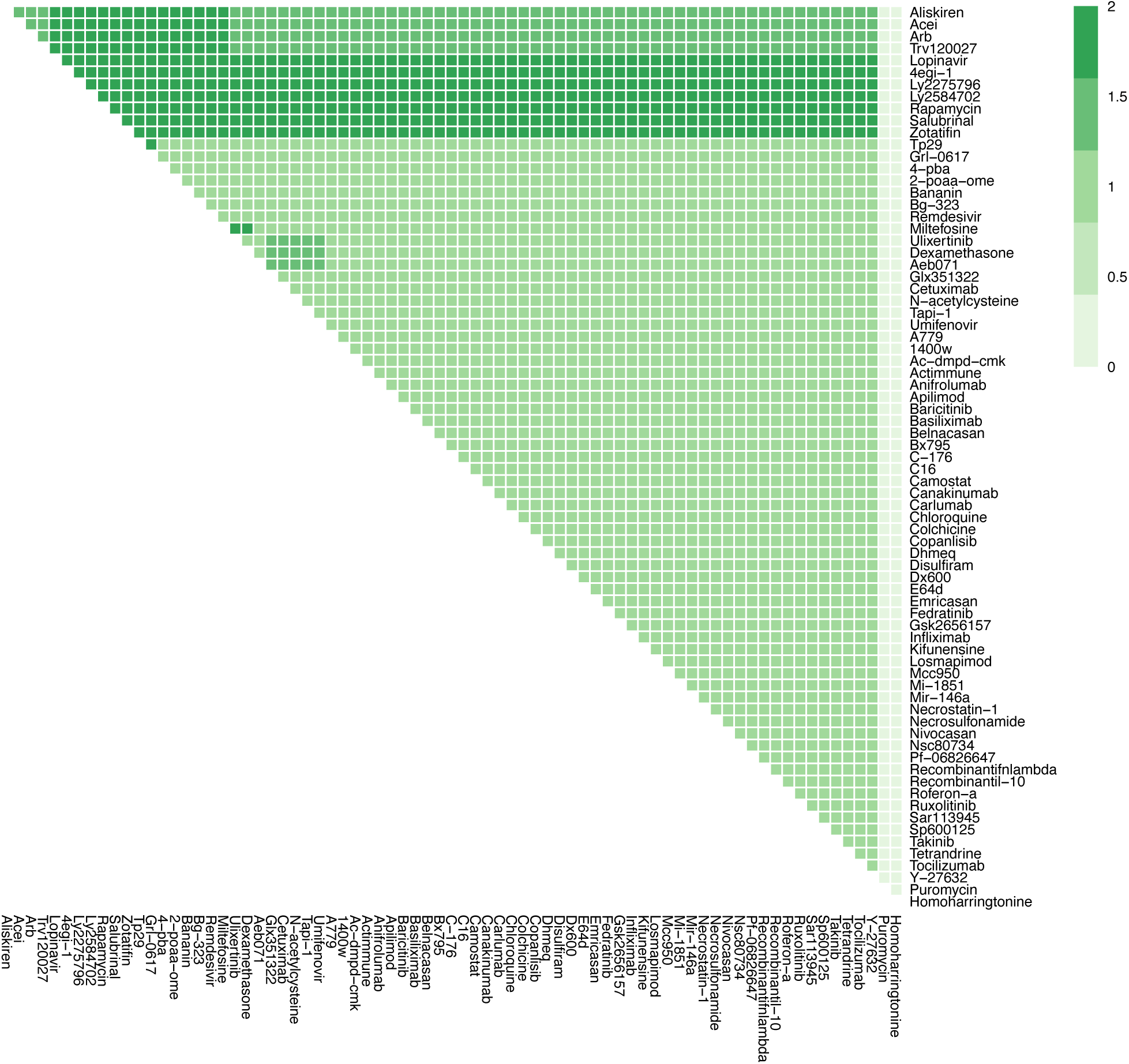
The effects of combination therapy on Host Protein Synthesis in late stage, severe COVID-19. The colour of the square corresponds to the level of the *HostProteinSynthesis* node at steady state under treatment by the drugs indicated on the x- and y-axis. This plot shows all drugs, including those at an experimental stage of development.

**Figure S9L.**
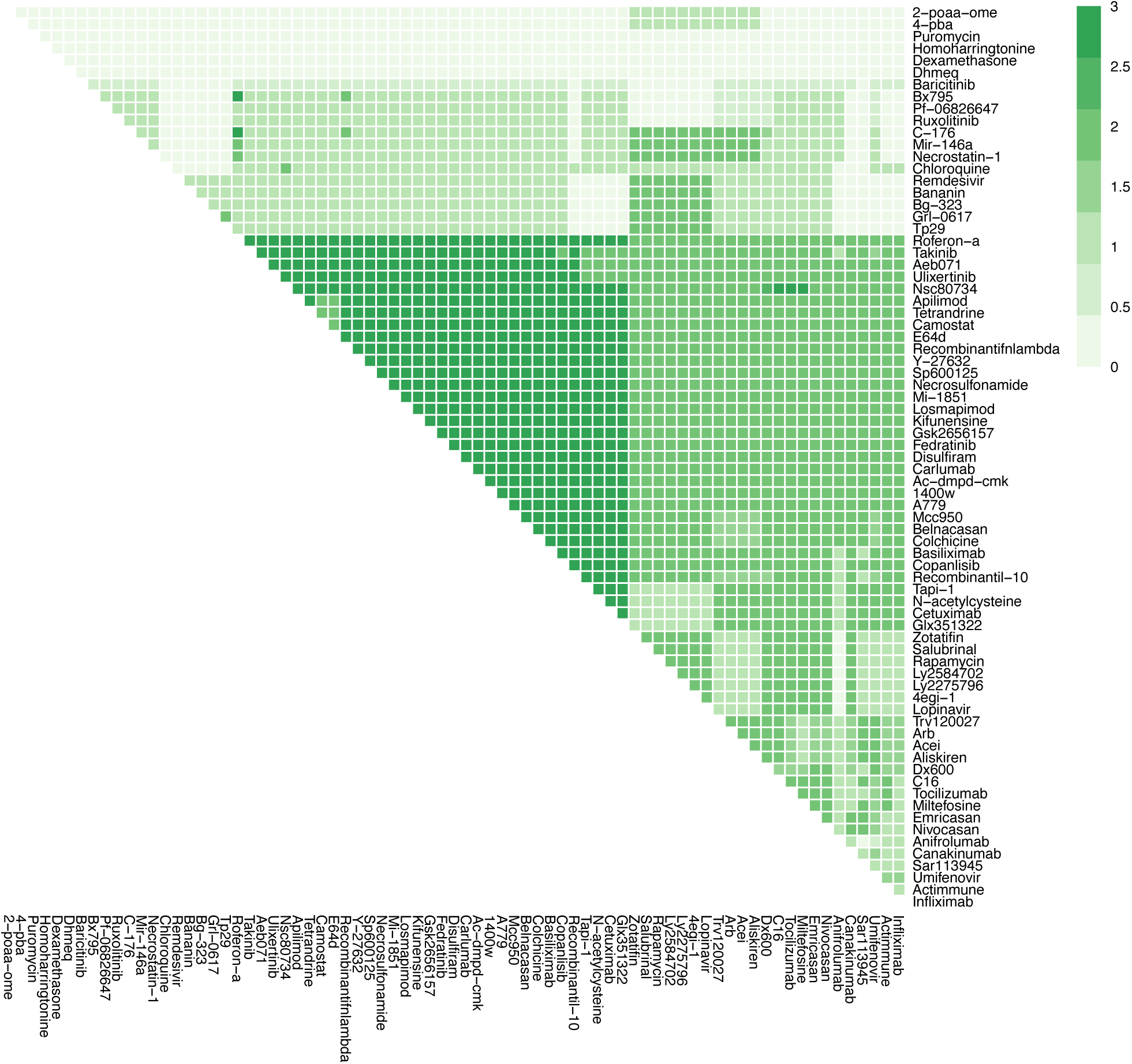
The effects of combination therapy on Inflammation in late stage, severe COVID-19. The colour of the square corresponds to the level of the *Inflammation* node at steady state under treatment by the drugs indicated on the x- and y-axis. This plot shows all drugs, including those at an experimental stage of development.

**Figure S9M.**
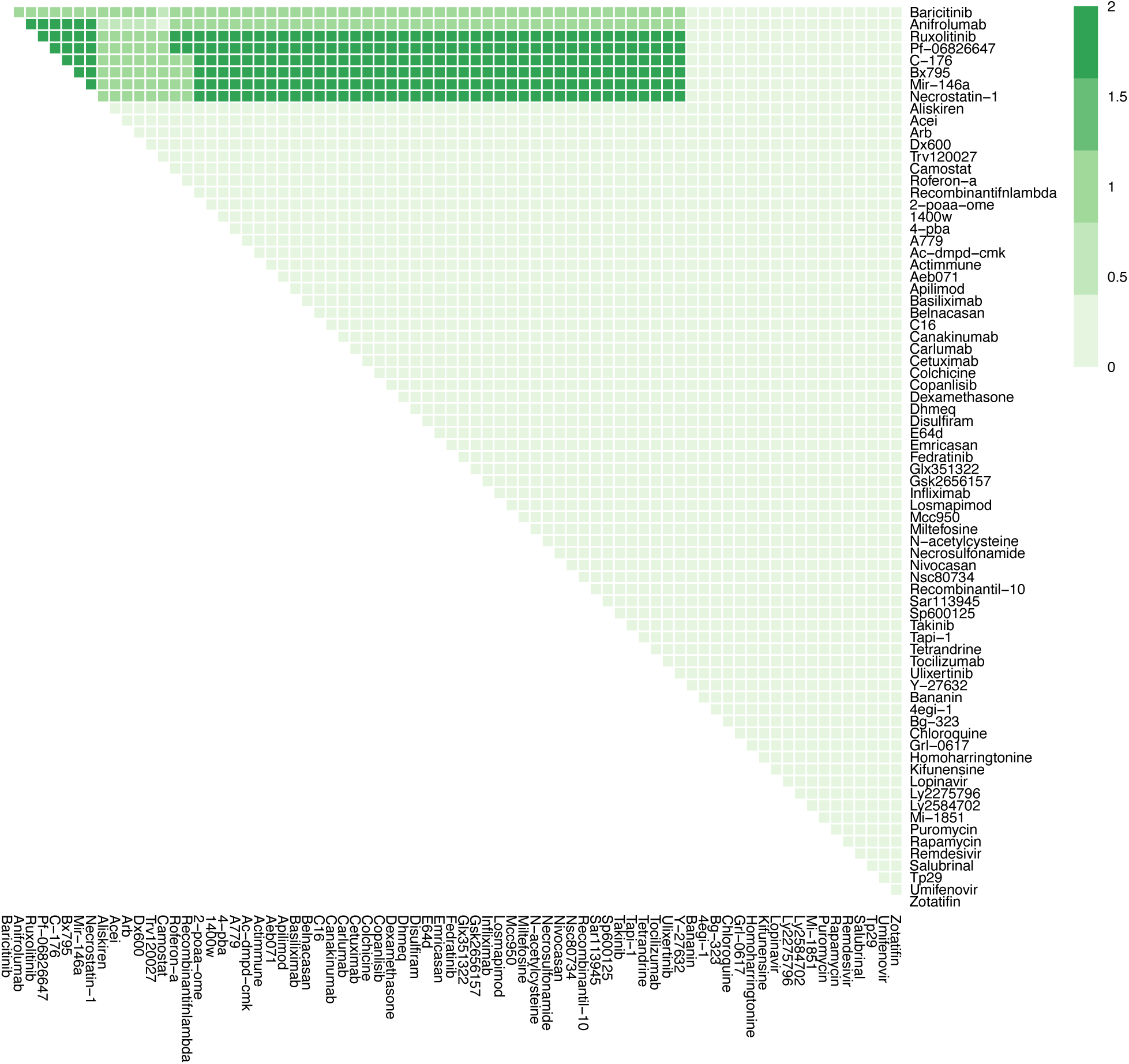
The effects of combination therapy on Syncytia Formation in late stage, severe COVID-19. The colour of the square corresponds to the level of the *SyncytiaFormation* node at steady state under treatment by the drugs indicated on the x- and y-axis. This plot shows all drugs, including those at an experimental stage of development.

**Figure S9N.**
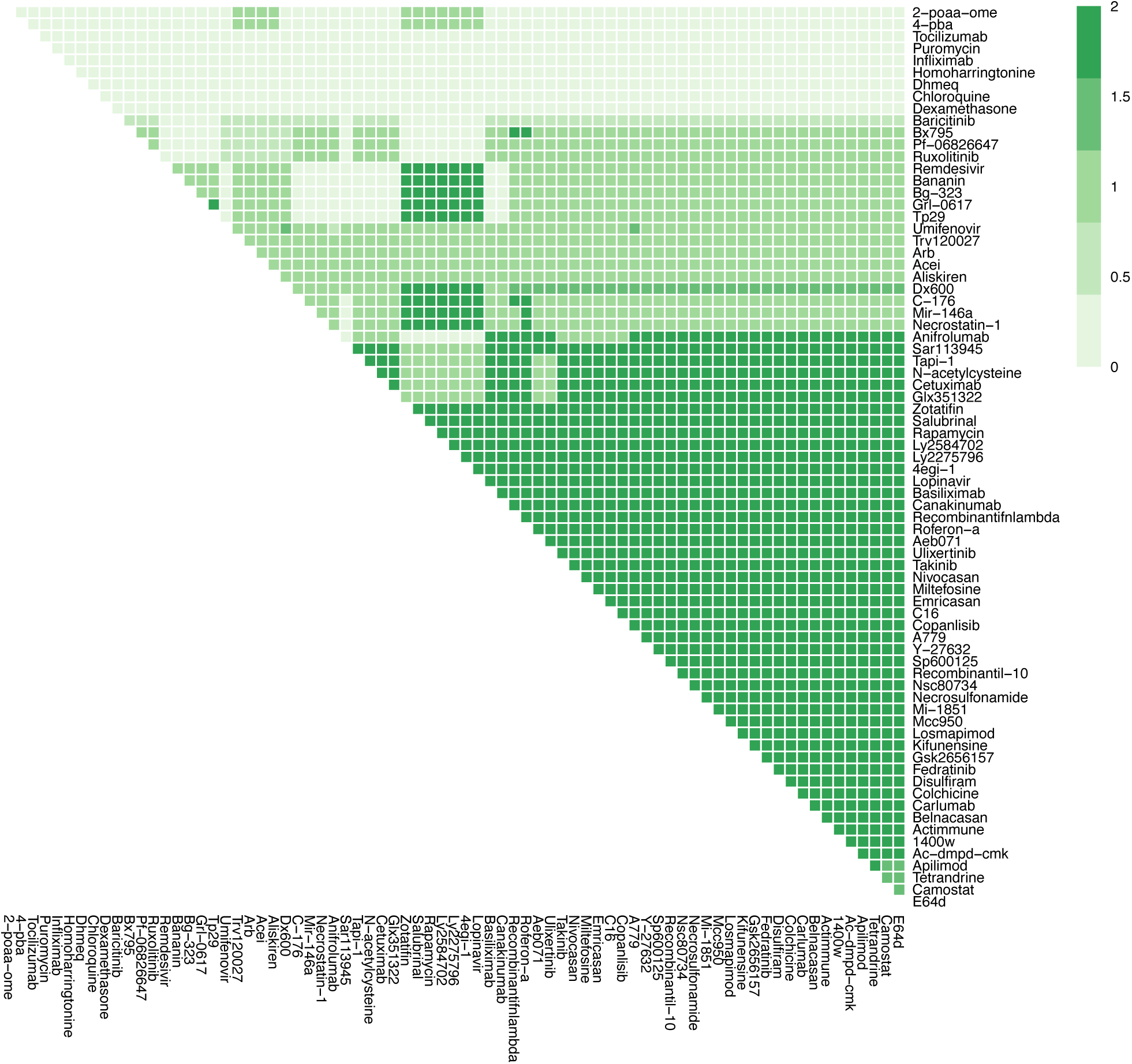
The effects of combination therapy on T-cell Infiltration in late stage, severe COVID-19. The colour of the square corresponds to the level of the *T-cellInfiltration* node at steady state under treatment by the drugs indicated on the x- and y-axis. This plot shows all drugs, including those at an experimental stage of development.

**Figure S9O.**
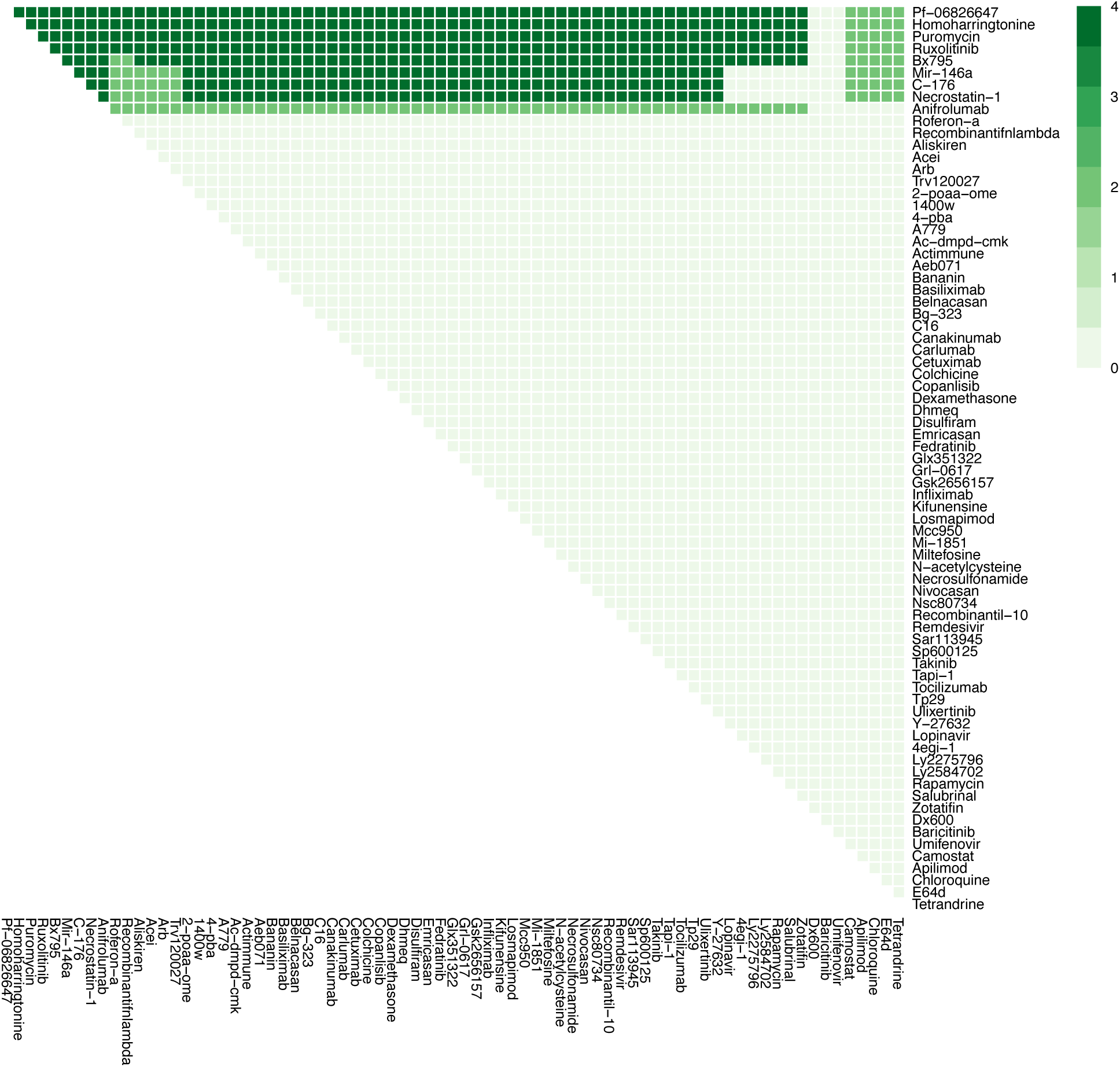
The effects of combination therapy on Viral Entry in late stage, severe COVID-19. The colour of the square corresponds to the level of the *ViralEntry* node at steady state under treatment by the drugs indicated on the x- and y-axis. This plot shows all drugs, including those at an experimental stage of development.

**Figure S9P.**
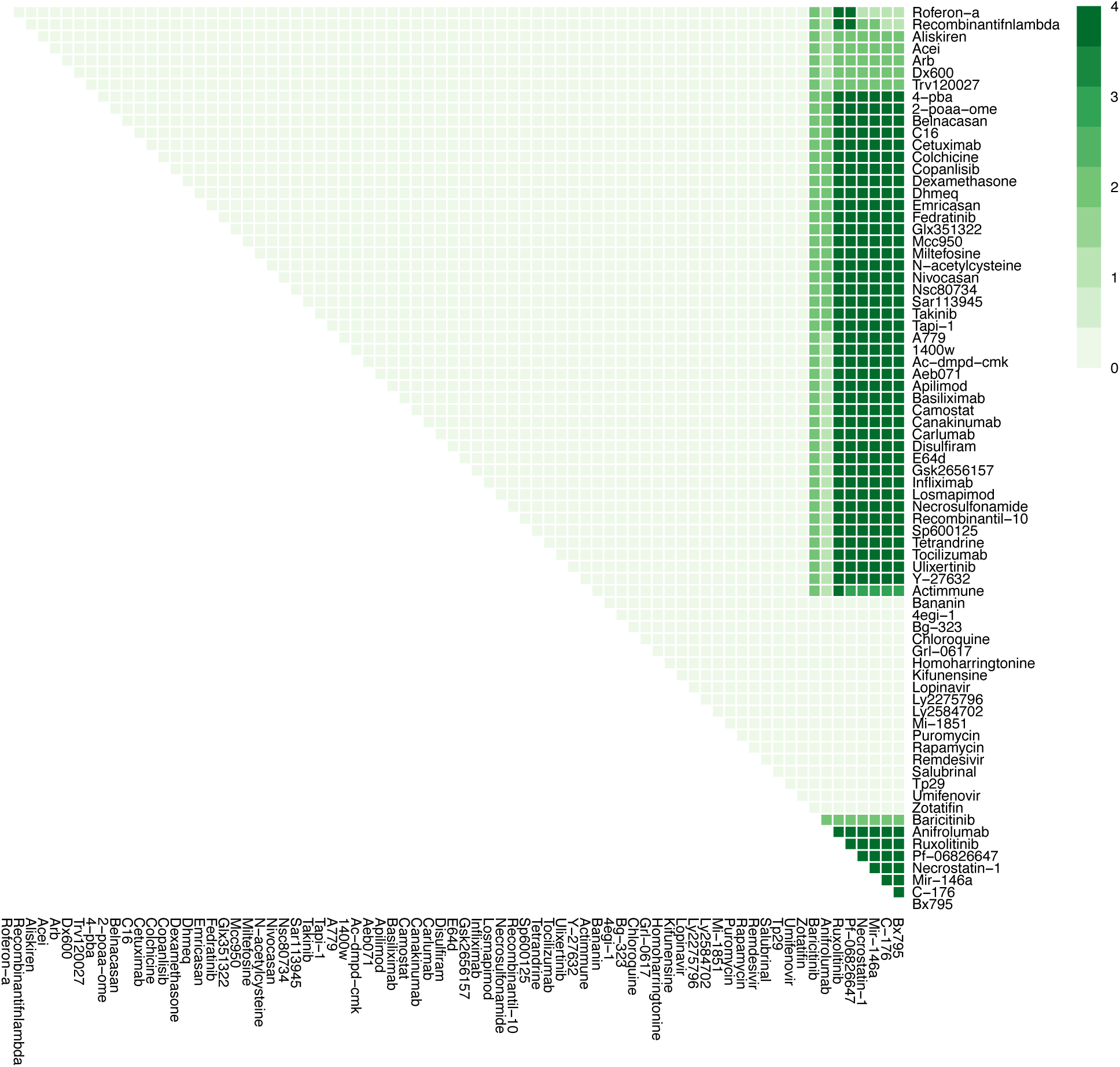
The effects of combination therapy on Viral Replication in late stage, severe COVID-19. The colour of the square corresponds to the level of the *ViralReplication* node at steady state under treatment by the drugs indicated on the x- and y-axis. This plot shows all drugs, including those at an experimental stage of development.

**Figure S10 A-D.**
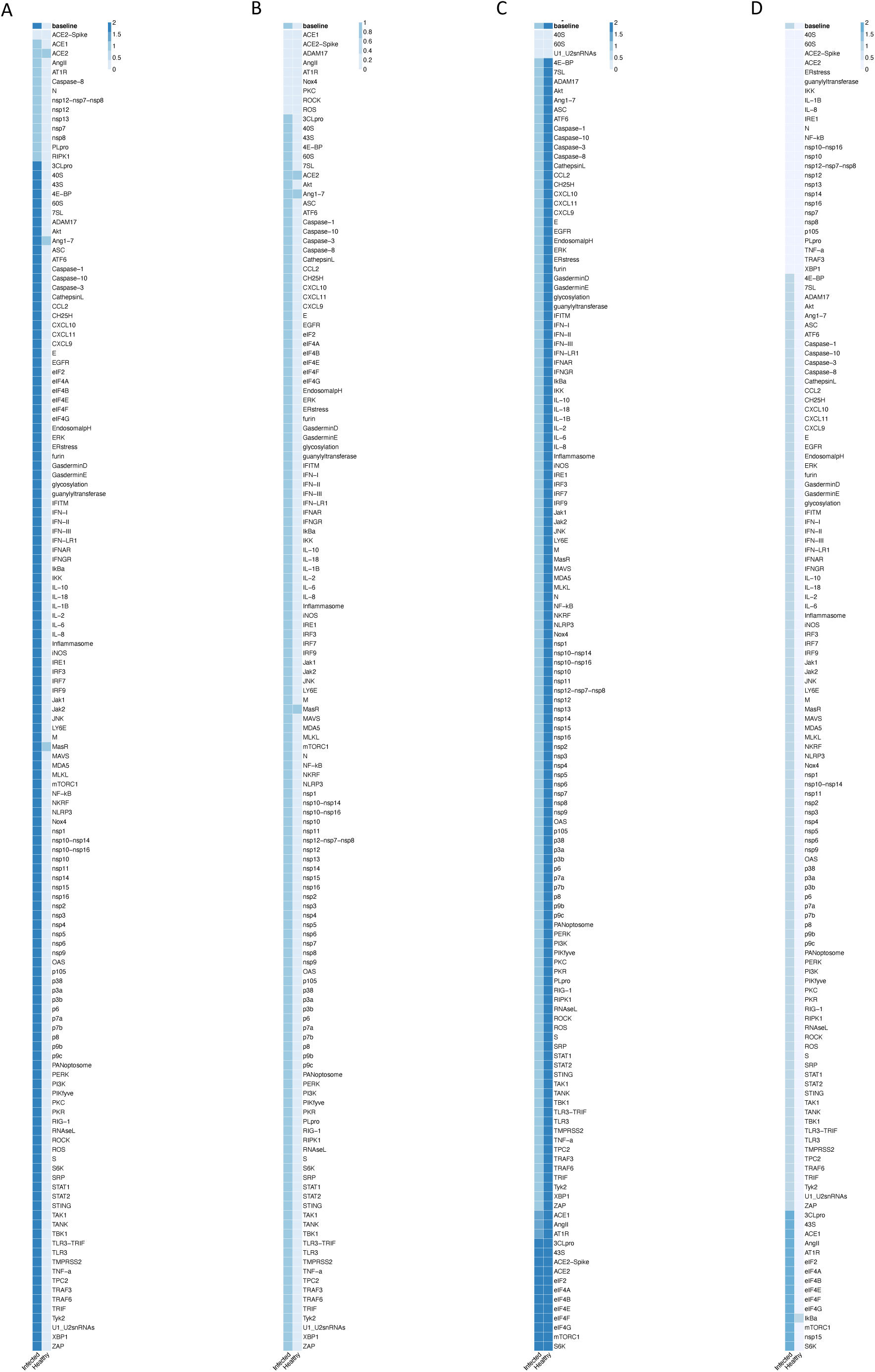
The effects of inhibition of individual nodes in early stage, severe COVID-19. Columns represent infected and healthy conditions, rows represent node inhibited. Showing the effect on: (A) Cell Death. (B) Fibrosis. (C) Host Protein Synthesis. (D) Inflammation.

**Figure S10 E-H.**
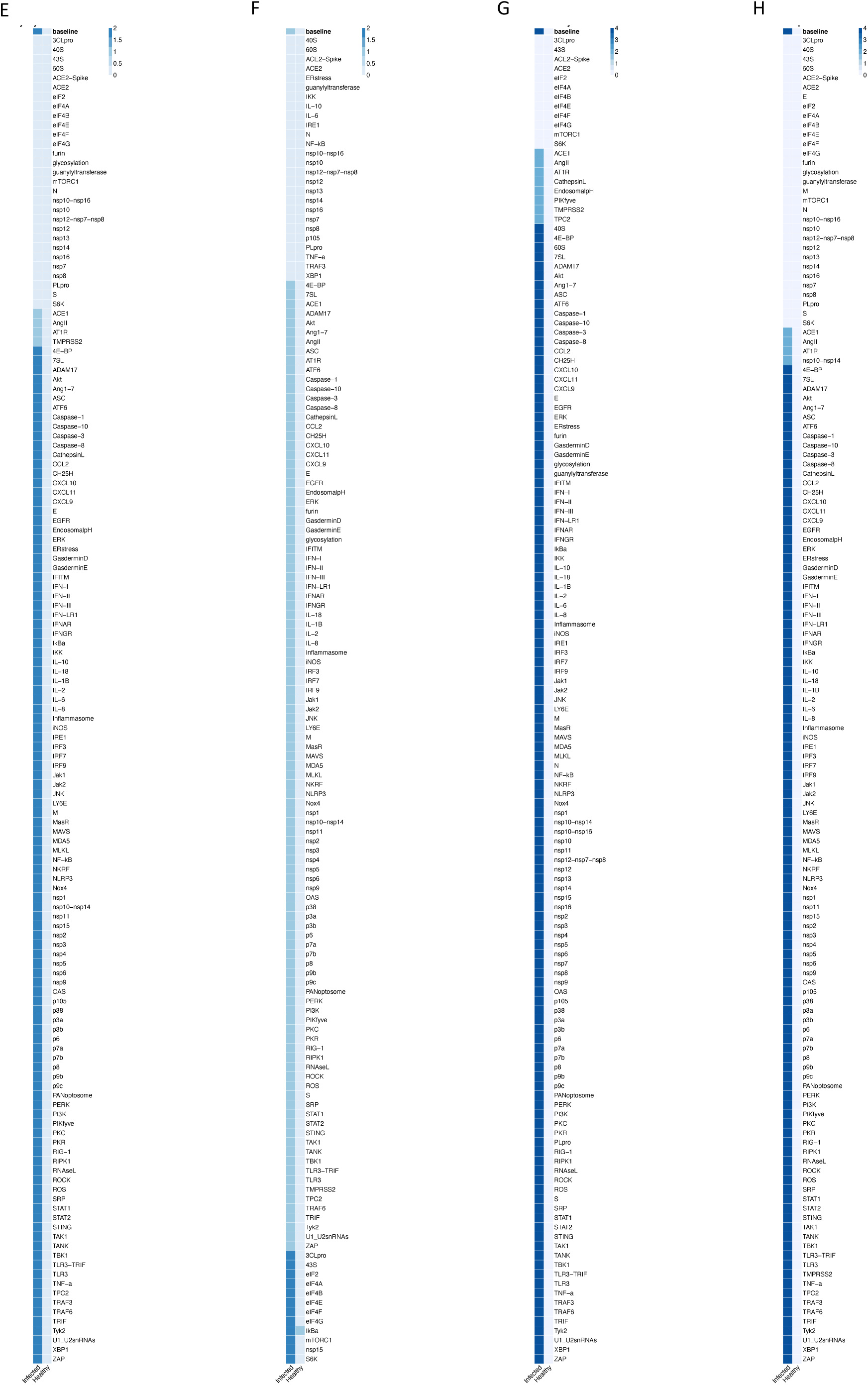
The effects of inhibition of individual nodes in early stage, severe COVID-19. Columns represent infected and healthy conditions, rows represent node inhibited. Showing the effect on: (E) Syncytia Formation. (F) T-cell Infiltration. (G) Viral Entry. (H) Viral Replication.

**Figure S10I.**
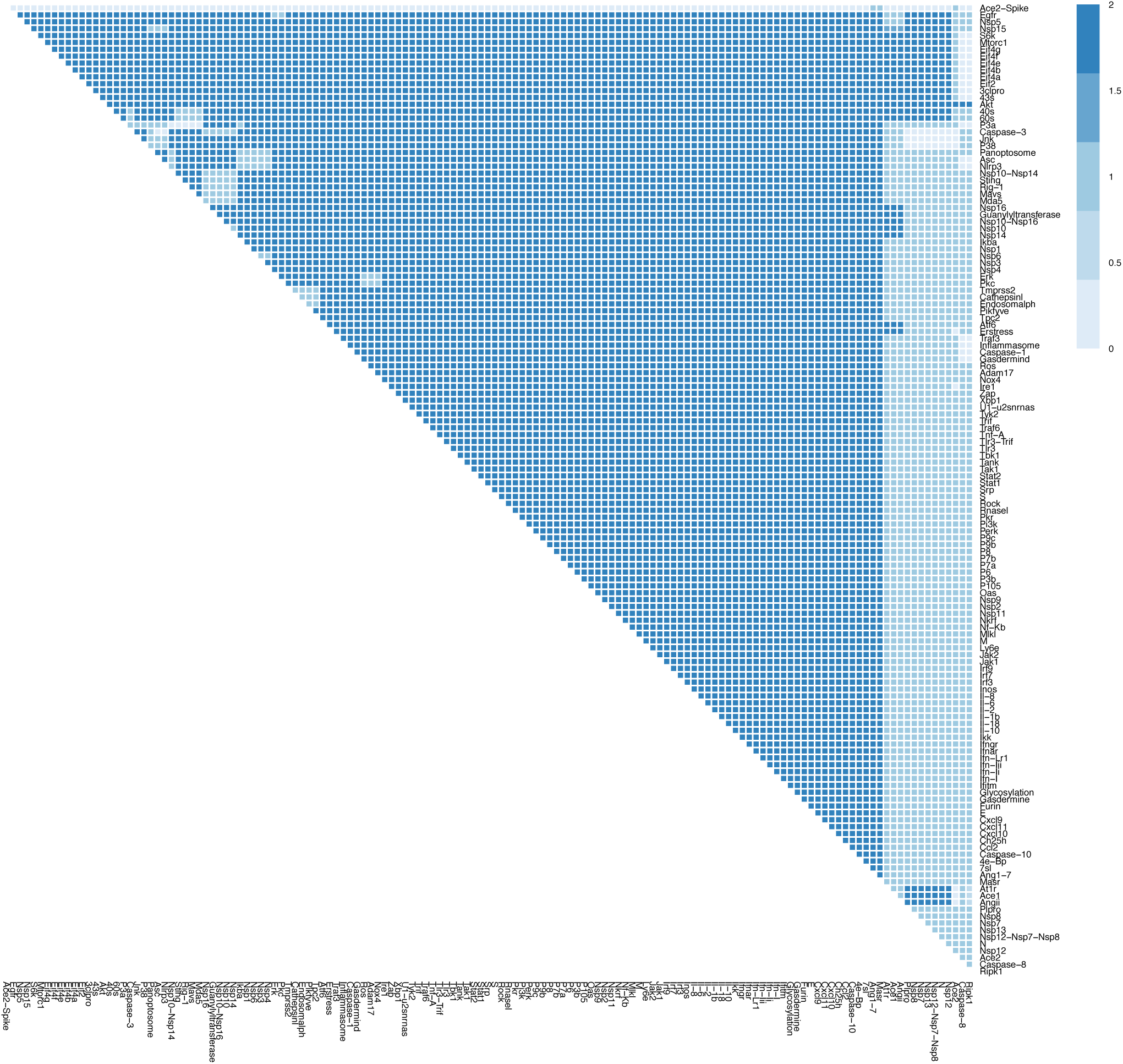
The effects of combination therapy on Cell Death in early stage, severe COVID-19. The colour of the square corresponds to the level of the *CellDeath* node at steady state upon inhbition of the nodes indicated on the x- and y-axis. This plot shows inhibition of all nodes representing druggable targets (proteins and complexes).

**Figure S10J.**
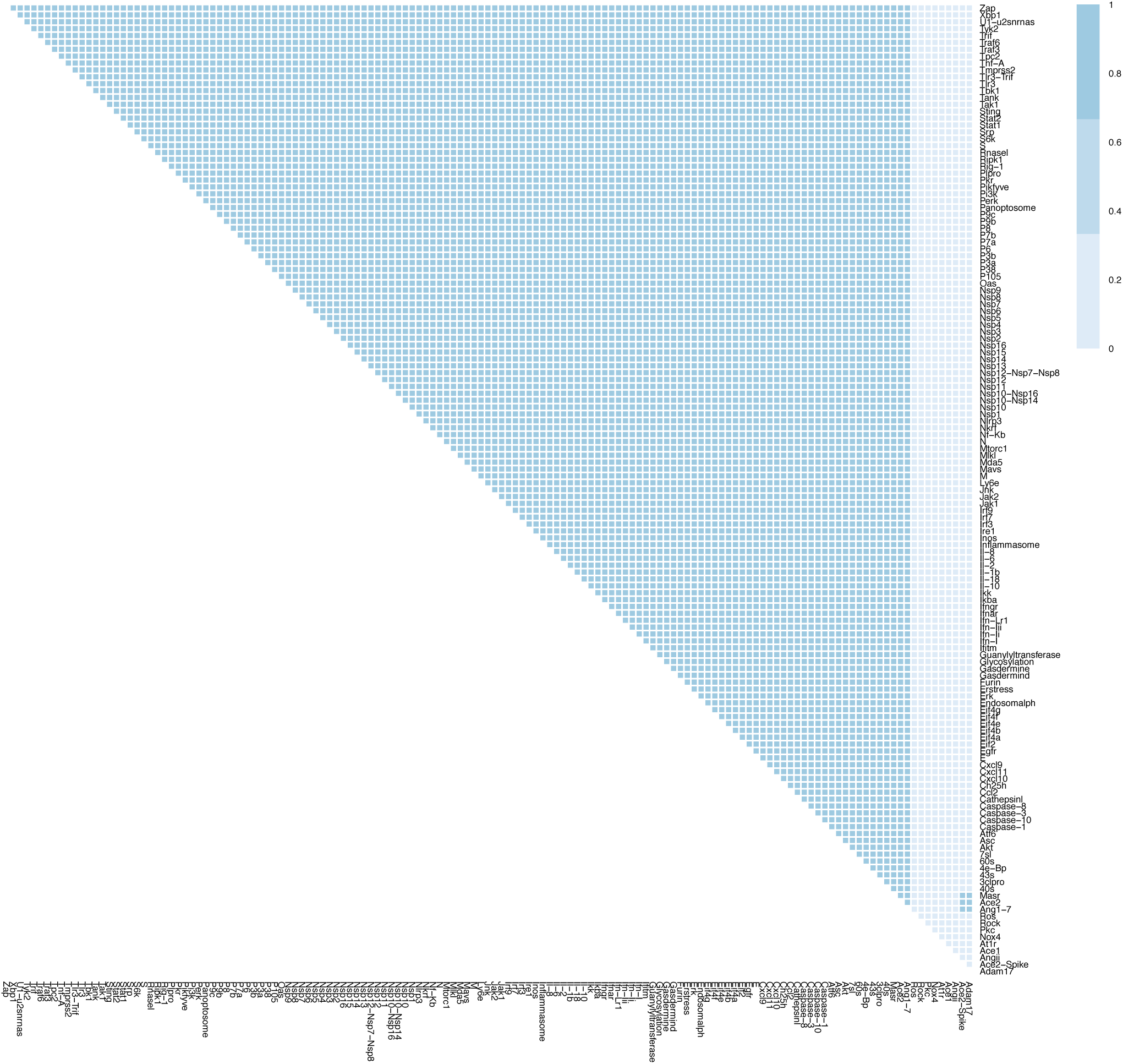
The effects of combination therapy on Fibrosis in early stage, severe COVID-19. The colour of the square corresponds to the level of the *Fibrosis* node at steady state upon inhbition of the nodes indicated on the x- and y-axis. This plot shows inhibition of all nodes representing drug-gable targets (proteins and complexes).

**Figure S10K.**
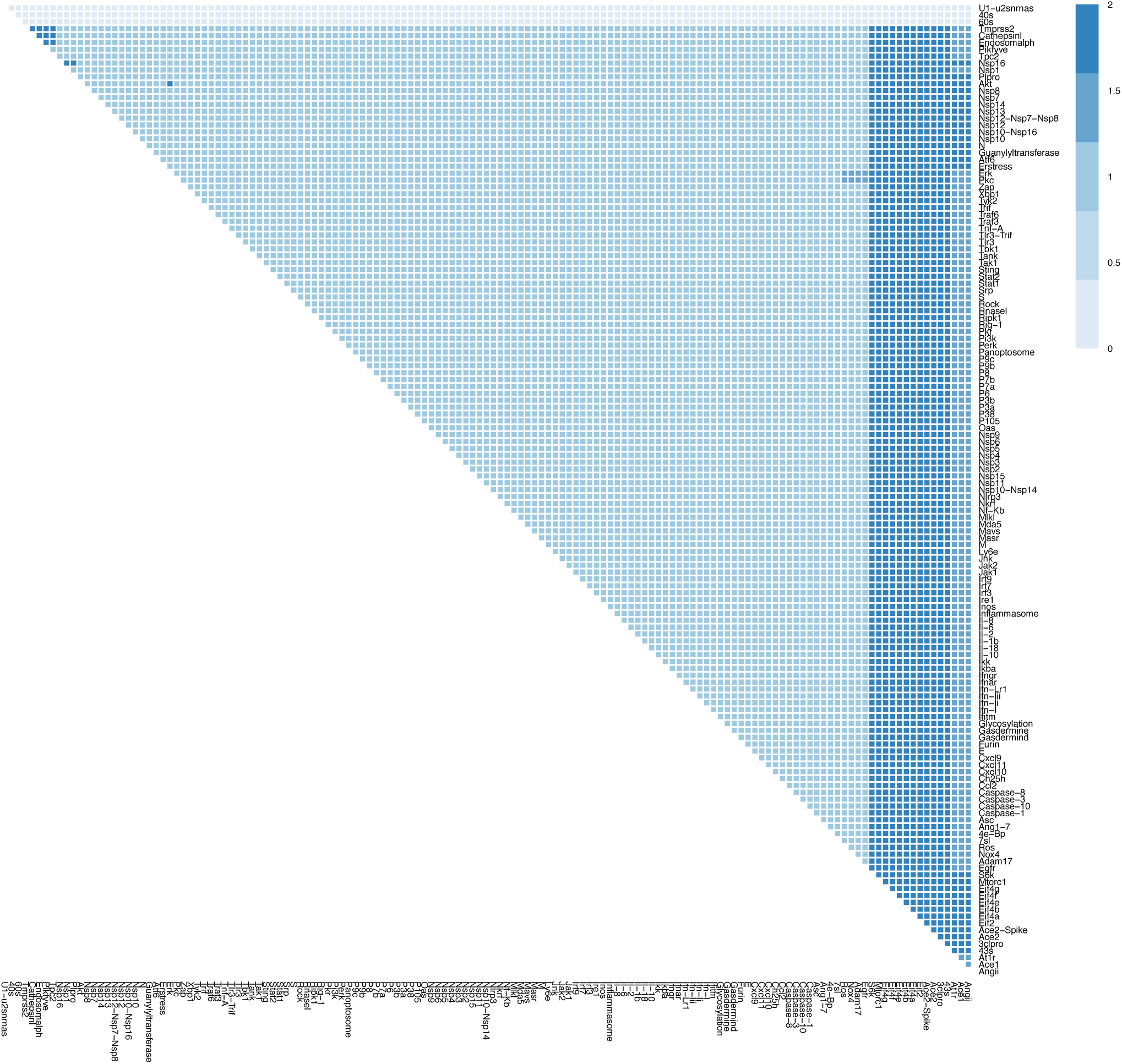
The effects of combination therapy on Host Protein Synthesis in early stage, severe COVID-19. The colour of the square corresponds to the level of the *HostProteinSynthesis* node at steady state upon inhbition of the nodes indicated on the x- and y-axis. This plot shows inhibition of all nodes representing druggable targets (proteins and complexes).

**Figure S10L.**
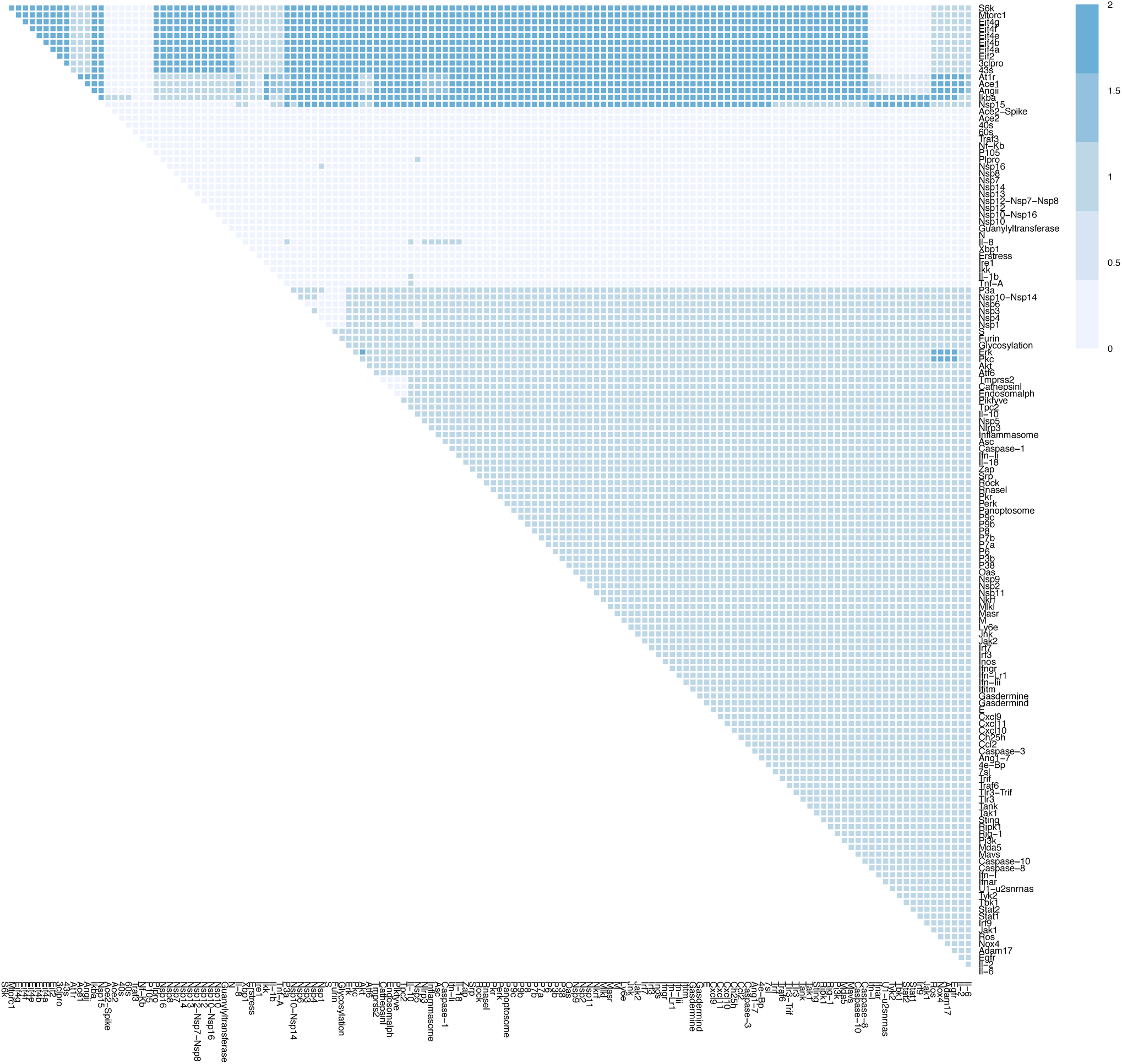
The effects of combination therapy on Inflammation in early stage, severe COVID-19. The colour of the square corresponds to the level of the *Inflammation* node at steady state upon inhbition of the nodes indicated on the x- and y-axis. This plot shows inhibition of all nodes representing druggable targets (proteins and complexes).

**Figure S10M.**
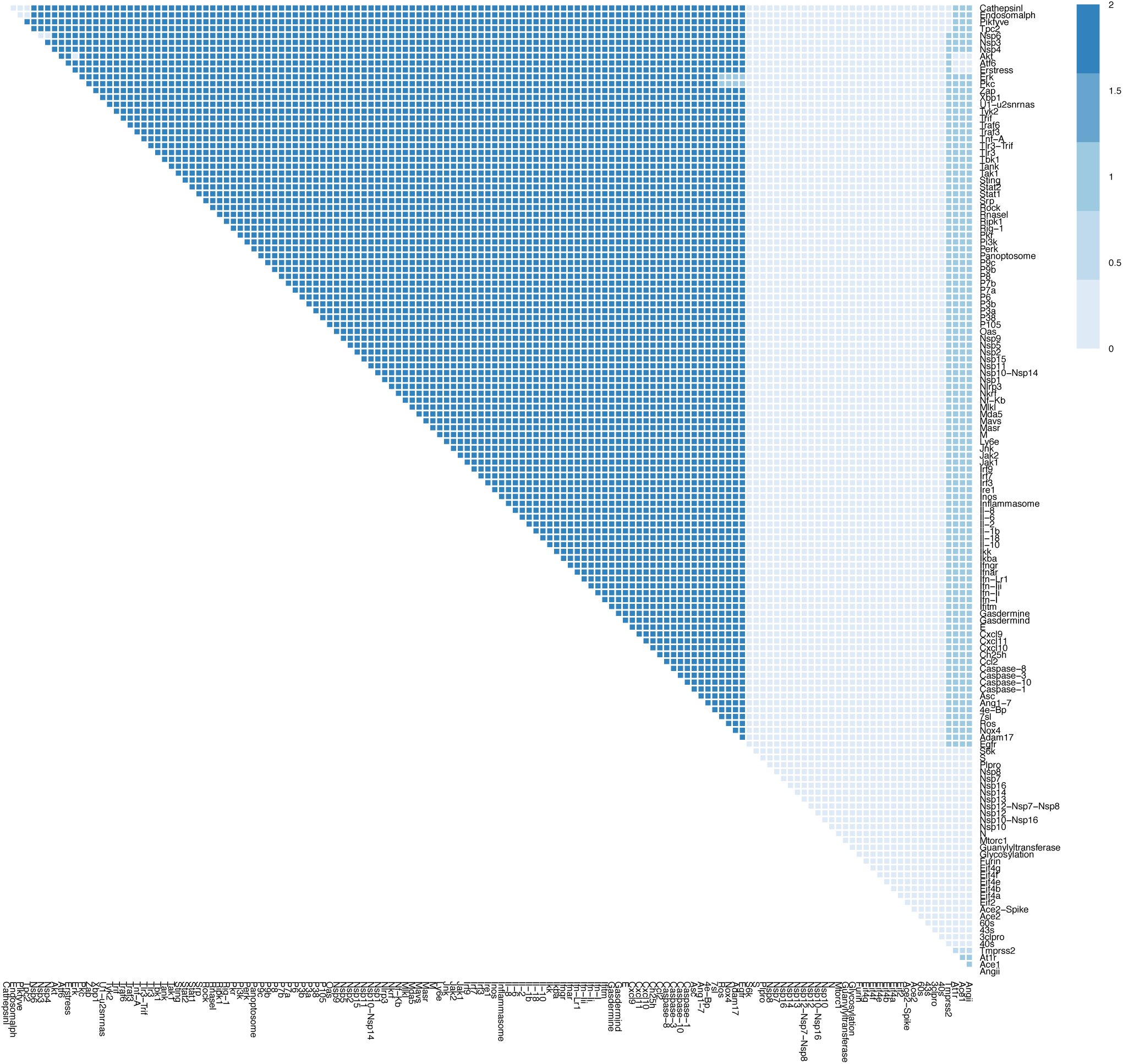
The effects of combination therapy on Syncytia Formation in early stage, severe COVID-19. The colour of the square corresponds to the level of the *SynctytiaFormation* node at steady state upon inhbition of the nodes indicated on the x- and y-axis. This plot shows inhibition of all nodes representing druggable targets (proteins and complexes).

**Figure S10N.**
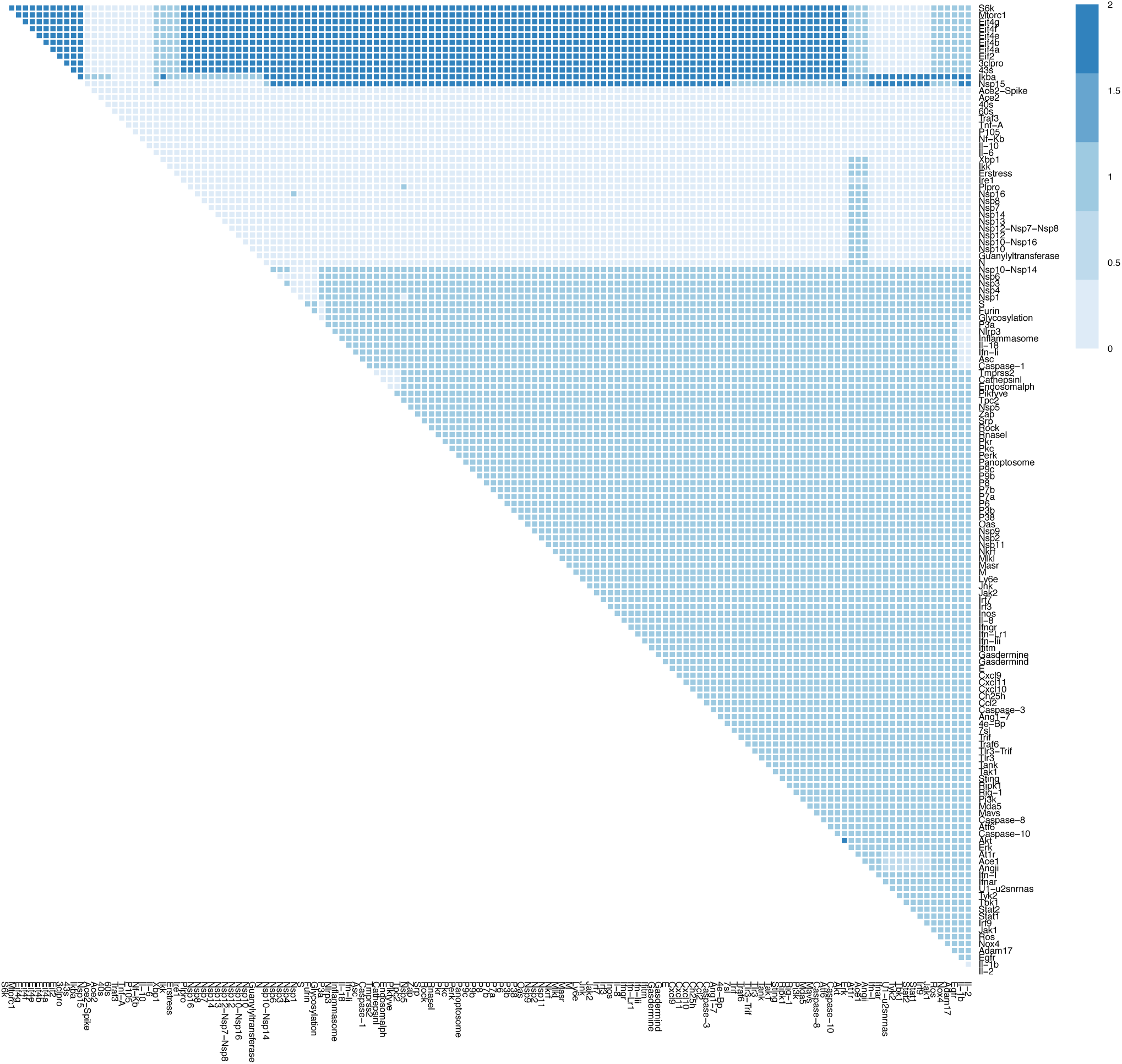
The effects of combination therapy on T-cell Infiltration in early stage, severe COVID-19. The colour of the square corresponds to the level of the *T-cellInfiltration* node at steady state upon inhbition of the nodes indicated on the x- and y-axis. This plot shows inhibition of all nodes representing druggable targets (proteins and complexes).

**Figure S10O.**
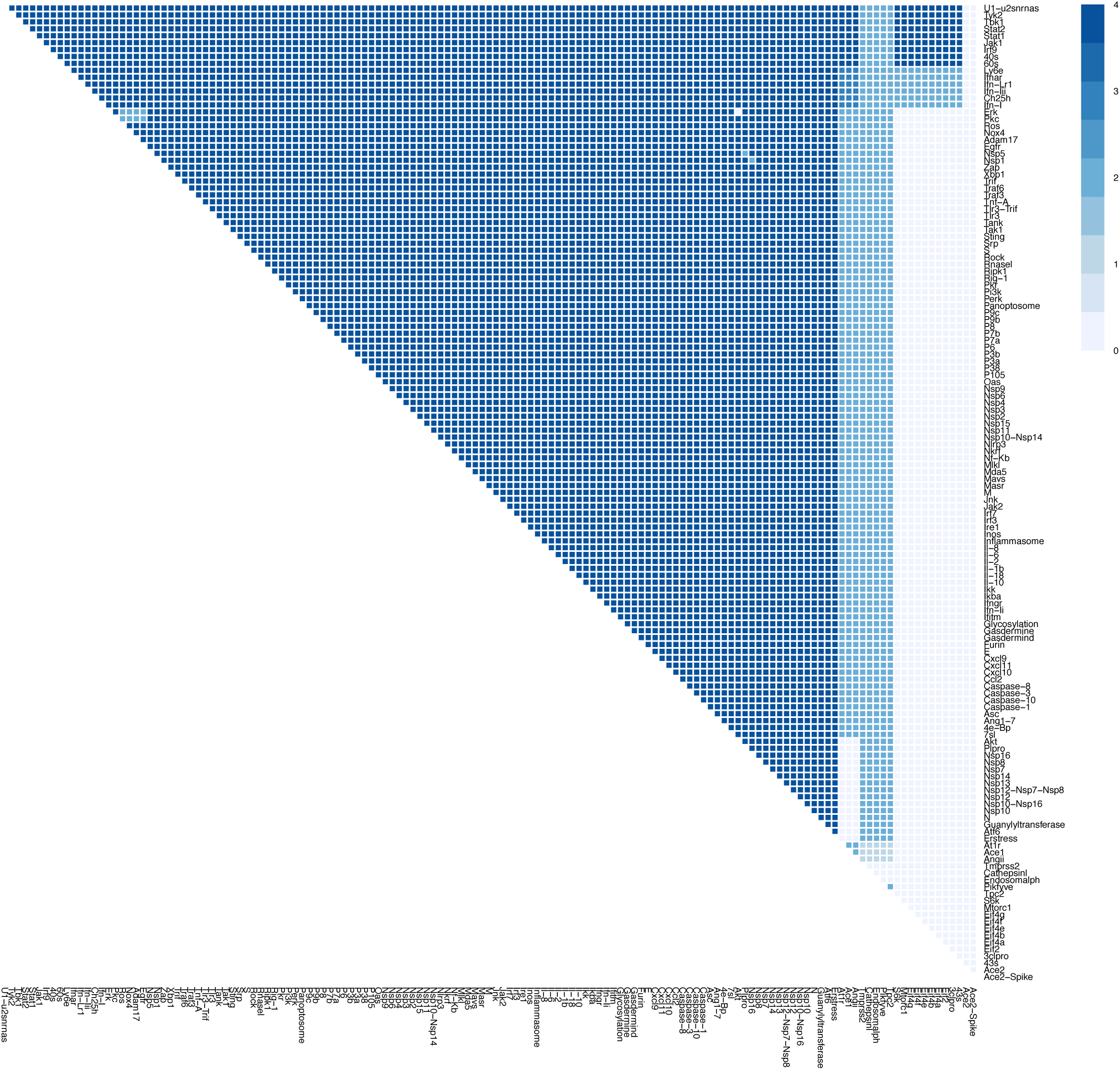
The effects of combination therapy on Viral Entry in early stage, severe COVID-19. The colour of the square corresponds to the level of the *ViralEntry* node at steady state upon inhbition of the nodes indicated on the x- and y-axis. This plot shows inhibition of all nodes representing druggable targets (proteins and complexes).

**Figure S10P.**
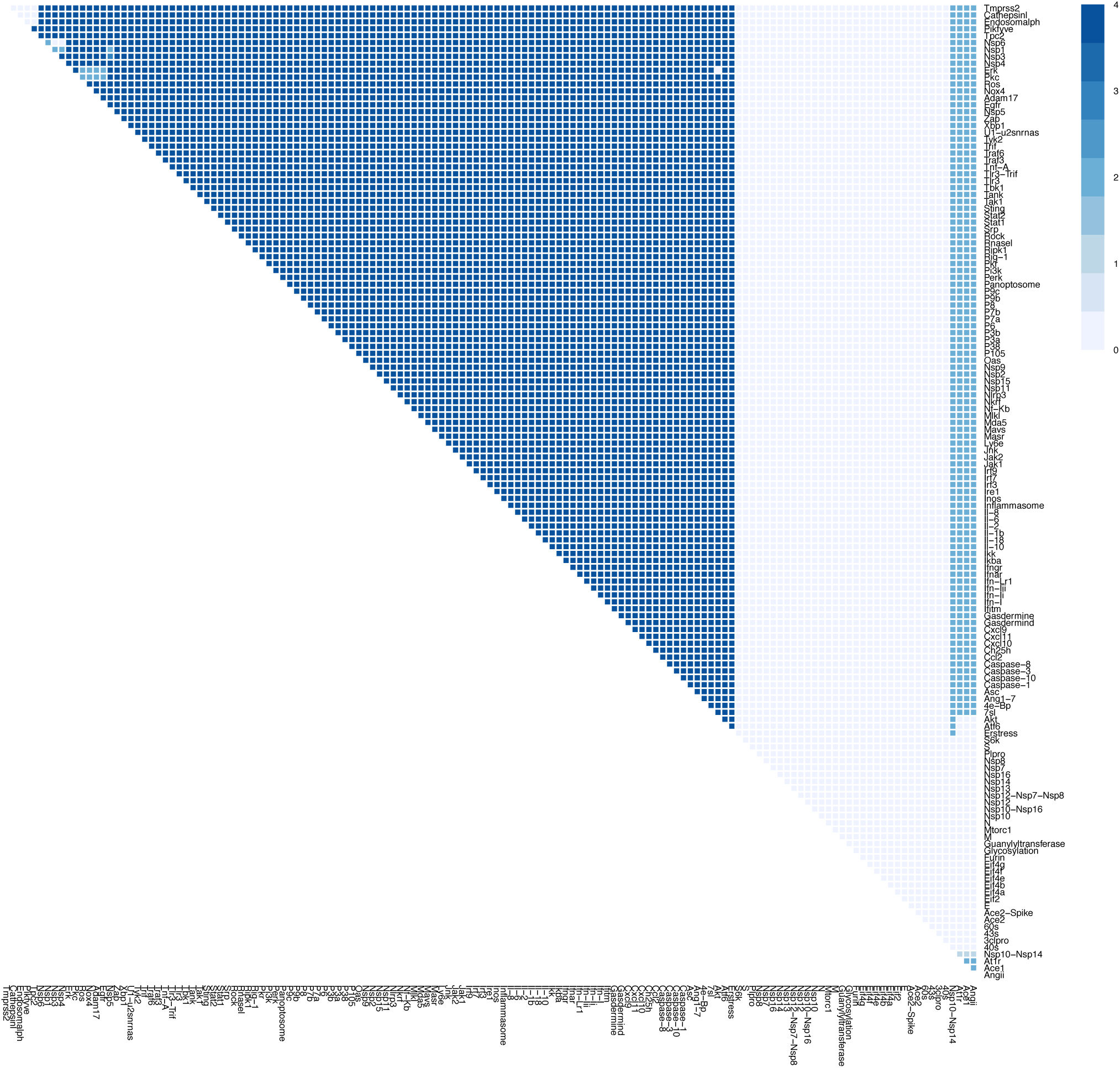
The effects of combination therapy on Viral Replication in early stage, severe COVID-19. The colour of the square corresponds to the level of the *ViralReplication* node at steady state upon inhbition of the nodes indicated on the x- and y-axis. This plot shows inhibition of all nodes representing druggable targets (proteins and complexes).

**Figure S11 A-D.**
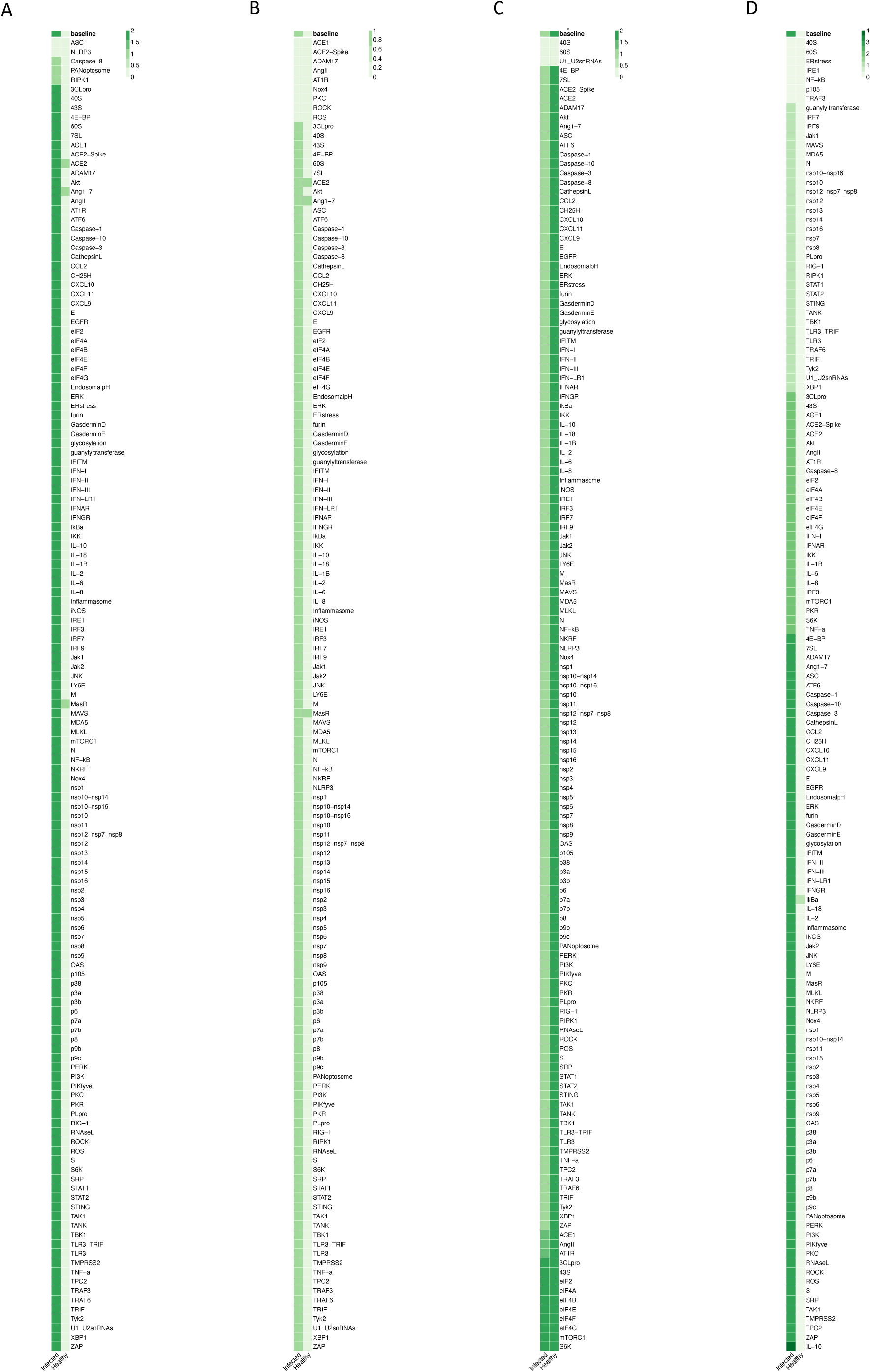
The effects of inhibition of individual nodes in late stage, severe COVID-19. Columns represent infected and healthy conditions, rows represent node inhibited. Showing the effect on: (A) Cell Death. (B) Fibrosis. (C) Host Protein Synthesis. (D) Inflammation.

**Figure S11 E-H.**
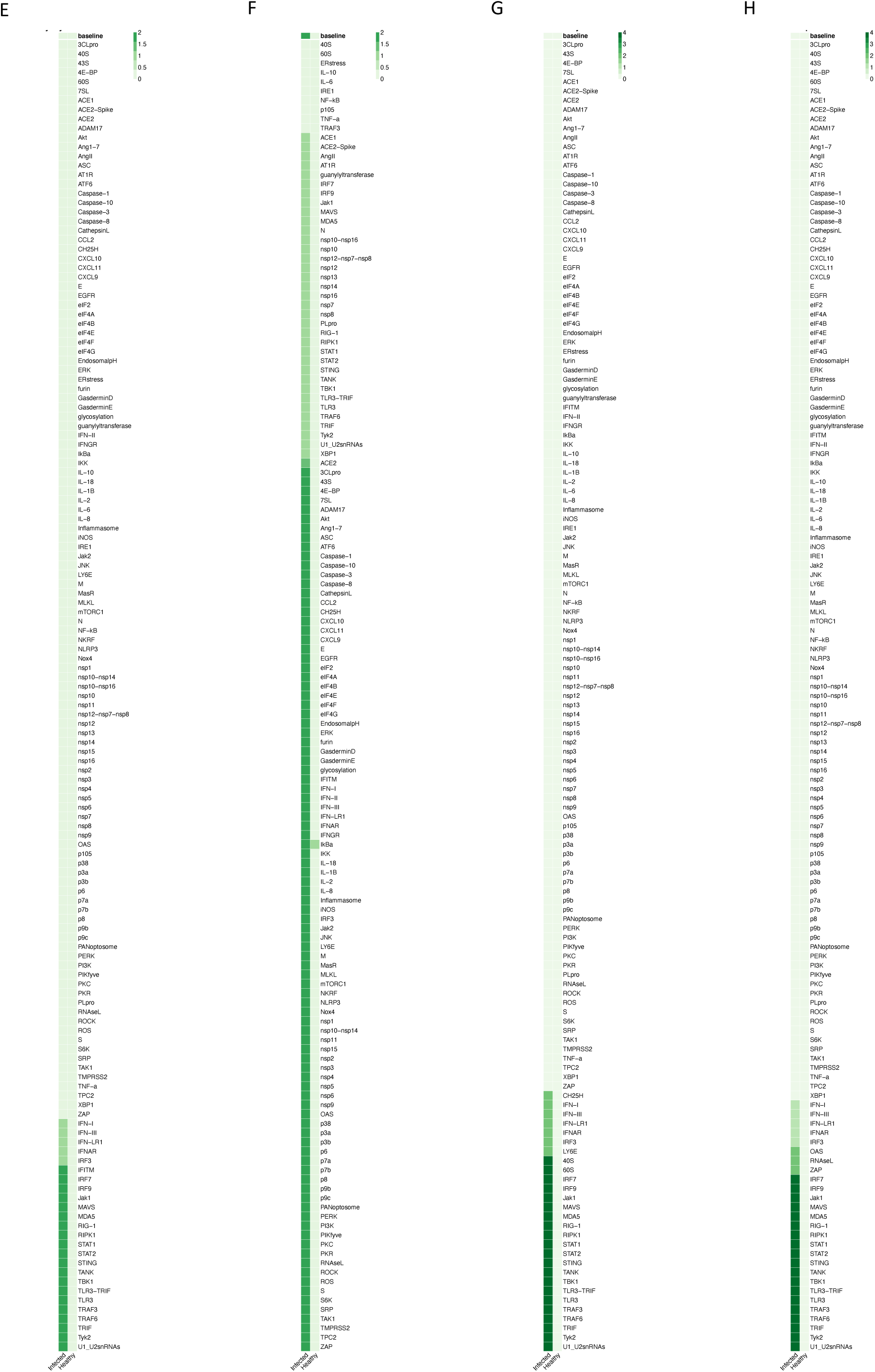
The effects of inhibition of individual nodes in late stage, severe COVID-19. Columns represent infected and healthy conditions, rows represent node inhibited. Showing the effect on: (E) Syncytia Formation. (F) T-cell Infiltration. (G) Viral Entry. (H) Viral Replication.

**Figure S11I.**
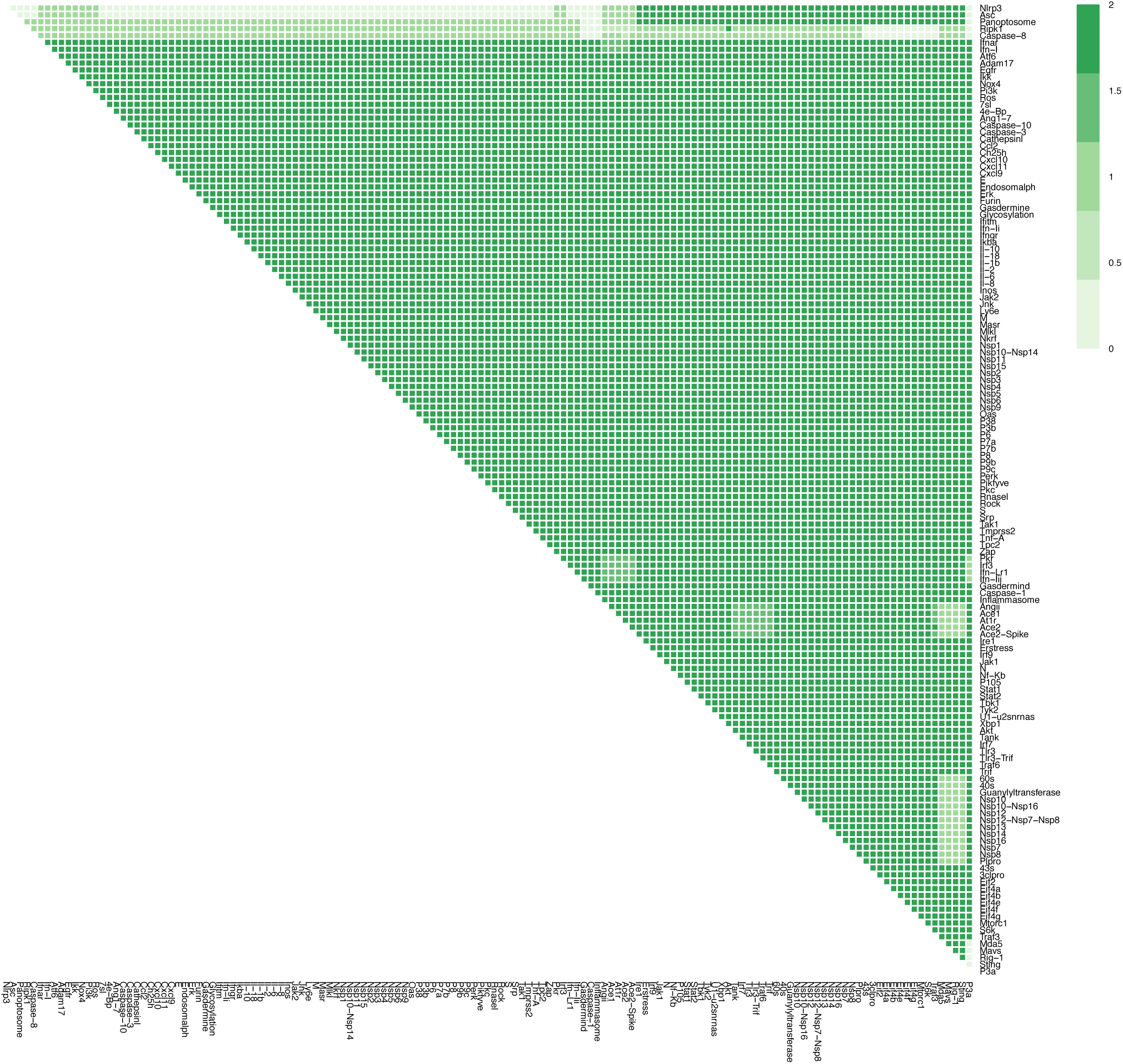
The effects of combination therapy on Cell Death in late stage, severe COVID-19. The colour of the square corresponds to the level of the C*ellDeath* node at steady state upon inhbition of the nodes indicated on the x- and y-axis. This plot shows inhibition of all nodes representing druggable targets (proteins and complexes).

**Figure S11J.**
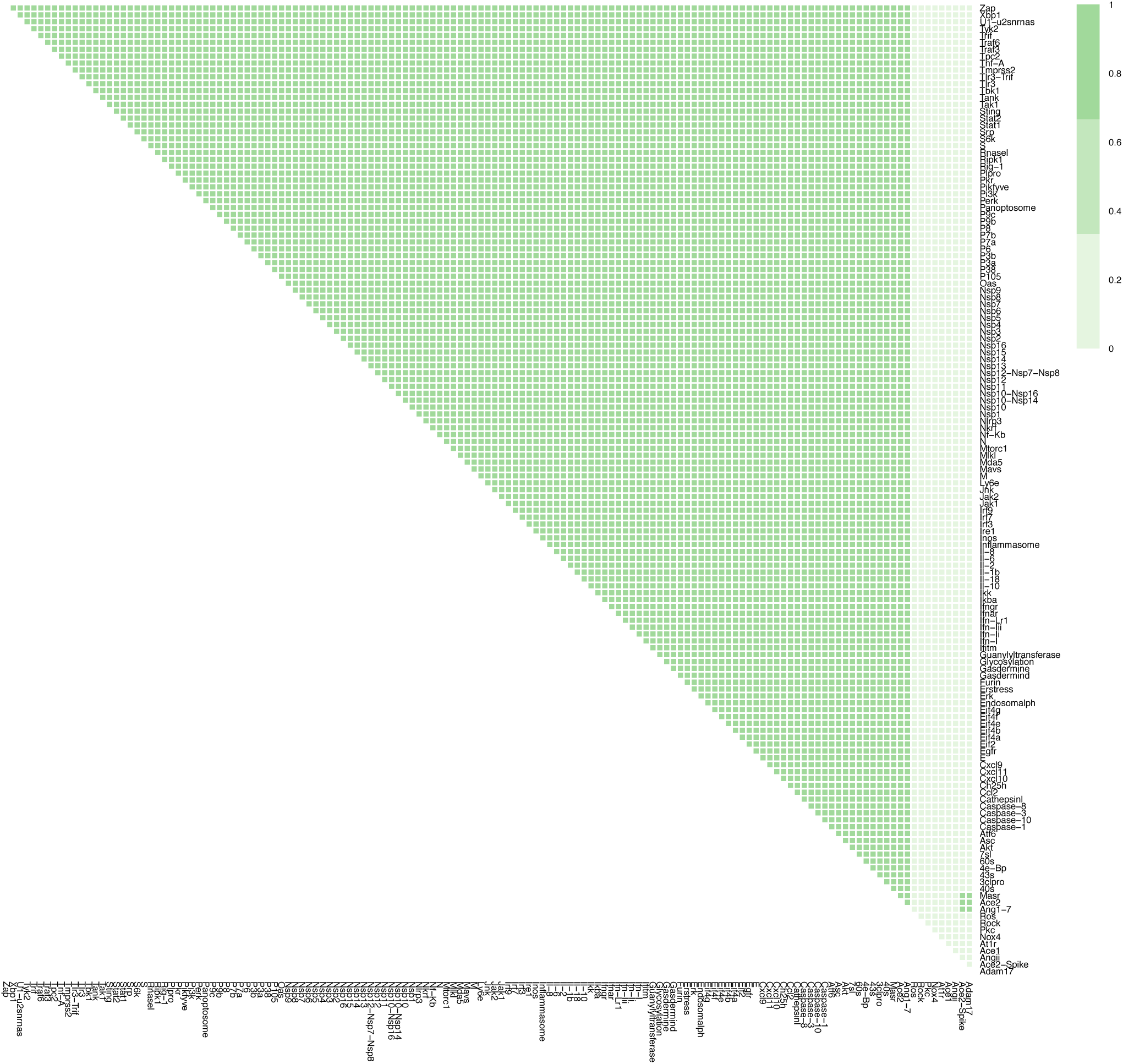
The effects of combination therapy on Fibrosis in late stage, severe COVID-19. The colour of the square corresponds to the level of the *Fibrosis* node at steady state upon inhbition of the nodes indicated on the x- and y-axis. This plot shows inhibition of all nodes representing drug-gable targets (proteins and complexes).

**Figure S11K.**
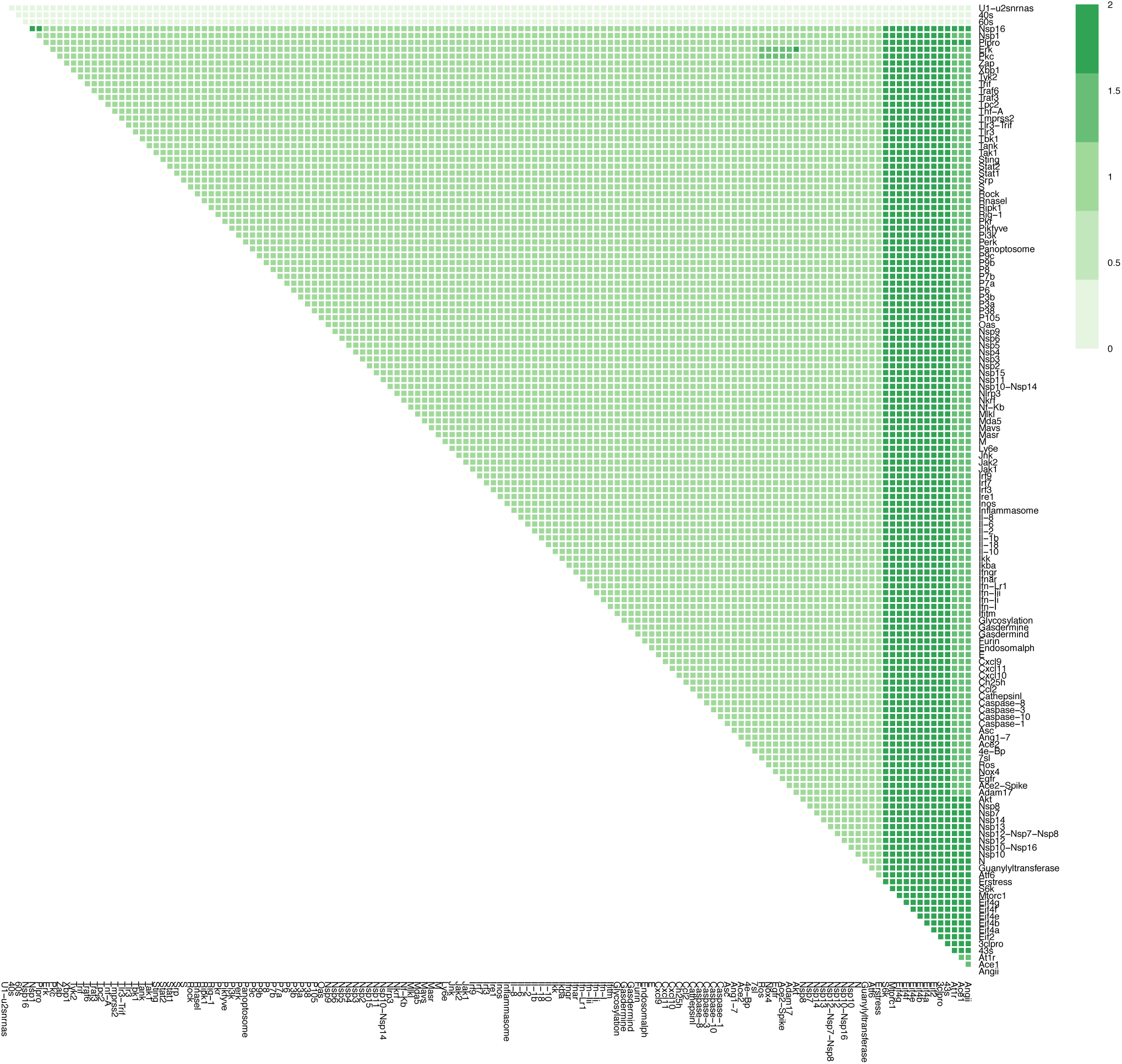
The effects of combination therapy on Host Protein Synthesis in late stage, severe COVID-19. The colour of the square corresponds to the level of the *HostProteinSynthesis* node at steady state upon inhbition of the nodes indicated on the x- and y-axis. This plot shows inhibition of all nodes representing druggable targets (proteins and complexes).

**Figure S11L.**
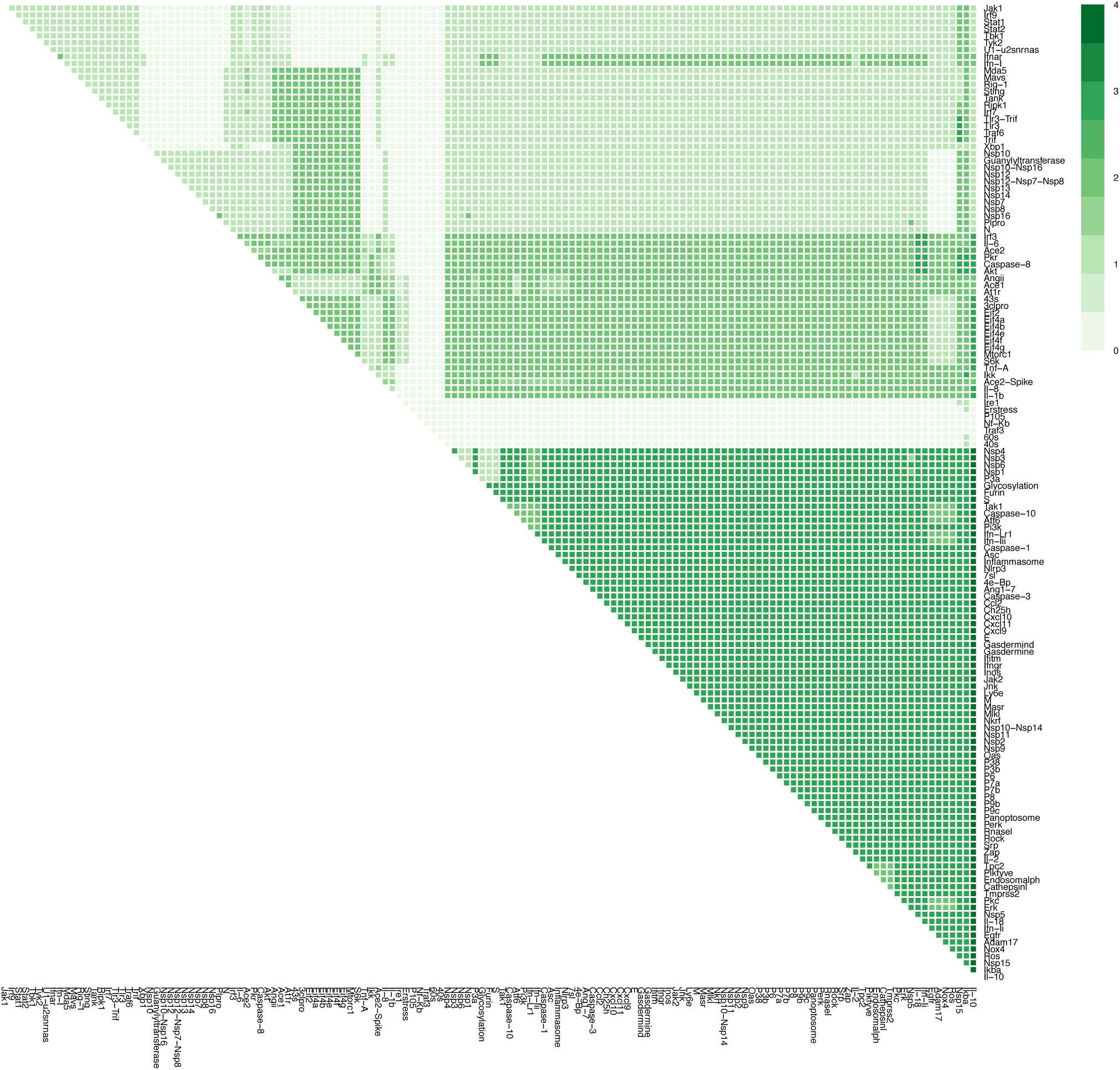
The effects of combination therapy on Inflammation in late stage, severe COVID-19. The colour of the square corresponds to the level of the *Inflammation* node at steady state upon inhbition of the nodes indicated on the x- and y-axis. This plot shows inhibition of all nodes representing druggable targets (proteins and complexes).

**Figure S11M.**
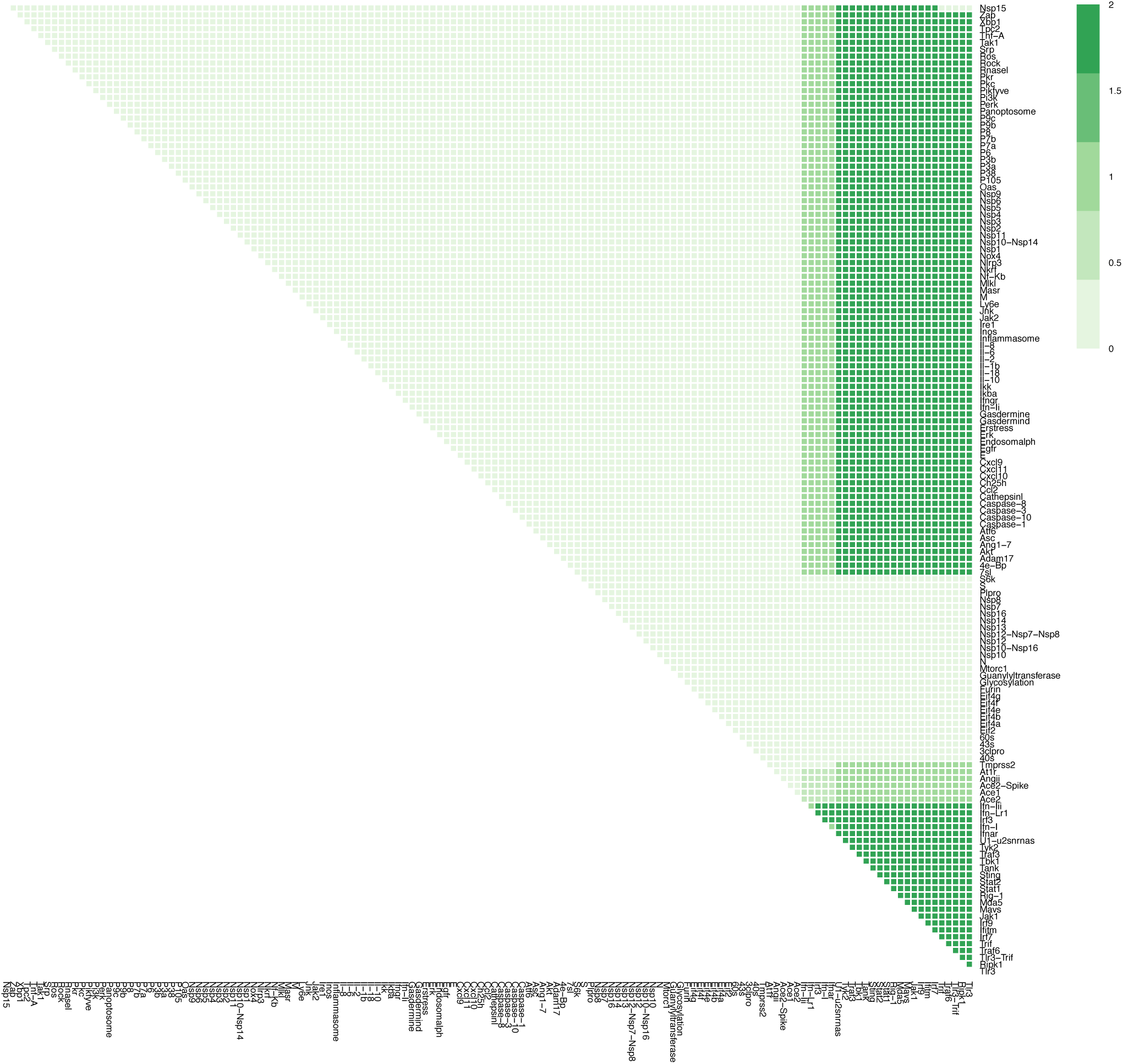
The effects of combination therapy on Syncytia Formation in late stage, severe COVID-19. The colour of the square corresponds to the level of the *SyncytiaFormation* node at steady state upon inhbition of the nodes indicated on the x- and y-axis. This plot shows inhibition of all nodes representing druggable targets (proteins and complexes).

**Figure S11N.**
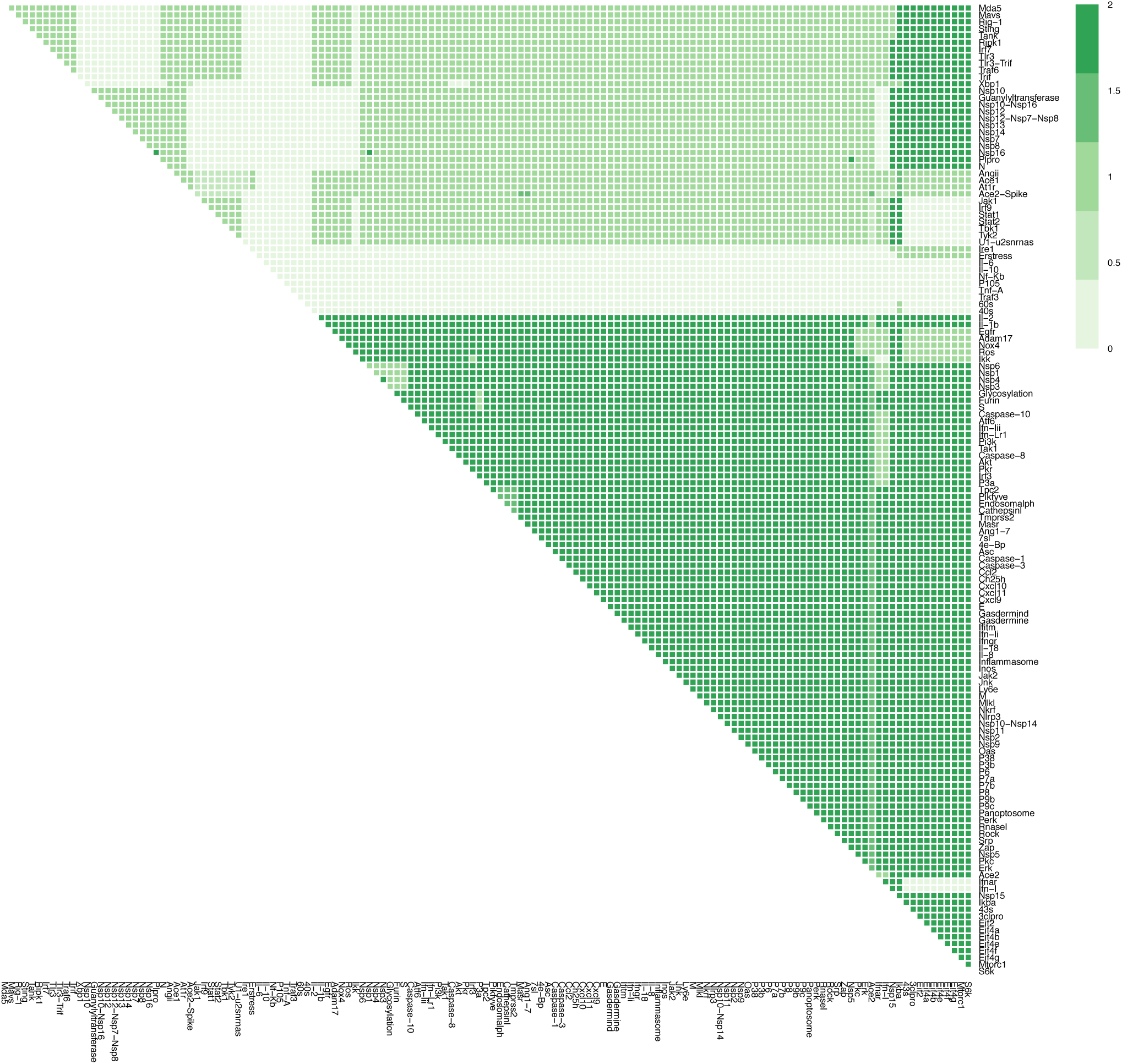
The effects of combination therapy on T-cell Infiltration in late stage, severe COVID-19. The colour of the square corresponds to the level of the *T-cellInfiltration* node at steady state upon inhbition of the nodes indicated on the x- and y-axis. This plot shows inhibition of all nodes representing druggable targets (proteins and complexes).

**Figure S11O.**
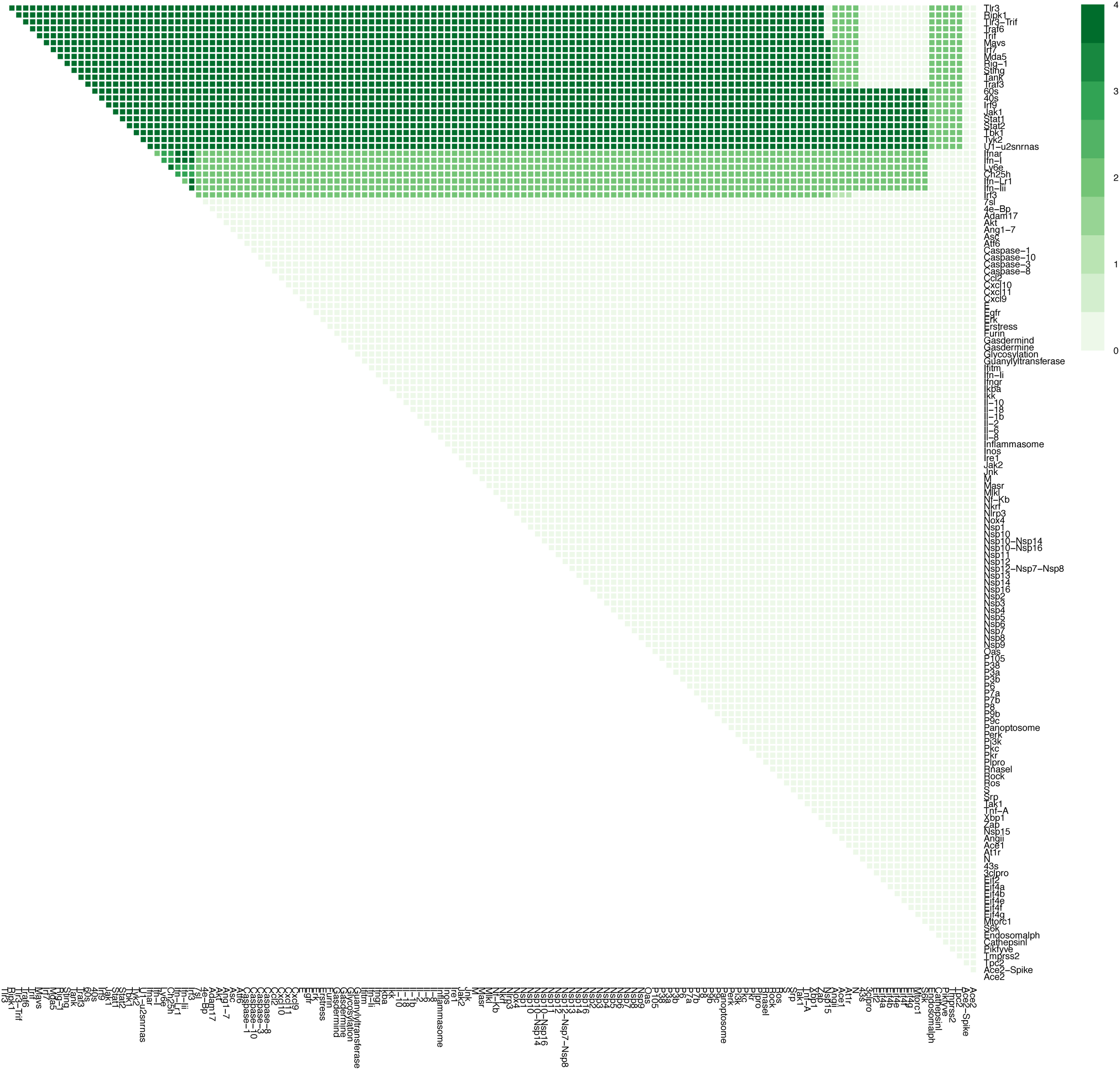
The effects of combination therapy on Viral Entry in late stage, severe COVID-19. The colour of the square corresponds to the level of the *ViralEntry* node at steady state upon inhbition of the nodes indicated on the x- and y-axis. This plot shows inhibition of all nodes representing druggable targets (proteins and complexes).

**Figure S11P.**
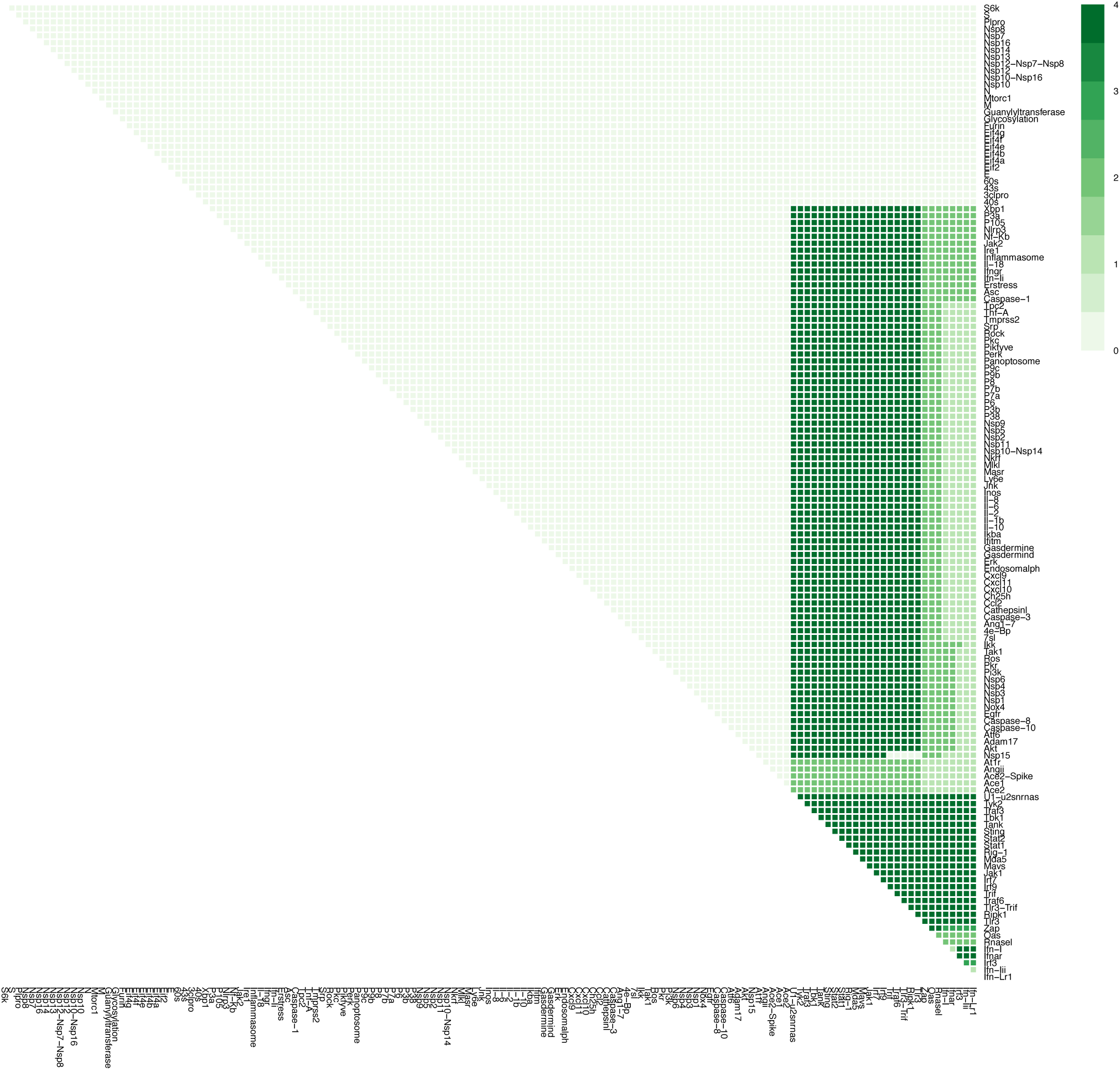
The effects of combination therapy on Viral Replication in late stage, severe COVID-19. The colour of the square corresponds to the level of the *ViralReplication* node at steady state upon inhbition of the nodes indicated on the x- and y-axis. This plot shows inhibition of all nodes representing druggable targets (proteins and complexes).

**Figure S12.**
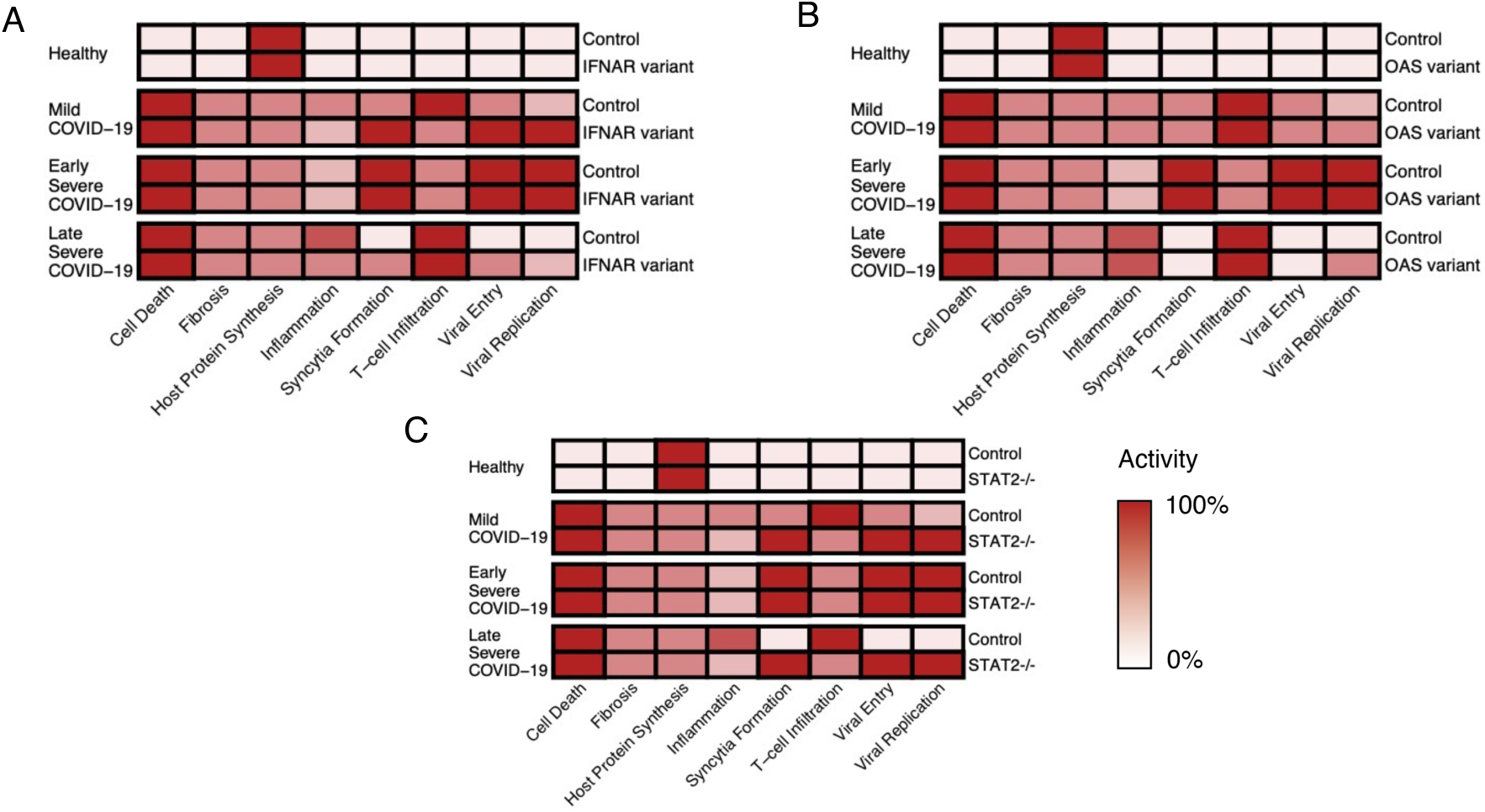
Genetic risk factors associated with severe COVID19. (A-B) The behaviour of loss-of-function mutations in IFNAR and OAS under the four conditions simulated by the model. (C) The behaviour of loss-of-function mutations in STAT2 under the four conditions simulated by the model. All nodes normalized to be between 0 and 100%.

**Table S1.**
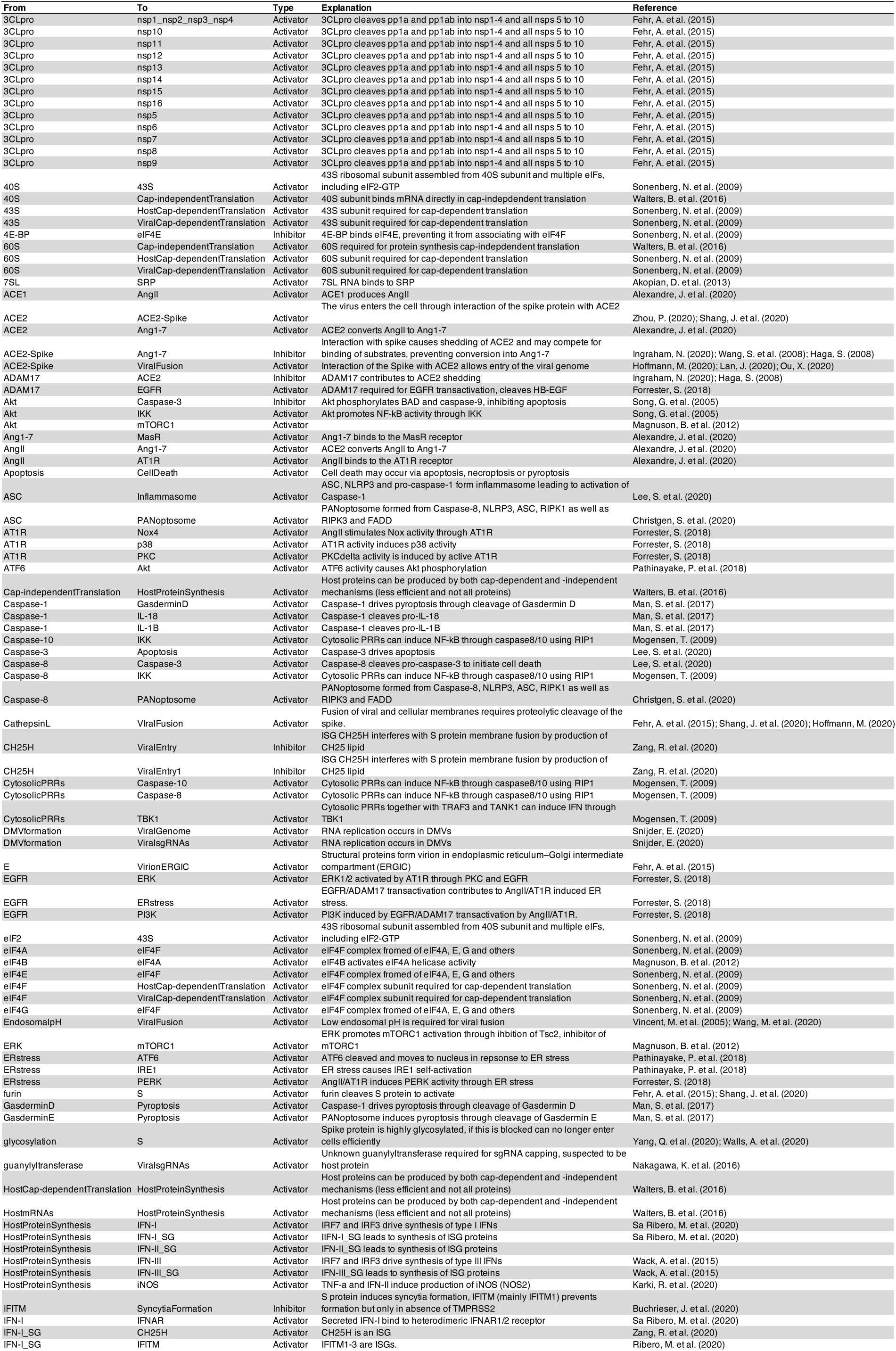
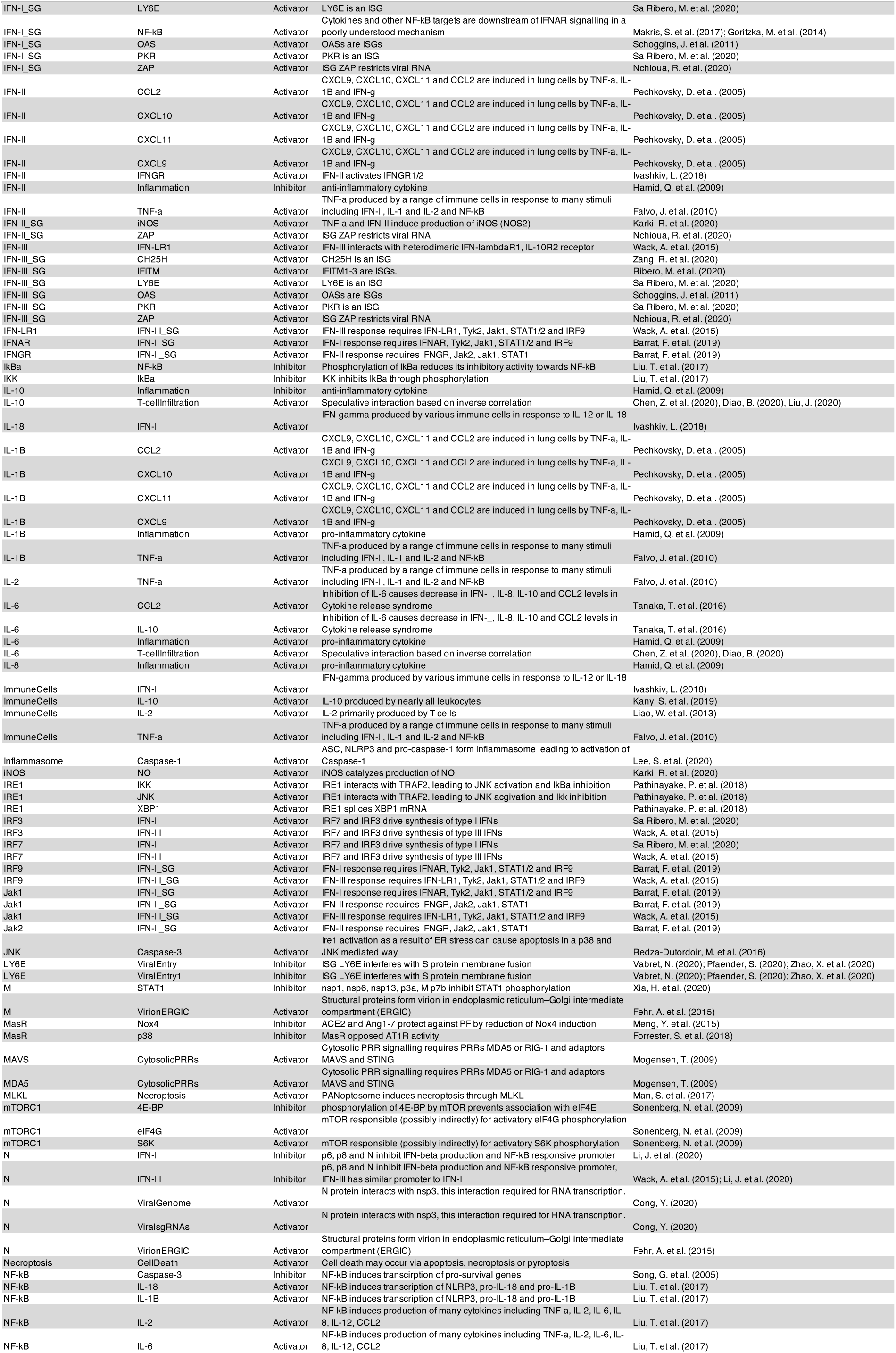
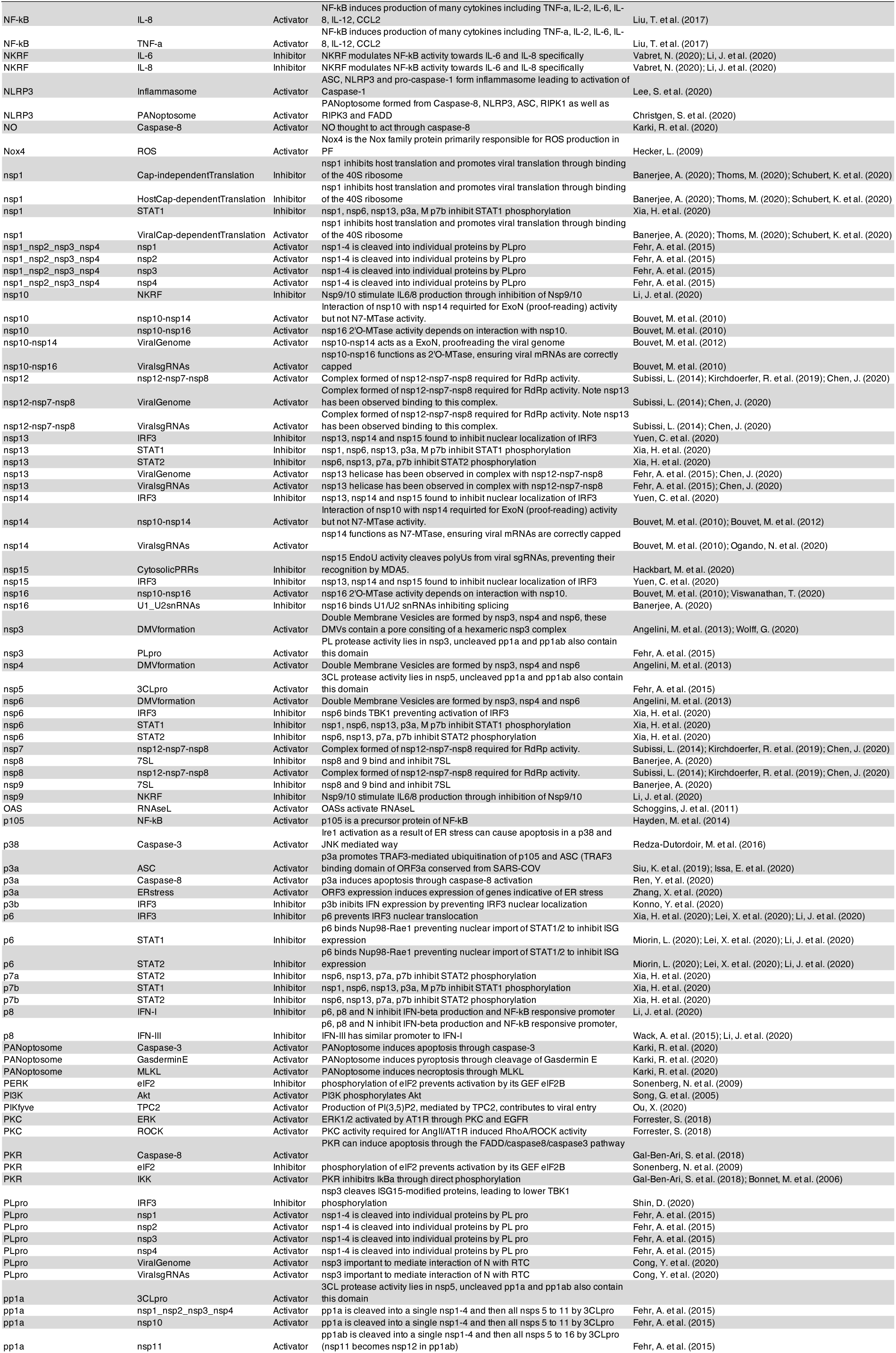
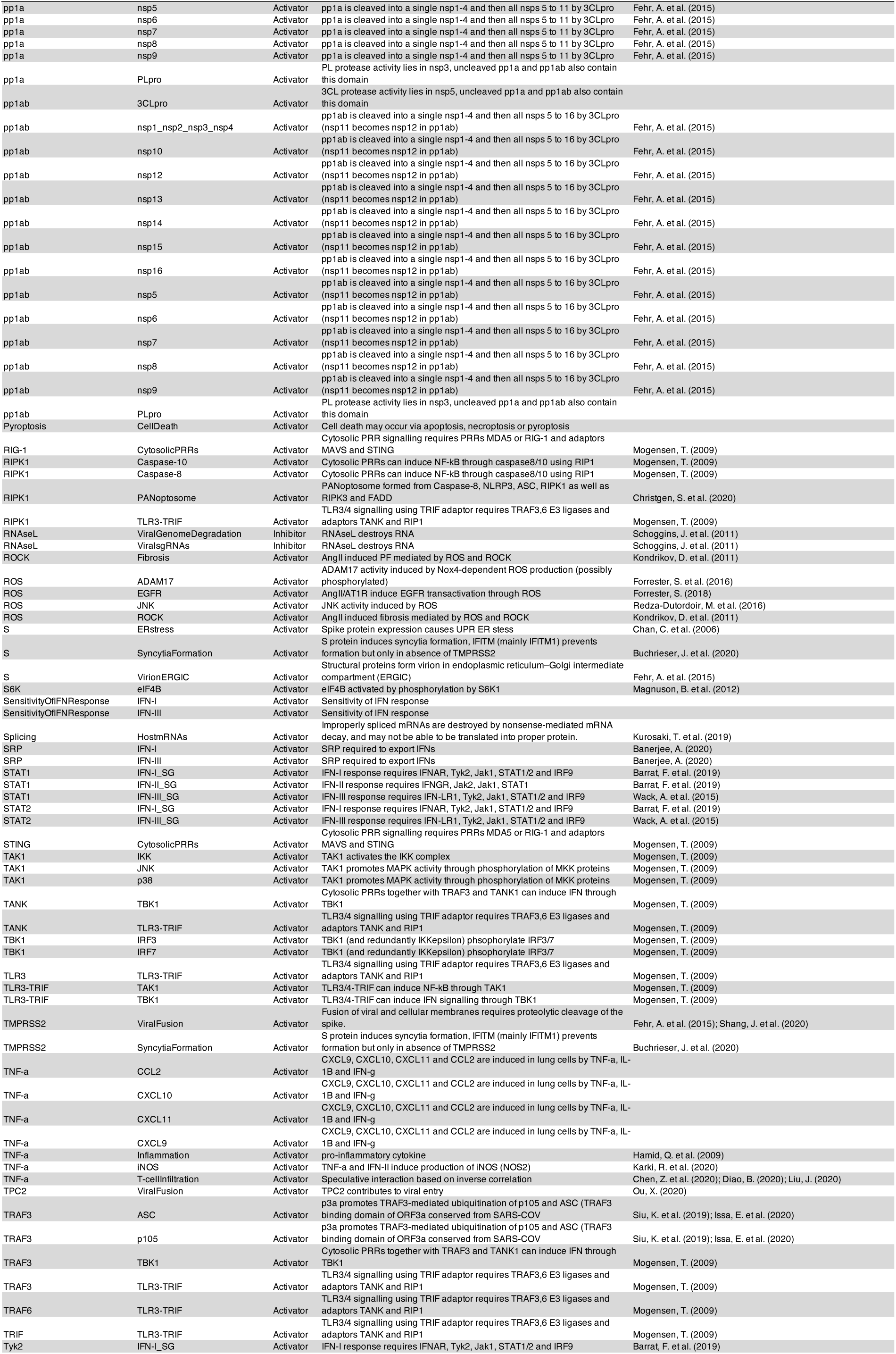
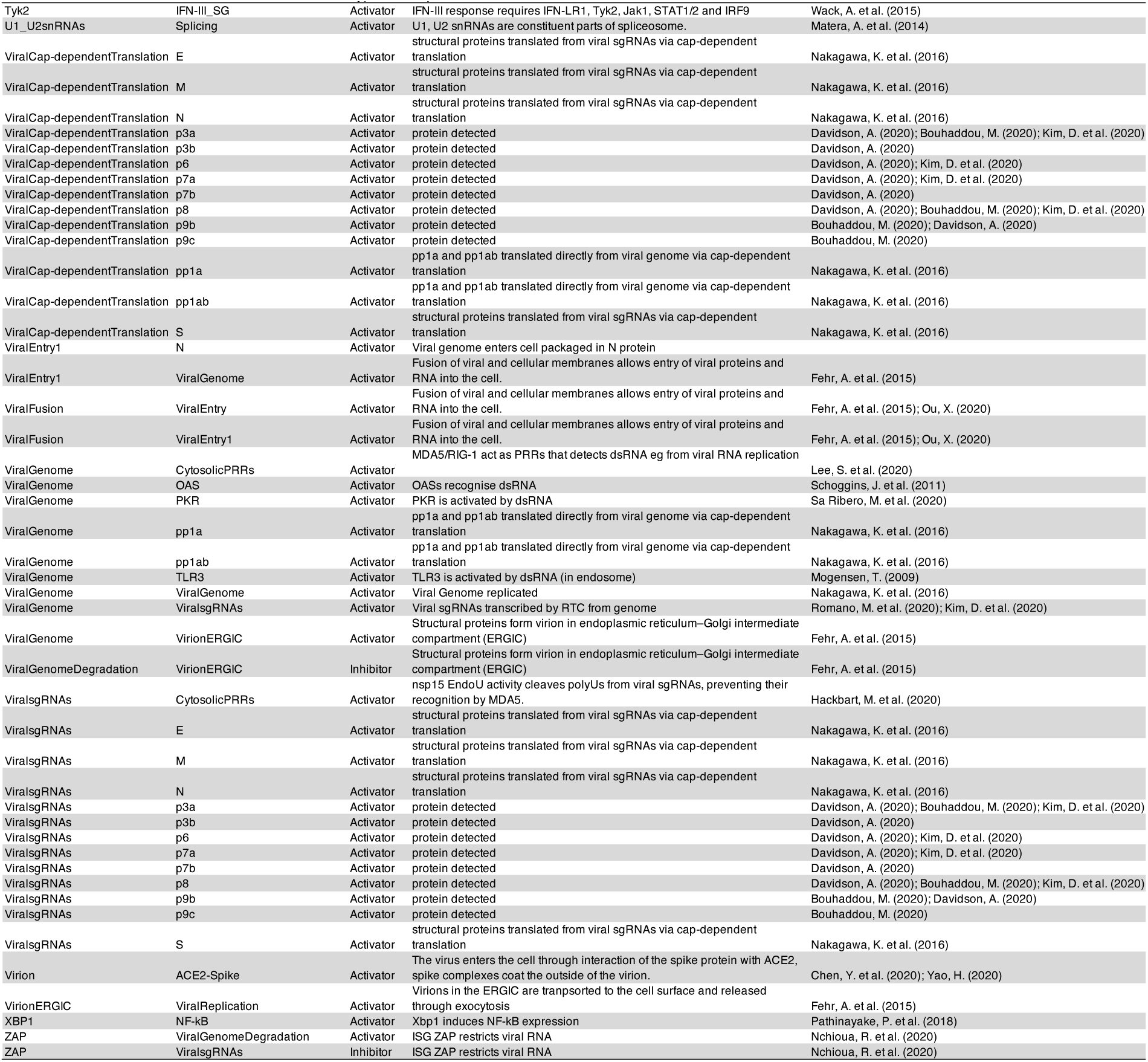
Regulatory edges in the model of SARS-CoV-2 infection. Each row shows a single edge, originating from the node in the “from” column and directed to the node in the “to” column. The table also contains details of the type of edge, a brief explanation of the mechanism of this regulation and a reference to the literature where required.

**Table S2.**
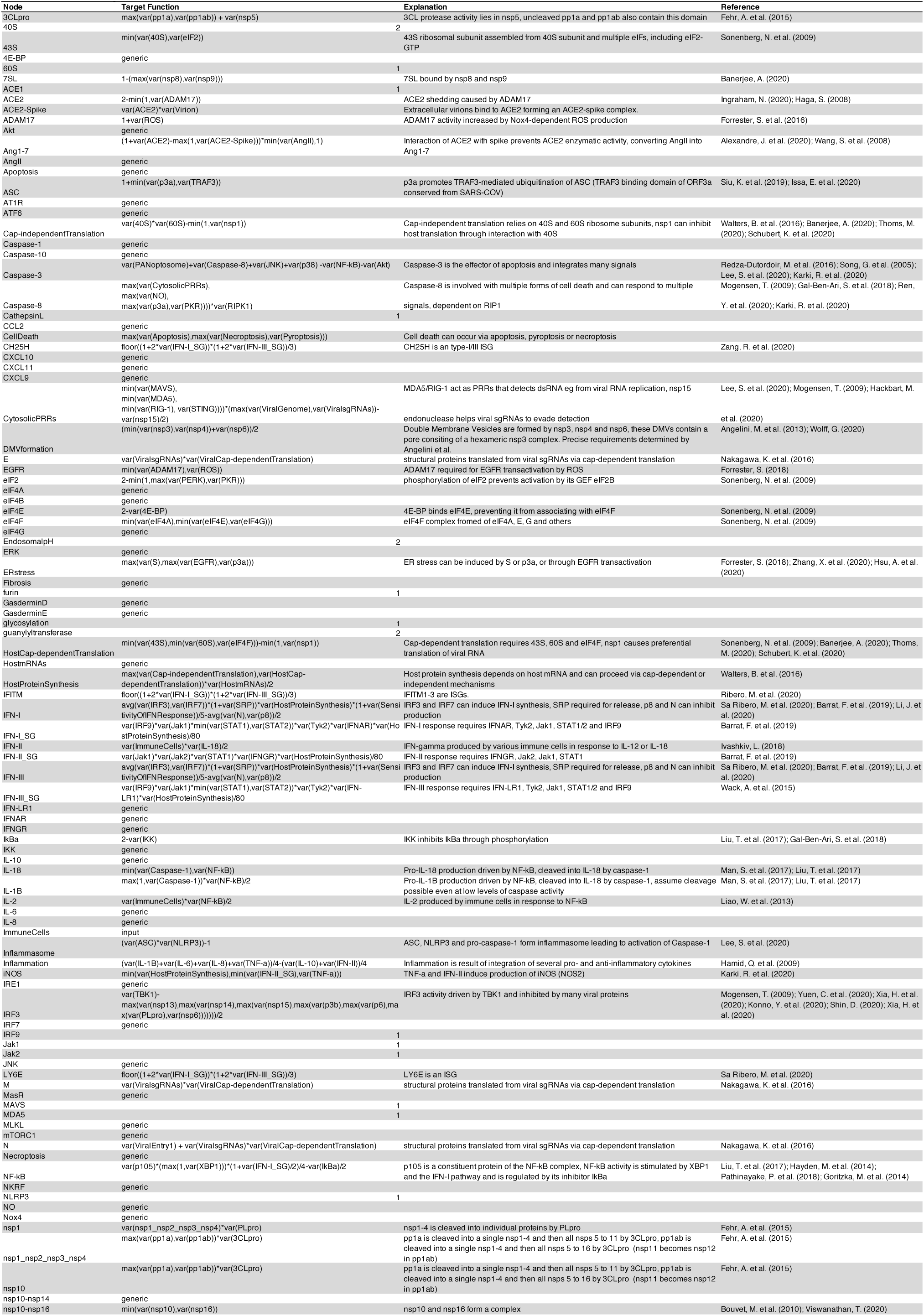
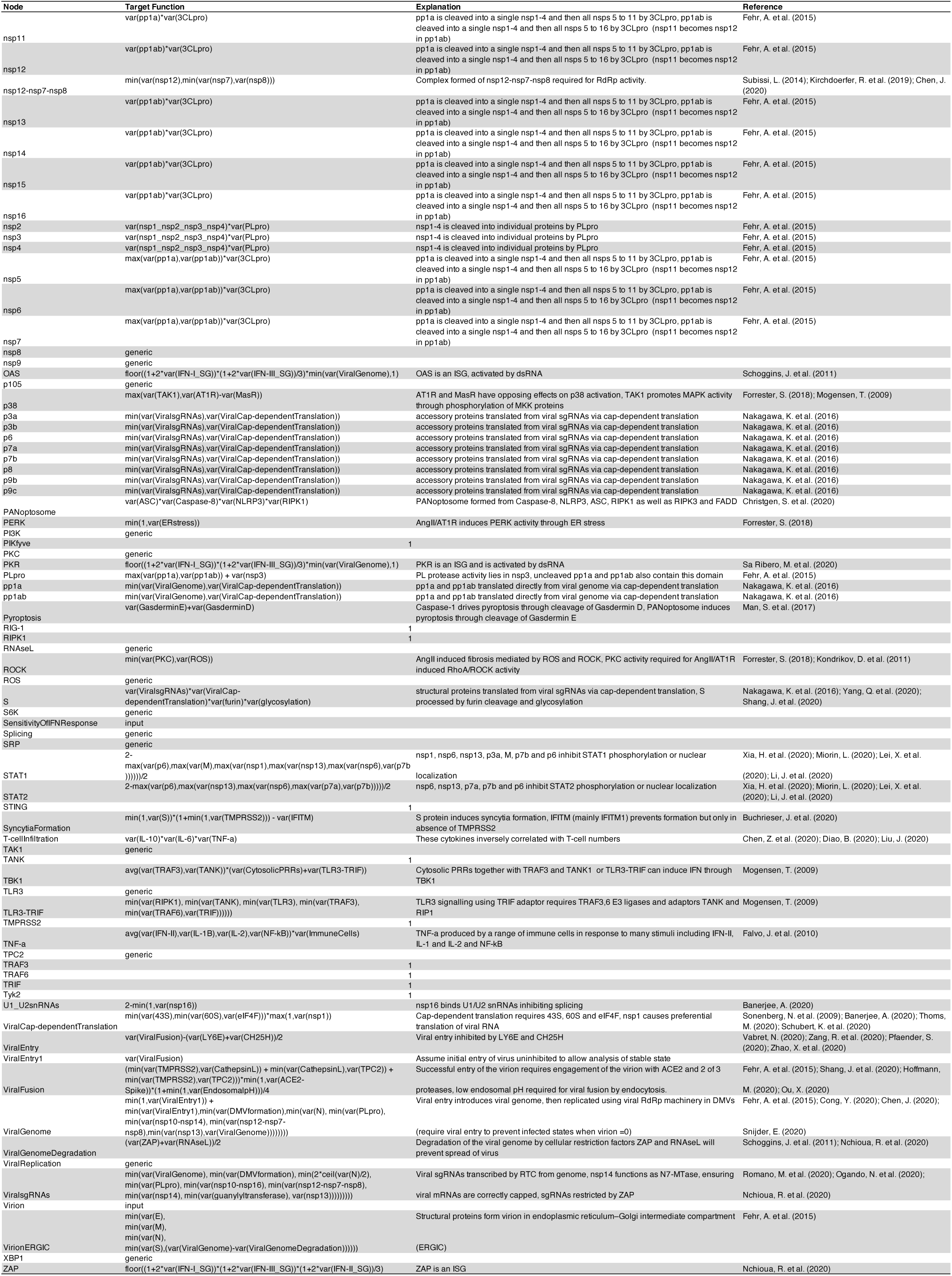
Table of target functions used in the model. The target function for each node is specified, along with a brief description of the mechanism and a reference where required.

**Table S3.** Constraints applied for each state of the model. For each model state, the constrained nodes and their value are given.

| Background | Node | Activity |
| --- | --- | --- |
| Uninfected | Virion | 0 |
|  | ImmuneCells | 1 |
|  | Sensitivity of IFN Response | 0 |
| Mild COVID-19 | Virion | 1 |
|  | ImmuneCells | 1 |
|  | Sensitivity of IFN Response | 1 |
| Early Stage, Severe COVID-19 | Virion | 1 |
|  | ImmuneCells | 1 |
|  | Sensitivity of IFN Response | 0 |
| Late Stage, Severe COVID-19 | Virion | 1 |
|  | ImmuneCells | 1 |
|  | Sensitivity of IFN Response | 2 |
|  | ViralGenome | 2 |

**Table S4.**
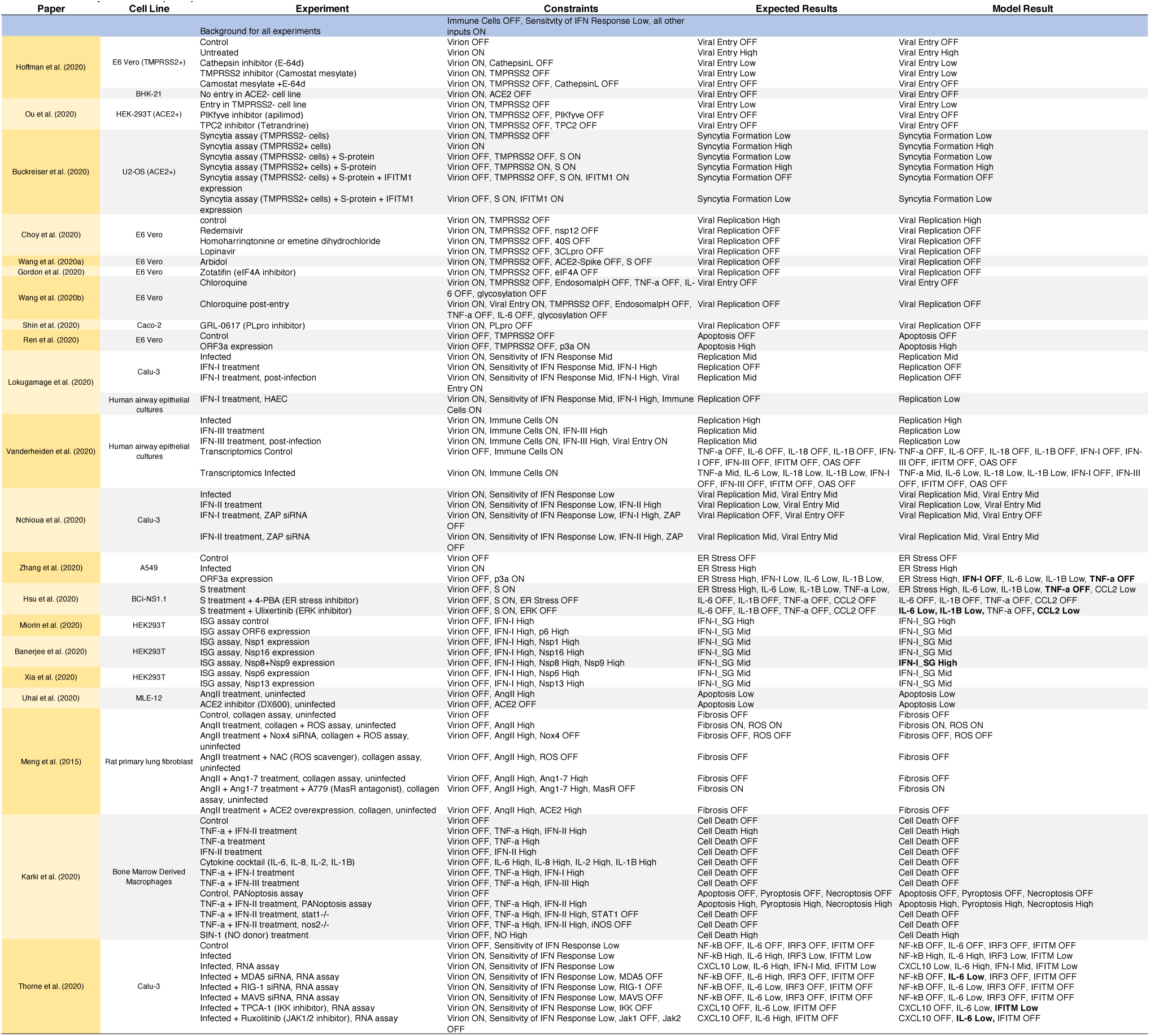
List of specification (tests) to validate the model. **In vitro experiments used to validate the model of SARS-CoV-2 infection.** For each experiment, the table shows the publication or pre-print, the cell line used, the constraints applied to the model and the expected outcome, based on the experimental result. The final column shows the result of the model simulations, with deviations from the expected results highlighted in bold.

**Table S5.**
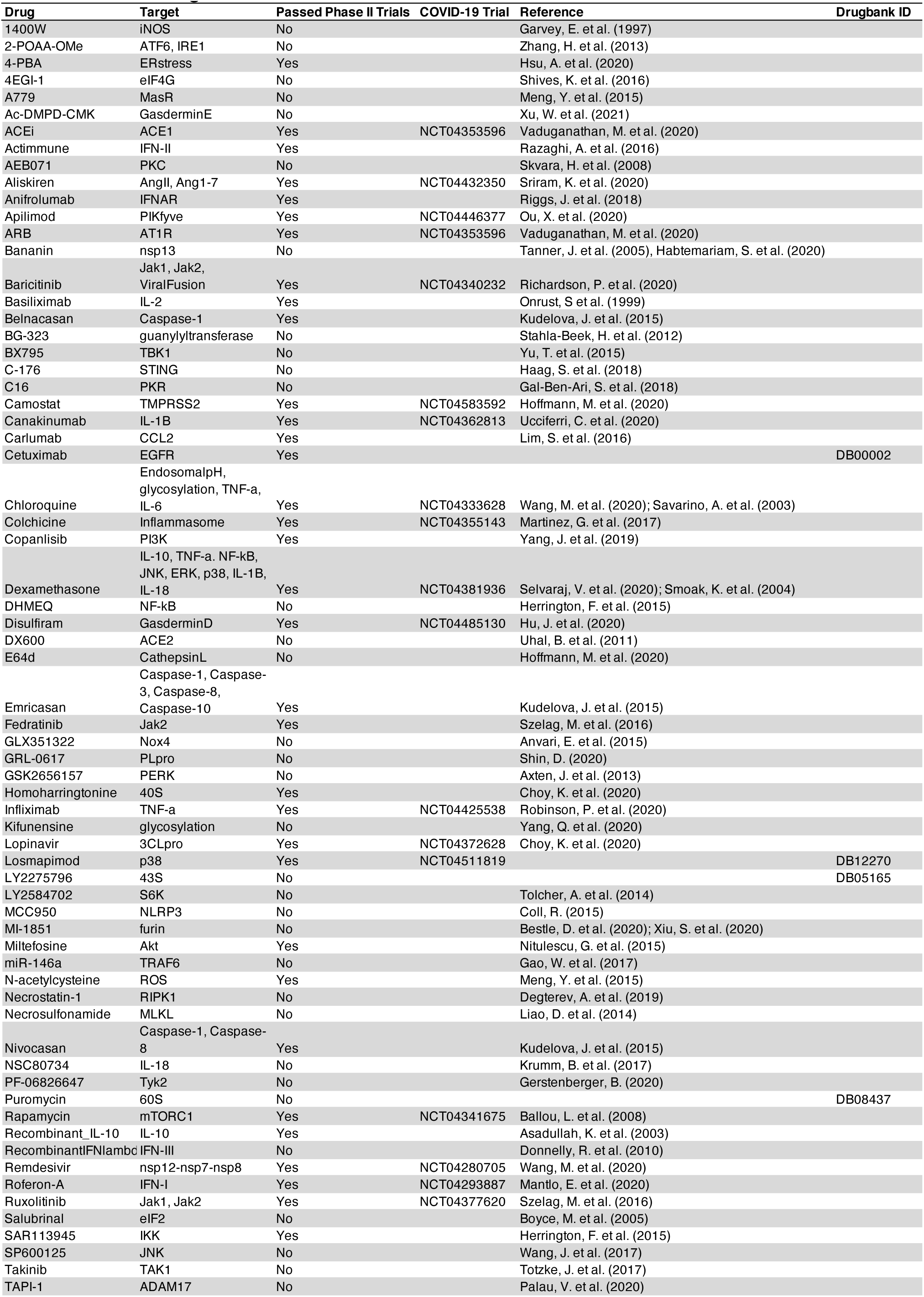
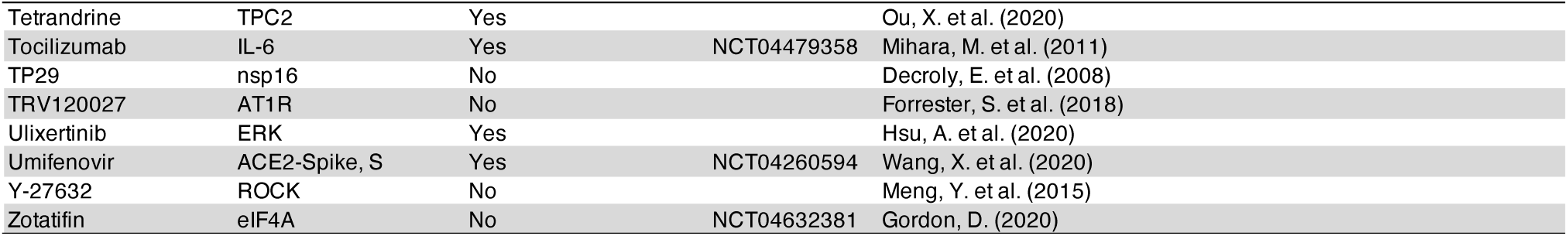
Table of drugs used for *in silico* drug repurposing screens. For each drug, the table contains its targets, whether it has passed phase II clinical trials (1 means it has), clinical trials identifier (if in trials to treat COVID-19), and a reference or drugbank ID.

**Table S6.**
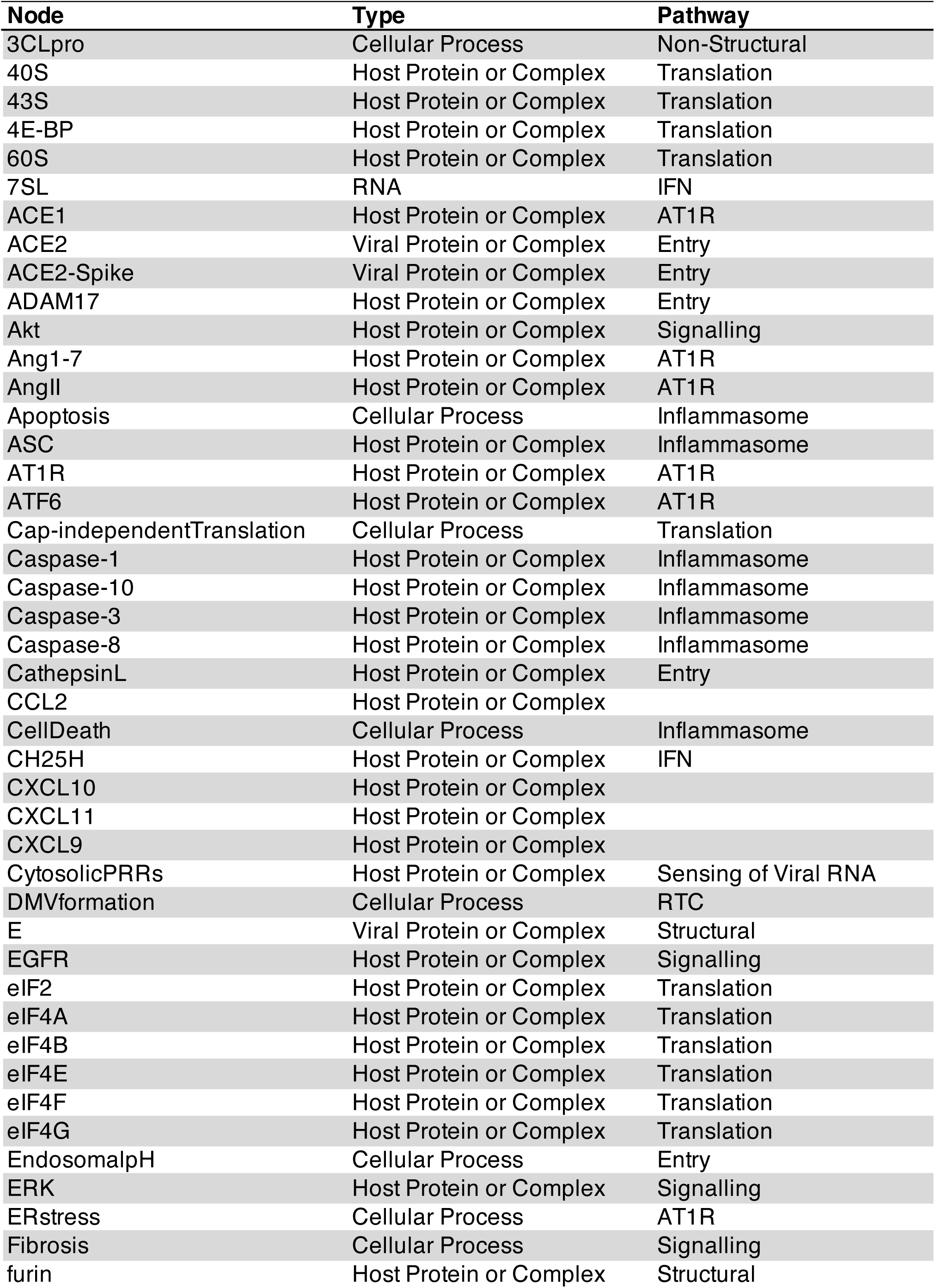
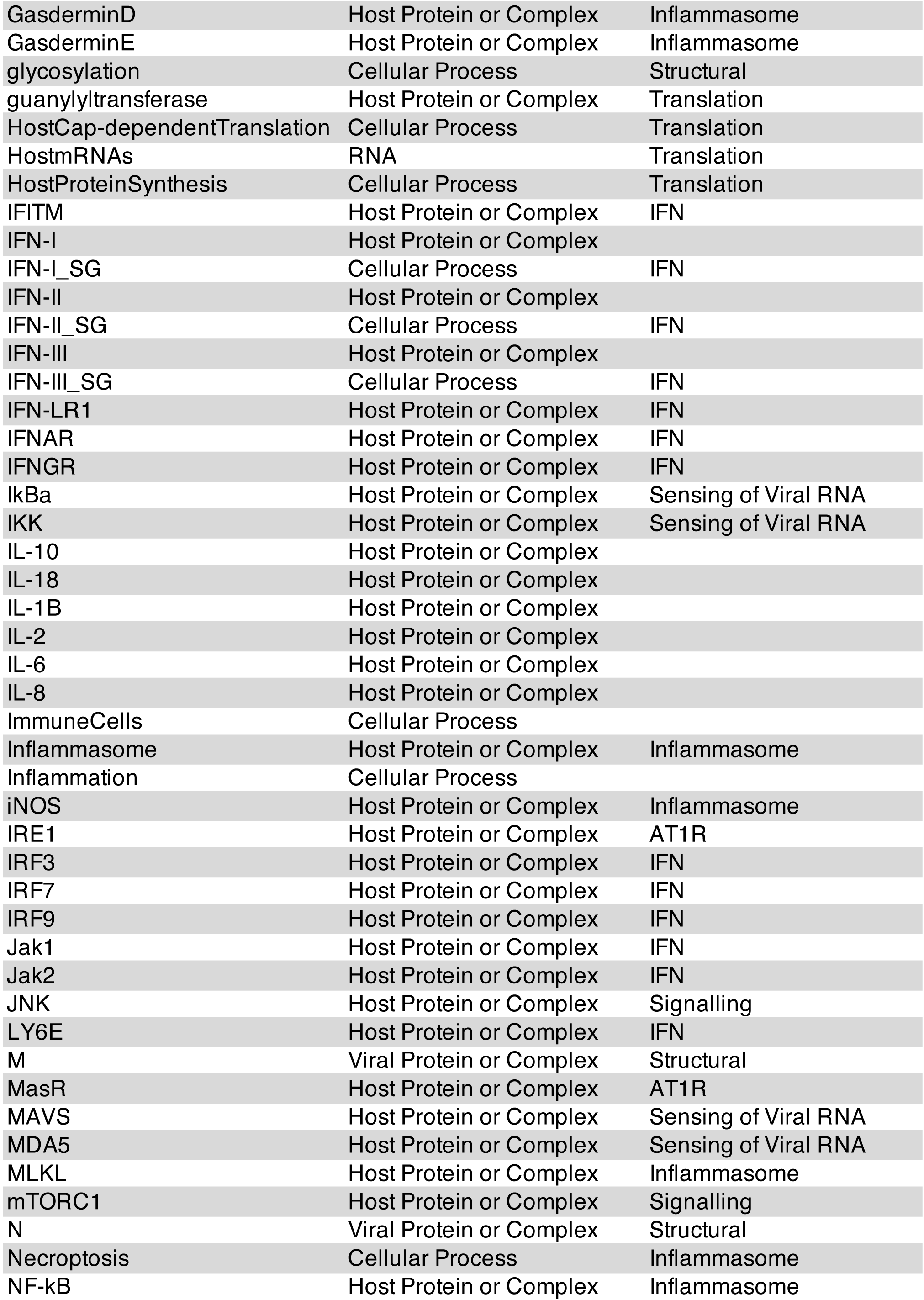
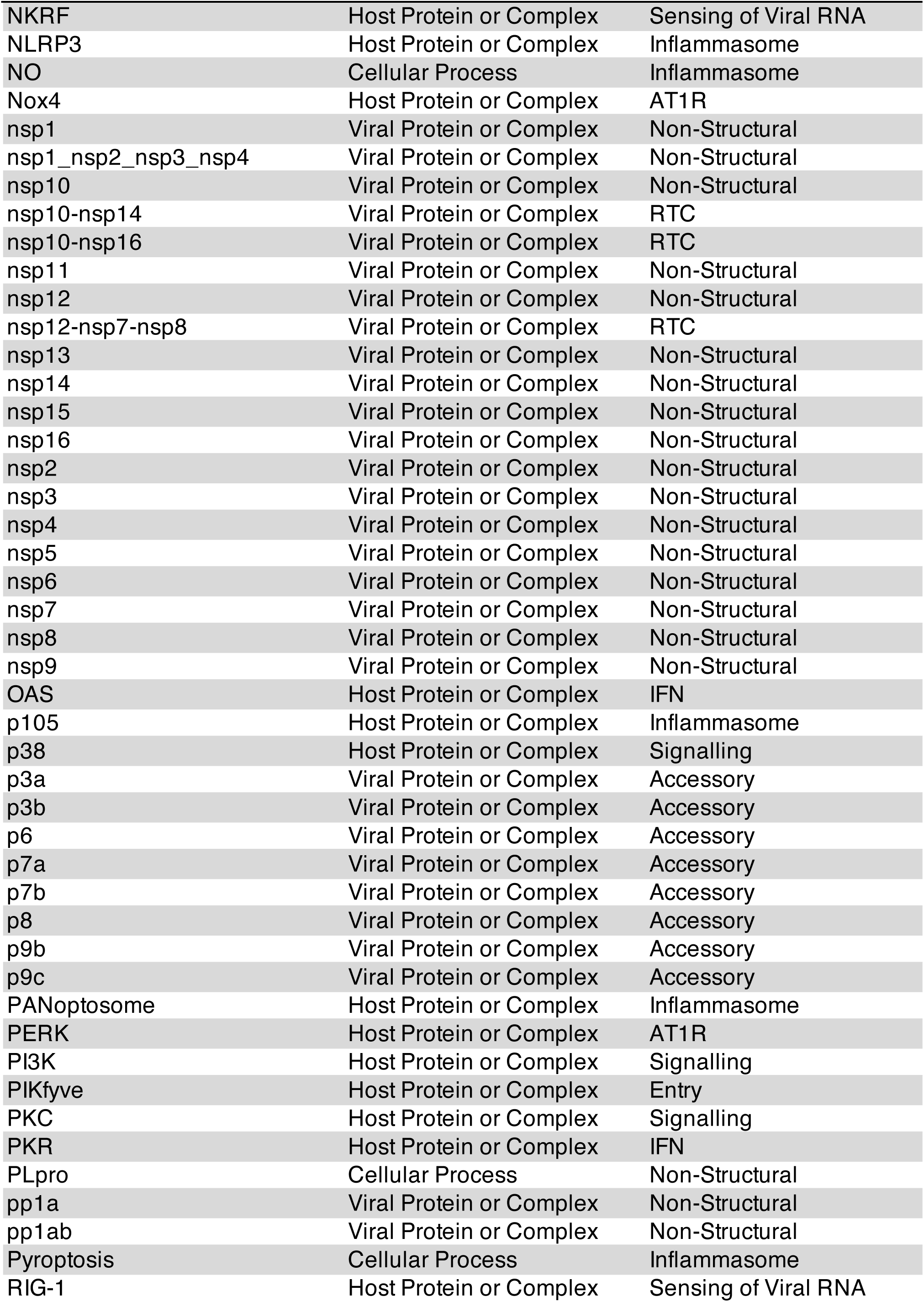
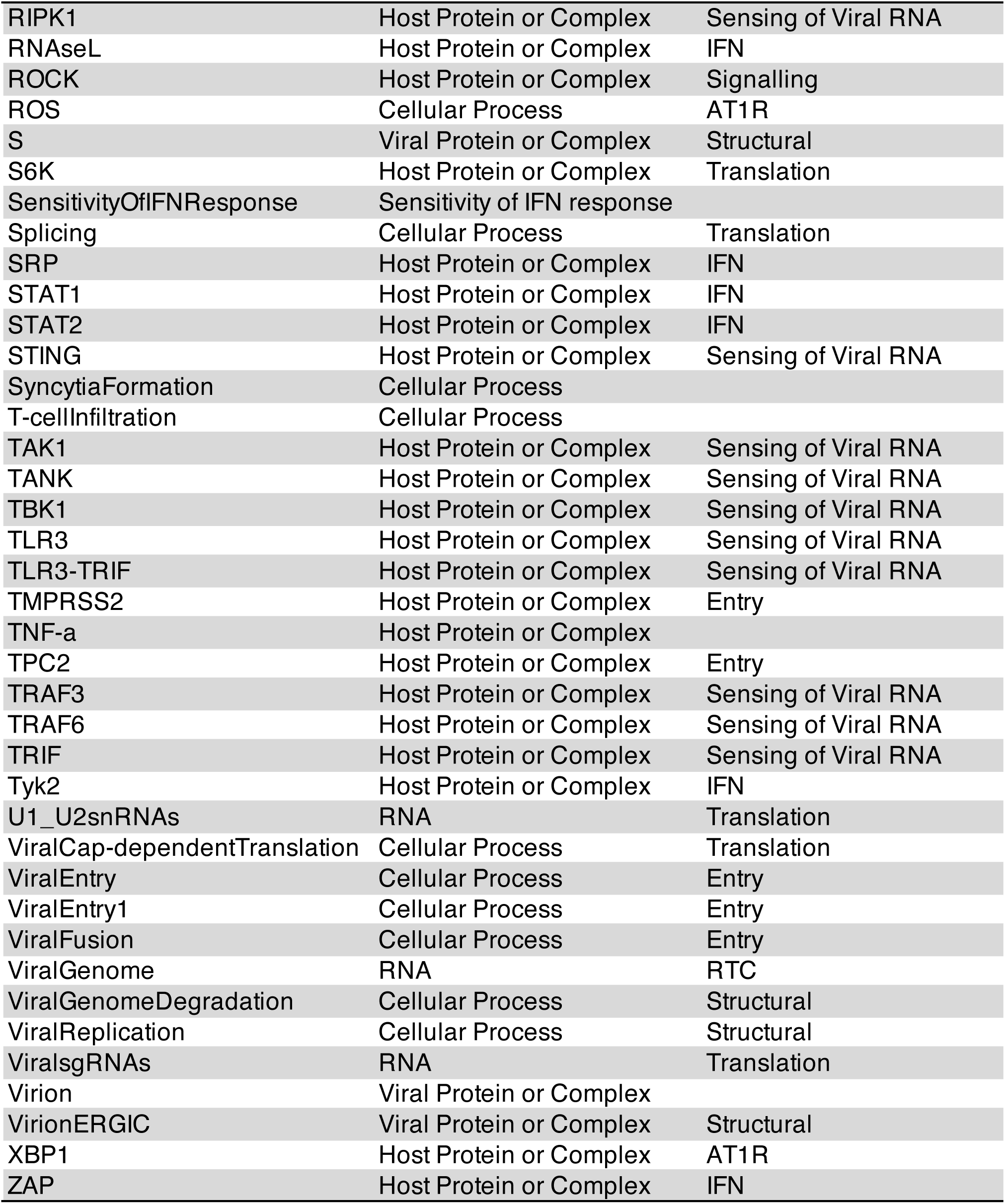
Categories of nodes in the model. For each node, the type and pathway within the model are specified.

**Table S7.** *In silico* simulation of Caco-2 cell drug treatments. The Caco-2 cell line is modelled *in silico* by setting the “ImmuneCells” and “SensitivityOfIFNResponse” nodes to zero, reflecting the fact that Caco-2 cultures consist of clonal colorectal epithelial cells and do not mount an IFN response to SARS-CoV-2 infection.

| Condition | Treatment | Inflammation | Viral Entry | Viral Replication |
| --- | --- | --- | --- | --- |
| Caco-2 | Control | 1 | 4 | 4 |
| Caco-2 | Apilimod | 1 | 2 | 4 |
| Caco-2 | Camostat | 1 | 2 | 4 |
| Caco-2 | Camostat + Apilimod | 0 | 0 | 0 |
| Caco-2 | Miltefosine | 1 | 4 | 4 |
| Caco-2 | Dexamethasone | 0 | 4 | 4 |
| Caco-2 | Dexamethasone + Miltefosine | 0 | 0 | 0 |

